# Dynamic matrices with DNA-encoded viscoelasticity for advanced cell and organoid culture

**DOI:** 10.1101/2022.10.08.510936

**Authors:** Y.-H. Peng, S. K. Hsiao, K. Gupta, A. Ruland, G. K. Auernhammer, M. F. Maitz, S. Boye, J. Lattner, C. Gerri, A. Honigmann, C. Werner, E. Krieg

## Abstract

3D cell and organoid cultures, which allow in vitro studies of organogenesis and carcinogenesis, rely on the mechanical support of viscoelastic matrices. However, commonly used matrix materials lack rational design and control over key cell-instructive properties. Herein, we report a class of fully synthetic hydrogels based on novel DNA libraries that self-assemble with ultra-high molecular weight polymers, forming a dynamic DNA-crosslinked matrix (DyNAtrix). DyNAtrix enables, for the first time, computationally predictable, systematic, and independent control over critical viscoelasticity parameters by merely changing DNA sequence information without affecting the compositional features of the system. This approach enables: (1) thermodynamic and kinetic control over network formation; (2) adjustable heat-activation for the homogeneous embedding of mammalian cells; and (3) dynamic tuning of stress relaxation times over several orders of magnitude, recapitulating the mechanical characteristics of living tissues. DyNAtrix is self-healing, printable, exhibits high stability, cyto-and hemocompatibility, and controllable degradation. DyNAtrix-based 3D cultures of human mesenchymal stromal cells, pluripotent stem cells, canine kidney cysts, and human placental organoids exhibit high viability (on par or superior to reference matrices), proliferation, and morphogenesis over several days to weeks. DyNAtrix thus represents a programmable and versatile precision matrix, paving the way for advanced approaches to biomechanics, biophysics, and tissue engineering.

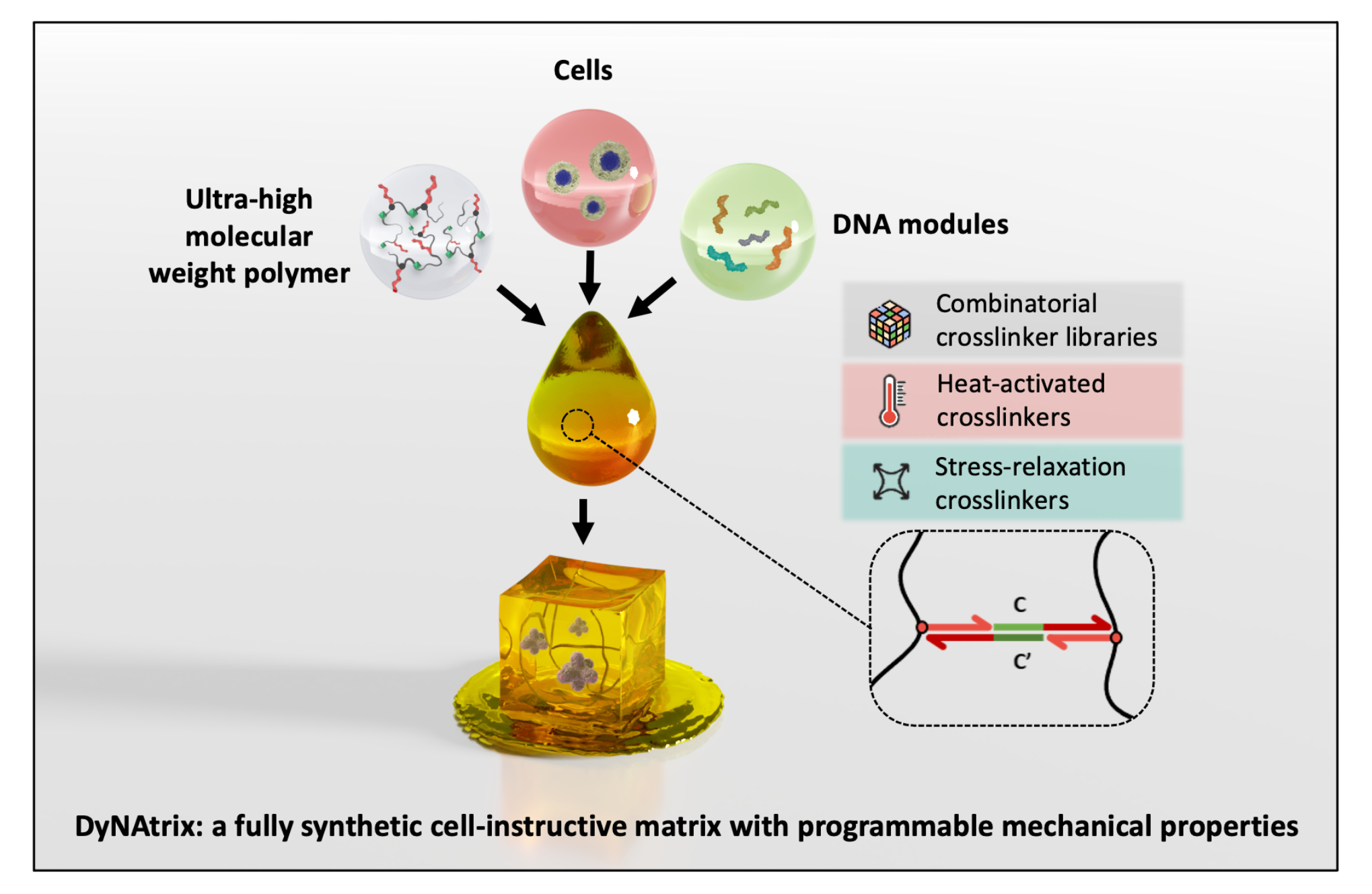

## Introduction

Supramolecular hydrogels are water-swollen three-dimensional (3D) networks of molecules, linked together by non-covalent bonds^1, 2^. The dynamic nature of these bonds gives access to unique properties, such as stimuli-responsiveness and self-healing^3, 4^. Hydrogels that contain self-assembling DNA are particularly interesting, as the *sequence-selective* recognition of complementary DNA strands enables modular construction and de-construction of complex functional materials^5–7^. With the help of DNA nanotechnology, the properties of these systems can be adjusted and dynamically changed *in situ*^8, 9^. DNA- based materials can thus be adapted to respond to biomolecules and cells to sense^10–13^, actuate^14, 15^, and release^16, 17^. Amongst their many applications, DNA-based hydrogels could support 3D cell cultures, for instance, to mechanistically study cell and developmental biology, recapitulate pathologies, and develop (personalized) therapies^17–20^.

The (micro)environment of cells in the extracellular matrix (ECM) is one of the key determinants for cell fate decisions^21^. In particular, the stiffness of ECM and ECM-mimicking materials can affect cell proliferation and morphology in 3D organoid cultures via mechanosensitive transcriptional regulators^22, 23^. Beyond stiffness, it has recently been recognized that matrix viscoelasticity entails other critical parameters^24, 25^. Specifically, the *stress relaxation* behavior describes the time-dependent change of mechanical stress in the material following a deformation. Faster stress relaxation can promote cell spreading^26^, migration^27^, and differentiation^28^. However, the most widely used hydrogel systems for 3D cell culture are naturally-derived basement membrane matrices, such as *Matrigel*, which do not allow precise tuning of critical material properties, including stress relaxation. They also exhibit significant batch-to-batch compositional variation and are prone to cause unexpected stimulation of cells^29^. More advanced materials that do offer viscoelastic tunability require significant modifications to the chemical composition of the system, whose influence on cells is fundamentally challenging to disentangle from mechanical effects. An ideal cell-instructive substitute material should be chemically defined and adapt its mechanical properties in a programmable manner^21, 30^. Moreover, it should be printable under mild cytocompatible conditions to enable top-down fabrication of complex 3D tissue templates^31–33^.

Synthetic DNA-crosslinked hydrogels appear uniquely suitable for addressing the need for a highly adaptive tissue culture platform. However, DNA oligonucleotides are typically required at very high concentrations (5 to 50 g/L). The high cost of oligonucleotide synthesis often makes these materials prohibitively expensive to be deployed at scale. Furthermore, large DNA concentrations can also induce undesired effects when in contact with living cells and tissues. For instance, DNA can electrostatically scavenge positively charged biomolecules^34^ or stimulate inflammatory response pathways^35, 36^. Thus, there has been significant interest in either reducing the cost of DNA material synthesis^37^ or supporting the network with secondary non-DNA crosslinks^14^.

We herein report a novel approach to a programmable dynamic DNA-crosslinked matrix platform (DyNAtrix). DyNAtrix is based on a diverse set of DNA modules that self-assemble with a biocompatible DNA-grafted ultra-high molecular weight (UHMW) poly(acrylamide-co-acrylic acid) backbone. By using novel crosslinker libraries—rather than single crosslinker splints—the formation of the supramolecular network and its mechanics can be uniquely controlled without changing crosslinker concentrations or other chemical components of the system. As a result, gelation occurs at very low DNA concentrations, where crosslink efficiencies approach the theoretical limit for affine molecular networks. Importantly, the DNA libraries provide dynamic control over viscoelasticity and plasticity, tunable crosslinking kinetics and mixing behavior, as well as adjustable degradability. These features demonstrate the unique programmability of soft materials powered by DNA nanotechnology as 3D cell culture matrices and printable bio-inks.

## Results

### Material concept and design

To create a programmable yet inexpensive and cytocompatible hydrogel, we first synthesized three derivatives of a UHMW poly(acrylamide-co-acrylic acid)-graft-DNA^38, 39^ copolymer (Figure 1a). We chose this backbone due to its high molecular weight, anticipated biocompatibility, and methanol responsiveness. The latter enables efficient removal of unreacted DNA and cytotoxic acrylamide monomers. We hypothesized that the UHMW backbone could serve as the majority structural component, while a relatively small amount of DNA would suffice to induce network formation and dictate material properties (Supplementary Note 3.1). The polymer is named **P_x_**, where **x** indicates the relative abundance of conjugated DNA strands (Figure 1a). We first synthesized the three derivatives **P_1_**, **P_5_**, and **P_10_** (Supplementary Method 2.1) and characterized them by asymmetrical flow field-flow fractionation with light scattering detection, gel electrophoresis, and nuclear magnetic resonance spectroscopy (Supplementary Figures S1-S3). **P_1_**, **P_5_**, and **P_10_** have a molecular weight of approximately 3 megadalton (MDa) (Supplementary Table S2) and an estimated average number of 3, 20, and 28 DNA strands per backbone, respectively (Supplementary Note 3.2). The covalently attached DNA strands serve as universal *anchor* sites for self-assembly with DNA crosslinkers. Moreover, they allow modular noncovalent extension of the material with other functional DNA constructs. A single polymer batch can thus be used to assemble materials with vastly diverse properties. For studies involving cell culture, additional synthetic peptide side chains were included in the synthesis. The peptides contain the arginylglycylaspartic acid (RGD) motif that facilitates cell adhesion. The two synthesized peptide-grafted polymer derivatives (termed 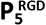 and 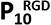) had similar yield and molecular weight as their peptide-free counterparts (Supplementary Tables S2, S3).

**Figure 1.**
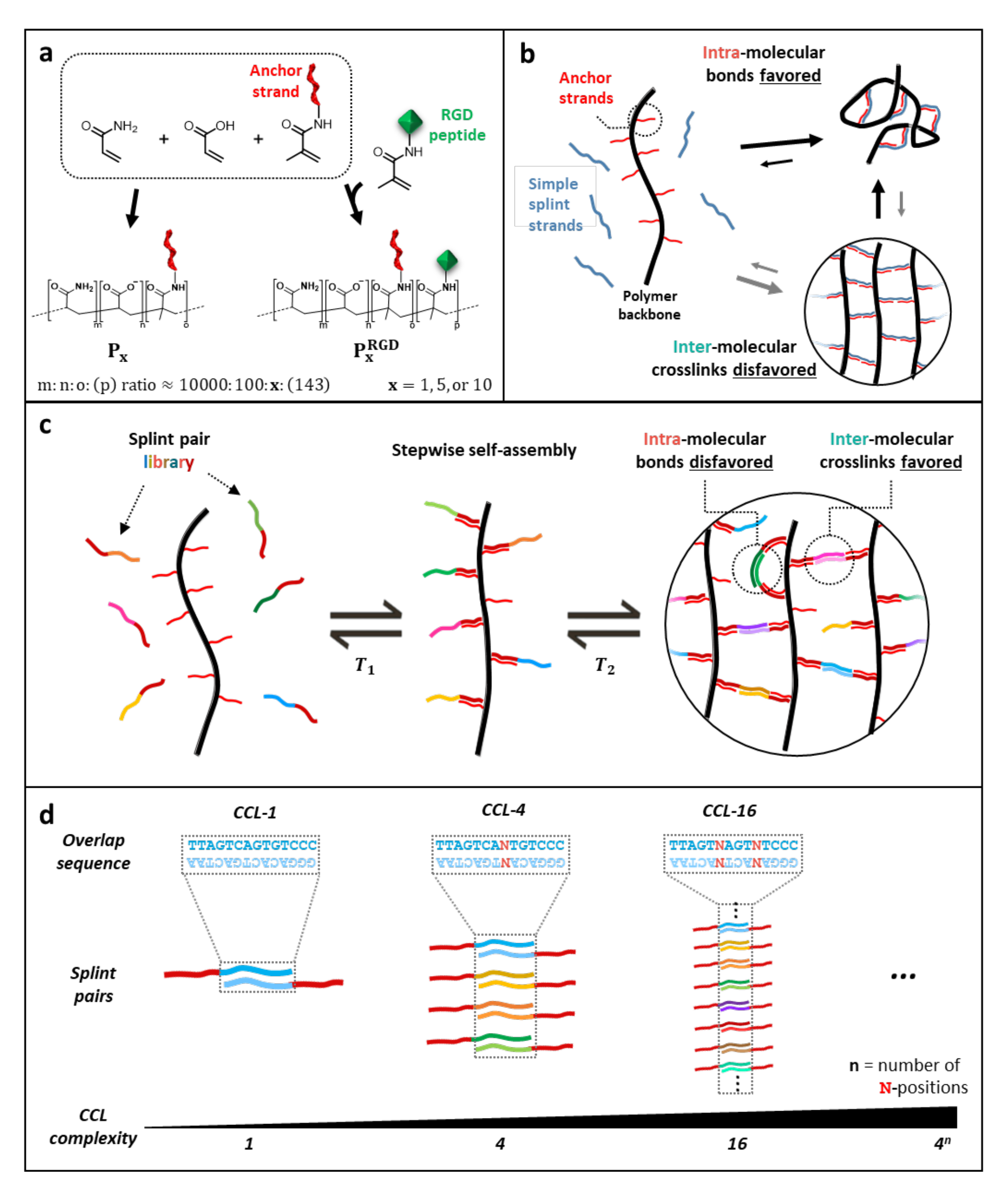
Basic concept and crosslinker design for DyNAtrix. (a) Polymer synthesis via free radical polymerization of acrylamide, sodium acrylate, and acryloyl-based monomers modified with anchor strand DNA (red). For cell culture, acrylamide-labelled cell adhesion peptides with RGD motif (green) were co-polymerized. (b) Scheme of intramolecular crosslinks predominant in high-molecular weight polymers at low concentrations, resulting in low crosslinking efficiency and poor mechanical strength. (c) Scheme of polymer crosslinking via combinatorial crosslinker library (CCL) splints to suppress the formation of intramolecular bonds and promote effective intermolecular crosslinks. (d) CCLs are generated by introducing ambiguous bases (N) in the overlap domain.

To our surprise, initial attempts to crosslink the polymers at low concentration with simple crosslinker splints failed to produce a stable hydrogel. As the mechanical strength of a hydrogel is proportional to the number of effective crosslinks (Supplementary Note 3.3), we hypothesized that polymer self-binding (i.e., ineffective *intra-*molecular bonds) dominated over network-forming (i.e. *inter-*molecular) crosslinks (Figure 1b). The predominance of undesirable intramolecular bonding is not specific to DNA gels, but a general phenomenon in crosslinked polymer gels, typically reducing crosslink efficiency to 20% or less^40, 41^.

The sequence-selectivity of DNA hybridization offered a unique solution to this problem: instead of using a single splint type, we constructed complex libraries based on a dual splint design (Figure 1c,d). All library members have an identical *adapter* domain that binds to the anchor strands below a specific melting temperature, T_1_. Additionally, each splint has an *overlap* domain that is designed to pair with only one other splint below a second melting temperature, T_2_. The melting temperature is predictable with high accuracy based on nearest-neighbor thermodynamic models^42^. We designed anchor-and overlap domains to have sufficiently separated melting temperatures, ensuring that upon slow cooling from 95°C anchor domains would bind prior to overlap domains (Supplementary Figure S4).

In principle, overlap domains can be designed with perfectly orthogonal sequences. However, this would require costly synthesis of many different oligonucleotides. Instead, we chose a combinatorial approach where overlap domains are diversified by introducing ambiguous bases (N) at specific positions (Figure 1d). N indicates a sequence position at which an equal mixture of A, T, C, and G nucleotides is used during oligonucleotide synthesis. The complexity (i.e., the number of distinct splint pairs) of this *combinatorial crosslinker library* (CCL) equals 4^n^, where n is the number of ambiguous bases in the sequence. As the DNA duplex is strongly destabilized even by a single base mismatch^43^, equilibrium binding strongly favors perfectly complementary binding partners. We carried out thermodynamic predictions for all possible splint combinations in libraries ranging from 1 (n=0) to 256 (n=4) splint pairs, confirming that highly selective splint pairing (>95%) is expected at equilibrium in all crosslinker libraries (Figure 2b, Supplementary Figure S5, Supplementary Table S1).

**Figure 2.**
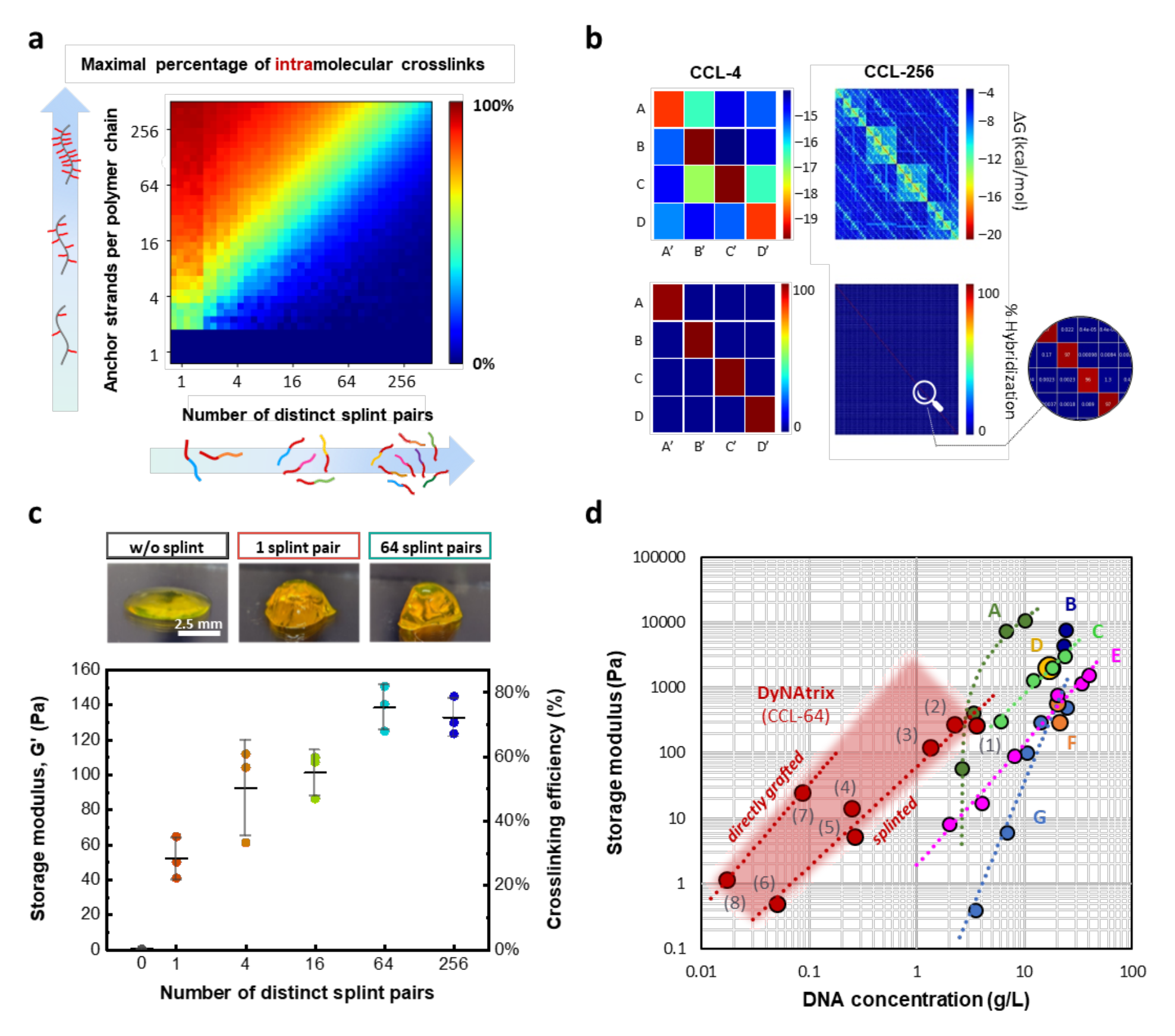
Theoretical prediction and rheological validation of DyNAtrix crosslinked via combinatorial crosslinker libraries (CCL). (a) Statistical simulation of the maximum percentage of intramolecular crosslinks as a function of CCL complexity and the number of anchor strands per polymer backbone. (b) The predicted minimum free energy (top) and the predicted hybridization interactions (normalized Boltzmann distribution, bottom) of crosslinker libraries CCL-4 and CCL-256 at equilibrium. High-resolution matrices for all crosslinker libraries are shown in Supplementary Files 1-10. (c) Top: Photographs of DyNAtrix, stained with SYBR-Gold. Bottom: Average stiffness of DyNAtrix (1% (w/v) **P**_5_) crosslinked via 1, 4, 16, 64, 256-splint library at 20°C. The corresponding crosslinking efficiencies were estimated from the affine network model (Supplementary Note 3.2). All samples have a total DNA content of 1.3 g/L. Error bars represent the standard deviation from three independent experiments. (d) Different DNA hydrogel designs give access to different concentration and stiffness regimes: comparison between previously reported synthetic DNA- crosslinked hydrogels (A: Lin et al. 2004^45^; B: Yue et al. 2019^47^; C: Du et al. 2019^48^; D: Cangialosi et al. 2017^49^; E: Yang et al. 2021^50^; F: Xing et al. 2018^46^; G: Li et al. 2015^19^) and several DyNAtrix gels (this study, red). DyNAtrix samples (1)-(6) were crosslinked with CCL-64 splints, while samples (7) and (8) had CCL-64 directly grafted onto the polymer backbone (cf. Figure 1c, Supplementary Figure S10). (1) 1% (w/v) **P**_10_, (2) 1.65% (w/v) **P**_5_, (3) 1% (w/v) **P**_5_, (4) 1% (w/v) **P**_1_, (5) 0.2% (w/v) **P**_5_, (6) 0.2% (w/v) **P**_1_, (7) 1% (w/v) **P_1_**, (8) 0.2% (w/v) **P_1_**.

### CCL complexity uniquely controls network formation and matrix stiffness, allowing gelation at exceptionally low DNA content

According to statistical simulations, the suppression of undesired intra-molecular crosslinks strongly depends on the complexity of the CCL, as well as the number of anchor strands per polymer backbone (Figure 2a). To suppress at least 80% of intramolecular crosslinks, **P_1_**, **P_5_**, and **P_10_** were predicted to require approximately 4, 40, and 60 splint pairs, respectively. We first studied the mechanical properties of CCL- crosslinked **P_5_** at 1% (w/v) polymer concentration by oscillatory rheology. The polymer was annealed with five CCL versions comprising 1, 4, 16, 64, and 256 splint pairs (*CCL-1* to *CCL-256*), and each sample contained an equal DNA concentration of 1.3 g/L. In all cases, the elastic moduli, G’, significantly exceeded the loss moduli, G’’, over a wide range of frequencies and strains, confirming that the combination of dual splint CCLs with the UHMW polymers generates stable gels (Supplementary Figure S6).

Adjusting CCL complexity represents a novel approach to tune the elasticity of a gel without changing the crosslinker or polymer concentration. Increasing the CCL complexity from CCL-1 to CCL-64 (at constant overall DNA concentration) improved crosslinking efficiency from approximately 28% to 76% (Figure 2c, Supplementary Note 3.1), which is in good qualitative agreement with the prediction (Figure 2a). The CCL complexity thus controlled G’ values in the range of 50 to 140 Pa, which is comparable to *Matrigel*, the most widely used cultural matrix for organoids. We note that our statistical simulation (Figure 2a) only predicts the upper limit for intramolecular bonds, but it does not account for kinetic or certain topological effects.^44^ For instance, increasing library complexity from 64 to 256 splint pairs did not further increase stiffness. We suspect that the marginal benefit of using *CCL-256* is likely offset by its slower binding kinetics, since overlap domains of polymer-bound crosslinker strands become increasingly unlikely to find and capture their zero-mismatch binding partners in very large libraries. We also note that CCLs are expected to improve crosslinking efficiency only when the solid content of the gel is low. This is because at high concentrations and under equilibrium conditions intermolecular bonds can outcompete intramolecular loops. The latter can also mechanically interlock, thereby contributing to the network’s elasticity^44^.

The high crosslinking efficiency and low mass fraction of DNA in the polymer chains enable the formation of stable gels at extraordinary low DNA content, up to two orders of magnitude lower than previously reported synthetic DNA-crosslinked hydrogels with well-characterized rheological properties^14, 19, 45–48^ (Figure 2d). *CCL-64* crosslinked **P_1_** and **P_5_** gels showed critical gelation concentrations (CGC) of approximately 2 g/L (corresponding to a total DNA content of 0.05 and 0.26 g/L, respectively). The CGC is close to the polymers’ apparent density, ρ_app_, indicating that gelation occurs efficiently as soon as polymer blobs start overlapping in solution (ρ_app_ ∼ 1.3 g/L for **P_1_** and **P_5_**; Supplementary Table S2). Even lower DNA contents (down to 0.017 g/L) were achieved when—instead of anchor strands—the *CCL-64* overlap domain was directly grafted onto the polymer backbone (Supplementary Figure S10; Figure 2d, samples (7) & (8)). The directly grafted crosslinker libraries thus offered a more efficient use of synthetic DNA, however, at the expense of “hard-coding” the crosslinking properties into the polymer. Further experiments were performed with crosslinker splints, as they enable rapid modular adaptation of the material properties without time-consuming polymer synthesis.

Due to the temperature-responsive melting of DNA, all CCL-crosslinked gels undergo a reversible sol-gel transition (Figure 3a, Supplementary Figure S8a). We observed that the melting point of DyNAtrix gradually decreases with increasing CCL complexity from 65°C to 53°C. This observation is expected: as the *variety* of different crosslinker splints increases, the *concentration* of each distinct splint pair decreases (since the total DNA concentration remains constant), thereby reducing the melting point. The observed shift of the gel melting is in perfect agreement with the predicted overlap melting from thermodynamic calculations (Supplementary Figure S8b).

**Figure 3.**
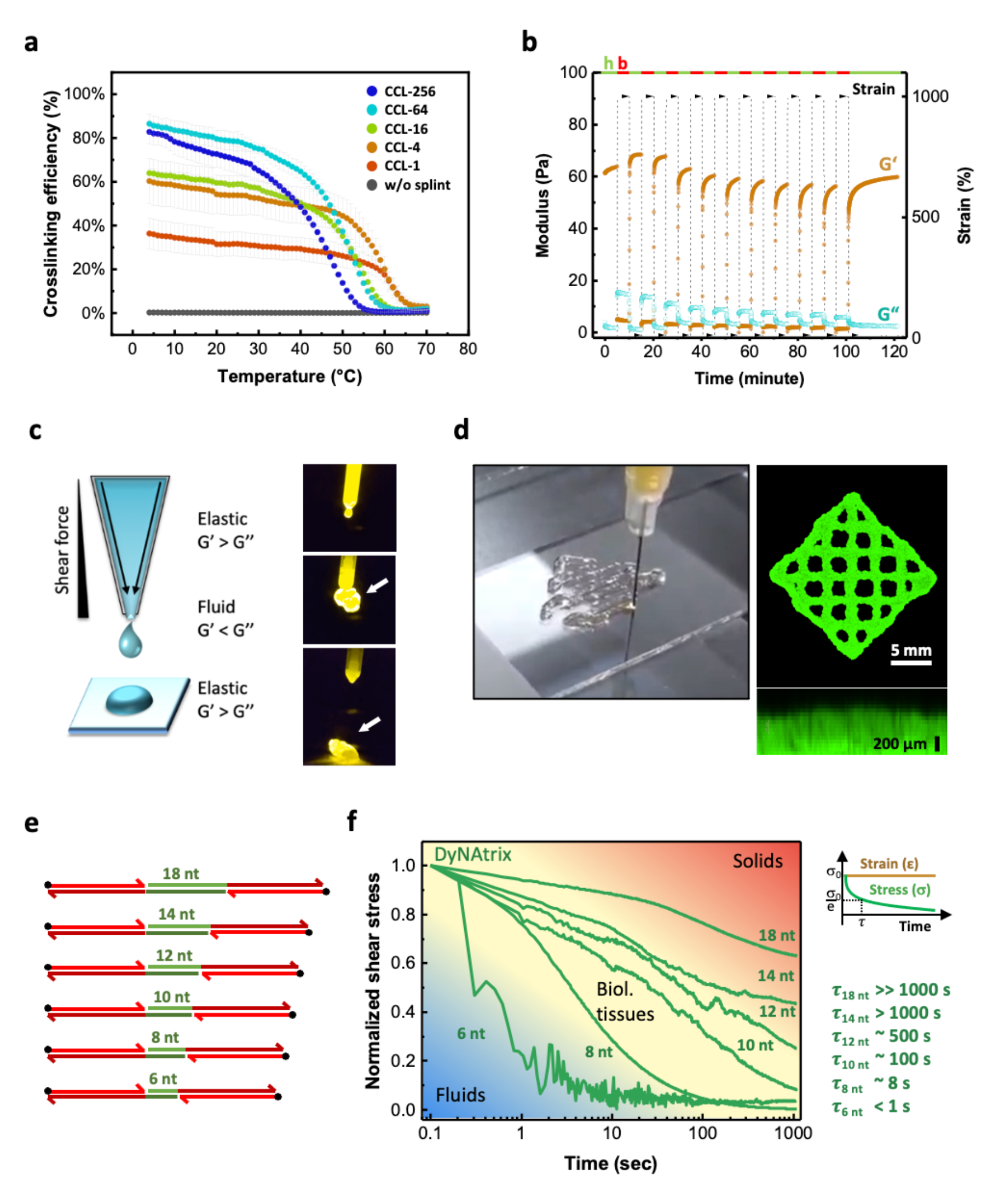
Thermoreversibility, self-healing, bioprinting, and tunable stress-relaxation of DyNAtrix. (a) Crosslinking efficiency of DyNAtrix (1% (w/v) **P_5_** with different CCLs) as a function of temperature. Crosslinking efficiencies were calculated from the affine network model (Supplementary Note 3.3, Supplementary Figure S8). Error bars represent the standard deviation from three independent experiments. (b) Self-healing of DyNAtrix (1% (w/v) **P_5_** crosslinked with CCL-4) was confirmed by repetitive strain recovery tests alternating between 1000% (breaking period, b) and 10% strain (healing period, h). The gel showed over 95% of recovery after 10 consecutive breakage-recovery cycles. (c) Scheme of extrusion printing (left) and photographs of DyNAtrix being extruded from a narrow tip (right, cf. Supplementary Video 1). The gels were prepared with 1% (w/v) **P_5_** crosslinked with CCL-64. (d) A grid printed on a BioScaffolder with a total height of ∼500 μm. The material was stained with SYBR-Gold for imaging purposes. (e) Scheme of stress-relaxation crosslinkers (SRCs) with variable overlap domains (green). The corresponding DNA sequences are shown in Supplementary Table S1. (f) Normalized stress relaxation curves of DyNAtrix crosslinked via different SRCs (green traces) covering the typical range of biological tissues25 (yellow background). Stress relaxation times (τ) were determined by the simple Maxwell model, where σ(τ) =σ_0_/e. The measurements were carried out at 37 °C with 15% strain.

### Rapid self-healing allows printing of complex patterns under cytocompatible conditions

Extrusion bioprinters require self-healing hydrogels that can be easily liquefied under mechanical forces and rapidly recover their viscoelasticity. Such properties allow the material to pass through the nozzle as a fluid, protect the cells from damaging forces, and quickly reform the network after extrusion. To explore the compatibility of DyNAtrix for extrusion bioprinting, we first tested its self-healing properties by oscillatory rheology (Figure 3b, Supplementary Figure S9). When a large strain (1000%) is applied, the supramolecular network breaks apart and transitions into a liquid state, as revealed by a drastic drop of its stiffness (G’ << G”) and increase of the phase angle from 3°±1° to 74°±1°. When reducing the strain, the gel state is quickly restored. After being subjected to 10 consecutive breakage cycles, the storage modulus recovered to over 95% of its original value, revealing that breakage occurs predominantly at the reversible DNA crosslinks rather than the covalent polymer backbone. Due to this rapid self-healing, DyNAtrix quickly solidifies when extruded through a positive displacement pipette (Figure 3c, Supplementary Video 1). This feature allowed extrusion printing of various patterns on a commercial *BioScaffolder* under mild cell-compatible conditions (Figure 3d). DyNAtrix-embedded cells showed high viability and proliferated after printing (Supplementary Figure S14), giving access to the application as *bio-ink* for fabricating complex two and three-dimensional tissue structures.

### Nanomechanical crosslinker stability encodes macroscopic stress-relaxation behavior

As demonstrated by Mooney, Chaudhuri and colleagues, matrix plasticity influences cell development.^28, 51^ Previously described approaches to alter plasticity require significant changes to the chemical composition or temperature—parameters that can influence cell development even without stress-relaxation effects. DNA crosslinkers can uniquely address this challenge, as the sequence and length of DNA duplexes dictate their stability under nanomechanical shear forces. For short DNA (<20nt), the force required to break the duplex into two separate strands grows linearly with the number of complementary bases^52^. Exploiting this phenomenon, we developed stress-relaxation crosslinkers (SRCs) by systematically changing only a few bases in the overlap domain sequence (Figure 3e, Supplementary Table S1). As the overlap domains are the weakest connection points in the supramolecular network, their rupture force was predicted to dictate the material’s stress relaxation behavior. Indeed, SRCs with overlap domains ranging between 6 to 18 nucleotides allowed the adjustment of the stress-relaxation time (τ) from less than 1 s to more than 1000 s. Importantly, the stress relaxation time should not influence the plateau stiffness of the material, as long as the lifetime of the overlap duplex is long compared to the timescale of the experiment. Frequency-dependent rheological measurements confirm that—despite the vastly different stress relaxation times—all SCR-crosslinked gels using 8-18nt overlap domains exhibit similar plateau stiffnesses (Supplementary Figure S7). Due to its sub-second relaxation time, gels crosslinked by 6-nt SRCs are only solid at high deformation frequencies. Overall, SRCs allow precise mimicking of the typical stress relaxation times that are found across diverse animal tissues^28^ (Figure 3f).

### Heat-activated crosslinkers enable homogeneous encapsulation of mammalian cells

The previous sections described how CCLs and SRCs permit predictable engineering of network formation, thermodynamic stability, and (nano)mechanical properties. We next exploited the tunable binding kinetics of DNA in order to make CCL-crosslinked DyNAtrix compatible with existing cell encapsulation workflows. In commercially available cell culture matrices, cells are typically first dispersed within a liquid precursor solution. Subsequently, gelation is induced to encapsulate the cells. For instance, *Matrigel* precursor solution is typically mixed with cells at 4°C, followed by incubation at ∼37°C. The heating step conveniently triggers gelation at the ideal temperature for mammalian cell culture. In contrast, DNA-crosslinked hydrogels typically require heating above the crosslinker melting temperature to become liquid (>50°C), which is incompatible with cell culture. As an alternative to thermal annealing, we first attempted to encapsulate cells by rapidly mixing two pre-annealed polymer precursor solutions (Figure 4b, left). However, the fast hybridization kinetics of DNA leads to crosslink formation before a homogeneous mixture can be achieved. The resulting supramolecular networks were highly inhomogeneous (Figure 4d, left) and exhibited poor mechanical stiffness (Figure 4c, green trace).

**Figure 4.**
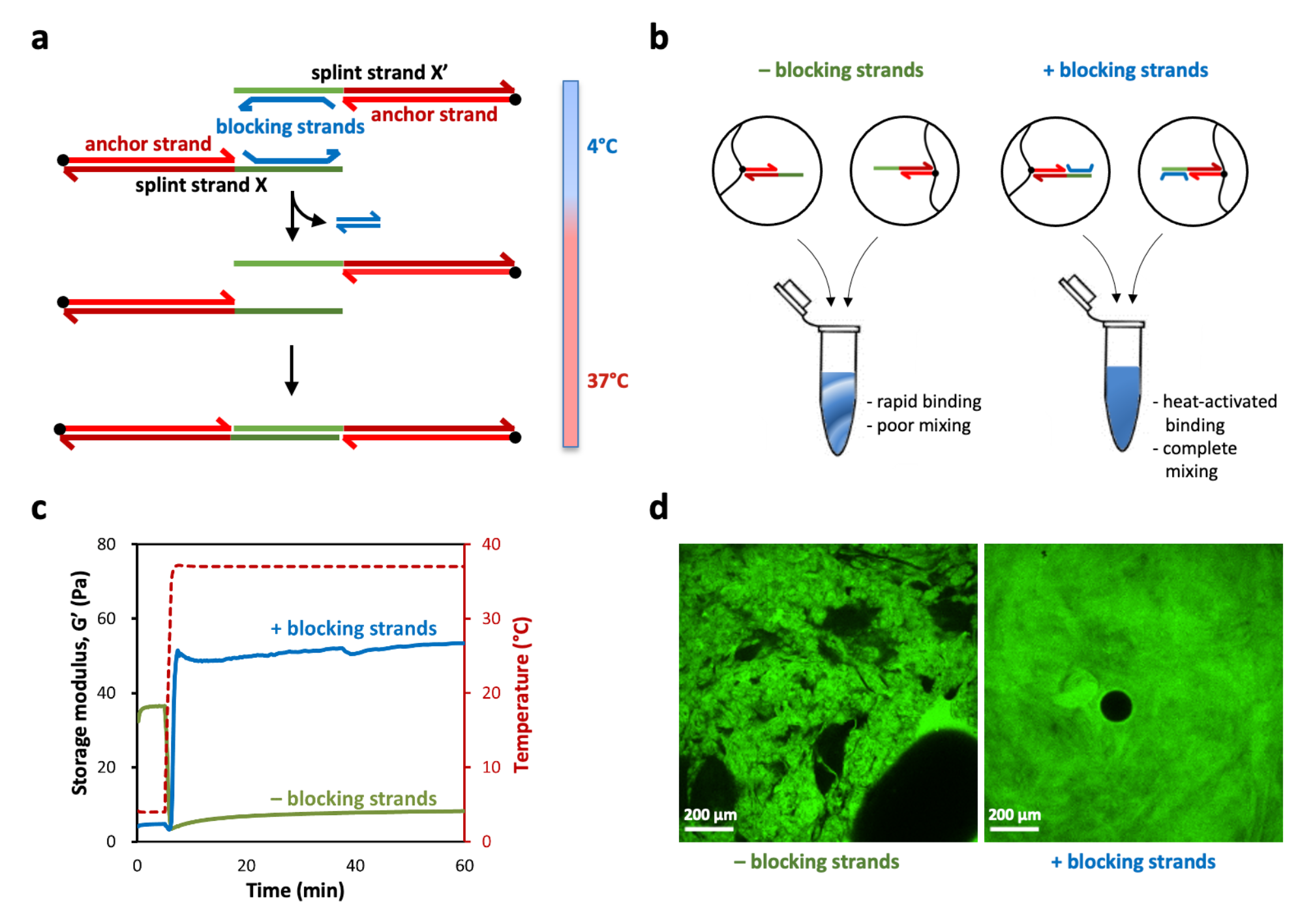
Heat-activated crosslinkers (HACs) enable controlled gelation of DyNAtrix under cell-compatible conditions. (a) Scheme of heat-activated crosslinkers (HACs) and their binding mechanism. (b) Scheme of liquid precursor mixing with vs. without blocking strands. (c) Storage modulus G’ of the gels (1% (w/v) **P_5_** + CCL-64), directly after mixing, with (blue) and without (green) blocking strands. (d) Confocal microscopy images (cross section) of DyNAtrix (1% (w/v) **P_5_** + CCL-64) formed with and without blocking strands. One polymer component was stained with a fluorescein-labeled DNA strand (green fluorescence), while the other polymer component was unlabeled. Left: poor mixing is typically observed in the absence of blocking strands. Right: Highly homogeneous molecular networks are observed with HACs (+blocking strands). The black circle is an air bubble.

As a solution, we aimed to mimic the convenient thermoresponsive profile of Matrigel by altering the DNA crosslinking kinetics. We designed heat-activated crosslinkers (HACs) by upgrading the CCL design with blocking strands (Figure 4a, Supplementary Note 3.4). The blocking strands serve as protection groups that bind to the overlap domain at low temperatures to prevent their premature crosslinking. The blocking strands were designed to spontaneously dissociate from the overlap domains at 37°C, thereby activating the crosslinker for binding. A mixture of two polymer precursor solutions is thus trapped in an off-equilibrium (metastable) liquid state at 4°C, such that the mixing with cells and medium can proceed to completion. In agreement with this design, heating to 37°C leads to rapid formation of a homogeneous gel (Figure 4d, right) with highly superior mechanical stiffness (Figure 4c, blue trace). Note that in absence of blocking strands the material was relatively stiff at 4°C and drastically softened upon heating to 37°C. This finding suggests that during initial mixing numerous *base-mismatched* crosslinks are rapidly formed, yet they are labile and dissociate upon heating, leaving behind a network with few effective crosslinks (Figure 4c, green trace). Due to the homogeneous matrix formation and superior mechanical properties, HACs (i.e., CCLs with blocking strands) were used in all subsequent cell culture experiments (Figure 5 and Figure 6).

**Figure 5.**
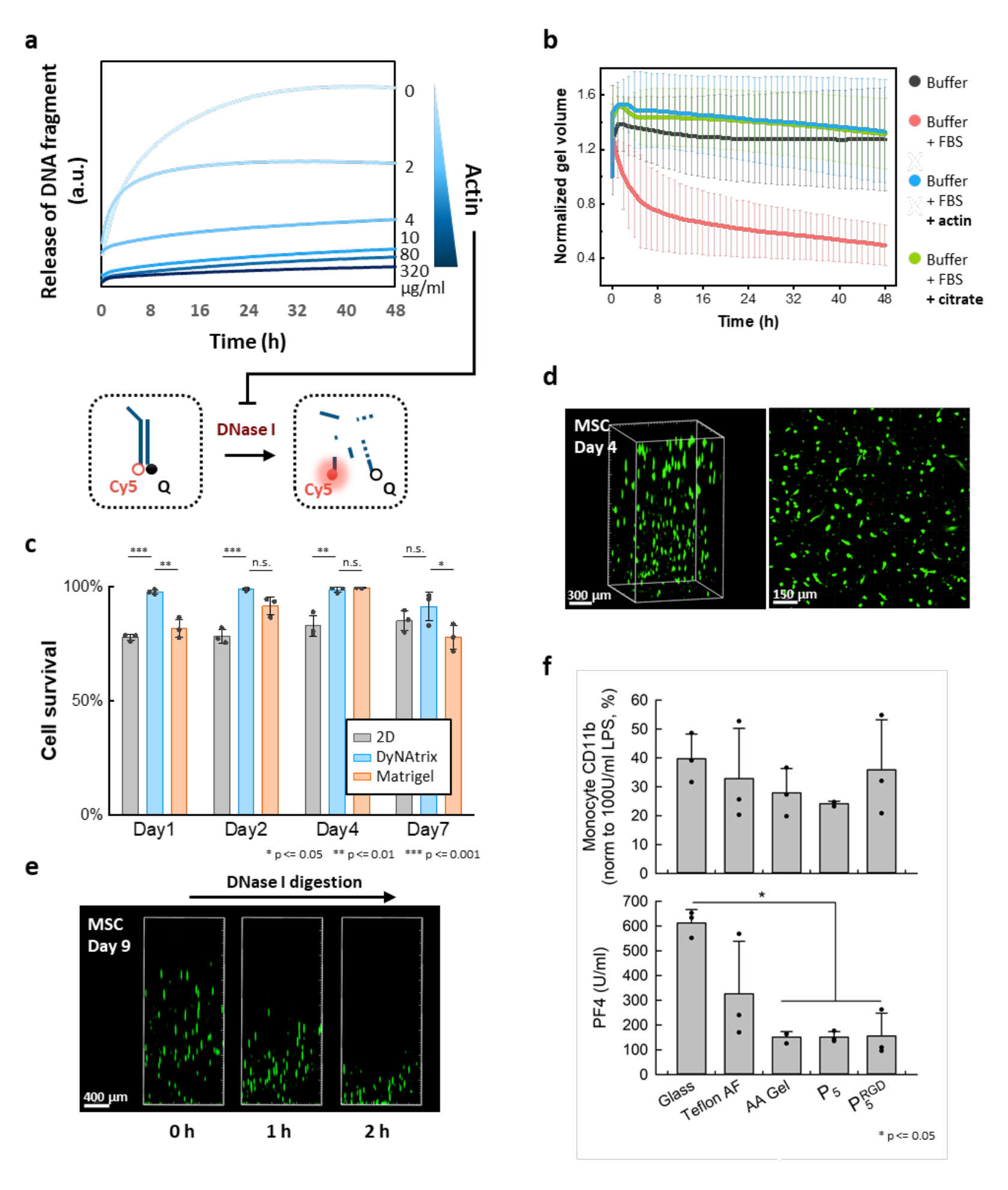
DyNAtrix exhibits tunable stability against degradation in cell culture, high biocompatibility, and DNase-I-mediated release of live cells. (a) Digestion experiment. A digestion probe containing a fluorophore (Cy5) and a fluorescence quencher (Q) was incubated in medium containing DNase-I and different concentrations of actin. Duplex digestion leads to dissociation of the strands, resulting in increased Cy5 emission. (b) Normalized volume of DyNAtrix in FBS-supplemented medium over time. The samples were incubated in buffer (negative control, black) or serum-containing medium without protection (red). The addition of actin (50 µg/mL, blue) or citrate (10 mM, green) effectively protected the gels from digestion. Error bars indicate the standard deviation from 4 independent experiments. (c) Viability of MSCs cultured in actin-protected DyNAtrix compared to Matrigel and a 2D culture plate. Error bars represents the standard deviation of 3 independent experiments. (n.s., not significant; *, p < 0.05; **, p < 0.01; ***, p < 0.001; two-way ANOVA test; Tukey’s multiple comparison test). (d) Live/Dead staining of MSCs in DyNAtrix (1% (w/v) 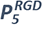 + CCL-64) after 7 days in culture. Live cells were stained with calcein-AM (green) and dead cells were stained with propidium iodide (red). (e) Release of live MSCs from DyNAtrix by addition of DNase I on day 9. (f) Quantification of DyNAtrix-triggered immune response and hemocompatibility. Activation of monocytes in human whole blood (top) was measured by flow cytometric quantification of CD11b expression. Platelet activation (bottom) was assessed by quantification of the hemostasis marker platelet factor 4 (PF4). CCL-64-crosslinked 1% (w/v) **P5** and 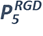 were compared with 3 reference substrates: glass, PTFE (Teflon AF), and covalently crosslinked acrylamide (AA) gel. Data were collected with a triplicate set of samples (n=3). (*, p < 0.05; one-way analysis of variance with Holm-Sidak post-hoc test).

**Figure 6.**
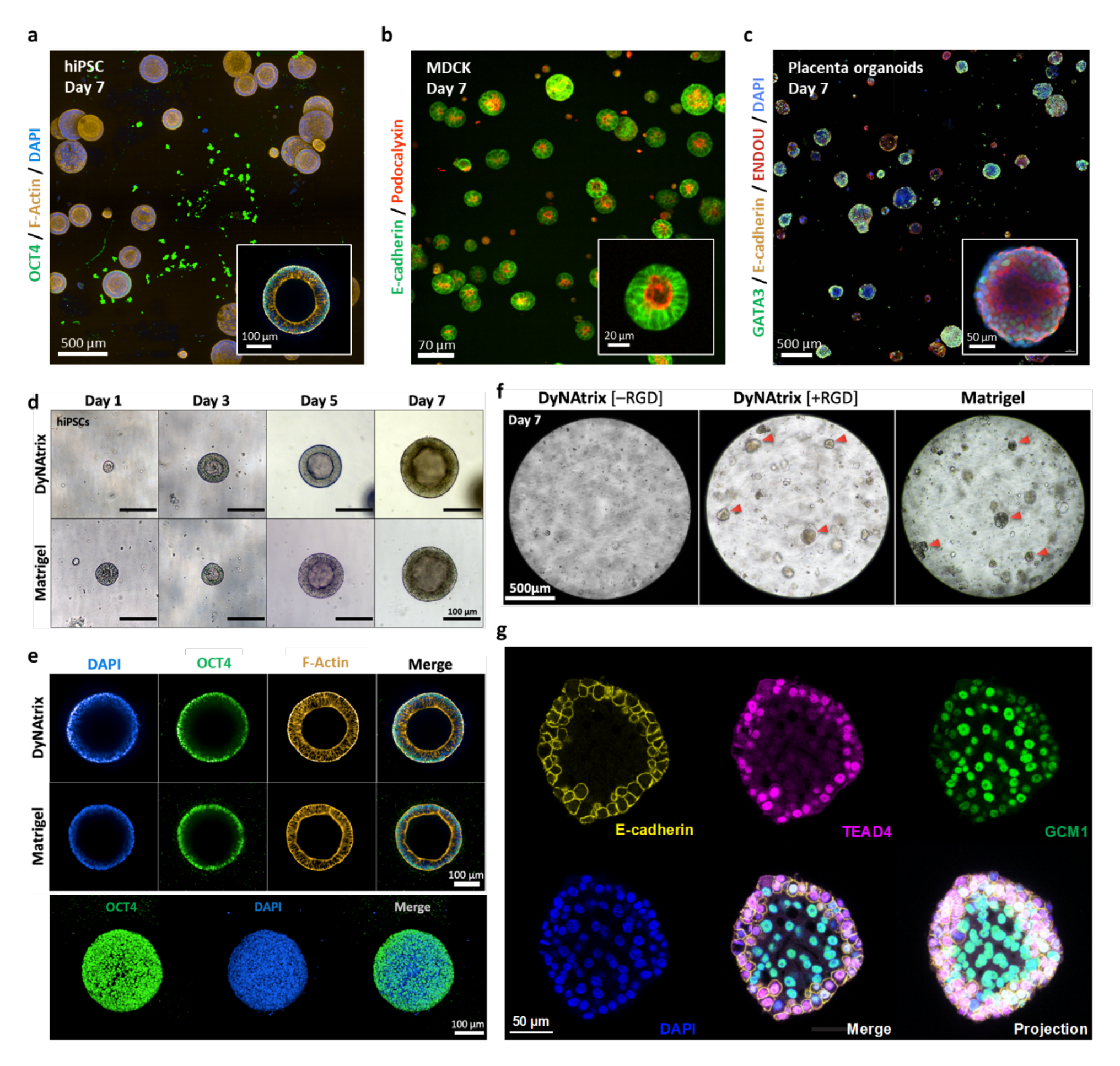
DyNAtrix is compatible with a variety of 3D cell and organoid cultures. Confocal images of (a) hiPSCs cysts embedded in DyNAtrix [+RGD]. Cells were stained for OCT4 (pluripotency), phalloidin (F-actin), and DAPI (nuclei), (b) MDCK cysts grown in DyNAtrix [+RGD] using heat-activated 18nt SRCs as crosslinker library. The cells were modified to express mNeonGreen-labelled E-cadherin and mScarlet-labelled Podocalyxin, and (c) Placenta organoids cultured in DyNAtrix [+RGD]. Cells were immunofluorescently stained for GATA3, E-cadherin, and ENDOU. (d) Representative microcopy images of hiPSC cysts grown in DyNAtrix [+RGD] in comparison with Matrigel. A comparison with DyNAtrix [–RGD] is shown in Supplementary Figure S15. (e) Representative confocal microcopy images of hiPSC cysts grown in DyNAtrix [+RGD] in comparison with Matrigel (day 7). Top: cross sections. Bottom: projection image of a cysts in DyNAtrix [+RGD]. (f) Representative brightfield images of placenta organoids grown in DyNAtrix [+RGD]/[–RGD] vs. Matrigel. Red arrows mark early organoids. (g) Confocal microscopy images of a placenta organoid in DyNAtrix [+RGD] stained for TEAD4, E-cadherin, GCM1 (day 7). A comparison with organoids in Matrigel is shown in Supplementary Figure S21.

### Matrix stability in cell culture, cytocompatibility, and tunable degradation

An immediate challenge for applying DNA-based materials in cell culture relates to their stability, as DNA is readily degraded by nucleases. DNase I, in particular, is present in fetal bovine serum (FBS) that is commonly added to culture media. It is thus necessary to protect DNA crosslinkers from DNase I activity, yet, on the flip side, a certain degree of enzymatic reconfigurability can be desirable. We therefore sought to identify conditions under which the enzymatic digestion of DNA-based gels can be either promoted or inhibited.

First, we tested the stability of unprotected gels in the presence of culture medium that was supplemented with 10% FBS. Gels were largely degraded within 48 hours (Supplementary Video 2). We then tested the effect of actin, which is a natural inhibitor of DNase I^53^. Using a fluorescently labeled DNA mock target, we showed that the rate of digestion by DNase I can be gradually tuned with different actin concentrations (Figure 5a). A concentration of 50 µg/mL was sufficient to suppress gel degradation for at least 48 hours (Figure 5b, Supplementary Video 3). 80 µg/mL actin reduced the rate of digestion by 97.5% (Supplementary Figure S25), preserving DyNAtrix in cell culture experiments for at least 2 weeks. A similarly effective protection can be achieved by the addition of citrate (10 mM), which chelates divalent cations that are required by DNase I. As the chelation of calcium ions was suspected to affect cell development, we chose actin as our default protection strategy. Supplementing both actin and DNase I to the culture provides external control over the material’s stability. This enables, for instance, gradual softening of the material or triggered release of cells at a chosen time point (Figure 5e).

To examine the cytocompatibility of the material, we first cultured mesenchymal stem cells (MSCs) in HAC-crosslinked and actin-protected 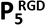 in DMEM + 10% FBS, and cell viability was probed by live/dead staining for up to one week. MSCs showed excellent viability (91±7% on day 7), significantly higher than on a reference 2D surface, and even outperforming Matrigel in most experiments (Figure 5c, Supplementary Figure S12). To demonstrate triggered release of live cells, 1 µL of recombinant DNase I solution (2 units / µL) was added onto the gel on day 9. The gel was promptly deconstructed within 2 hours, and live cells settled to the bottom of the culture plate (Figure 5e, Supplementary Video 4).

### Innate immune response and hemocompatibility

To further evaluate the properties of DyNAtrix as a biomaterial, we incubated HAC-64-crosslinked **P_5_** and 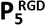 (1%, w/v) with fresh human whole blood and tested for various markers of inflammation and hemostasis. We compared these two gels to three reference substrates: (I) reactively cleaned glass, (II) Teflon™ AF (a benchmark material for medical devices), and (III) covalently crosslinked polyacrylamide (AA) gel. DyNAtrix exhibited low activation of monocytes (Figure 5f) and granulocytes (Supplementary Figure S11b,c), as indicated by low levels of CD11b expression. Similarly, we measured low levels of C5a, which is a marker for the activation of the complement system (Supplementary Figure S11a). Thus, DyNAtrix did not induce a significant innate immune response in vitro. Moreover, both gels showed very low release of platelet factor 4 (PF4, Figure 5f) and prothrombin fragment 1+2 (F1+2, Supplementary Figure S11d). The results indicate reduced blood coagulation and platelet activation with respect to glass and Teflon™ AF, and similar to the AA gel reference. These results suggest potential suitability for in vivo studies and medical applications. No statistically significant differences were observed between gels constructed from **P_5_** and 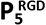.

### DyNAtrix supports proliferation and guides morphogenesis of diverse cell types and organoids

We finally sought to validate the suitability of DyNAtrix for advanced cell culture and organoid research. To this end, three different cell types were embedded in DyNAtrix [+RGD] (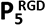 or 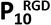, 1% (w/v), +HACs) and DyNAtrix [–RGD] (**P_5_** or **P_10_**, 1% (w/v), +HACs): (1) human induced pluripotent stem cells (hiPSCs), which offer patient-specific platforms for *in-vitro* disease modeling at the cellular level (Figure 6a,d,e), (2) Madin-Darby canine kidney (MDCK) cells, which serve as a model for developing epithelial tissues (Figure 6b), and (3) human trophoblast stem cells (hTSCs) that develop into placenta organoids, which allow in-vitro studies of human placental development and pregnancy-related disorders (Figure 6c,f,g). We assessed viability, proliferation and self-organization of the cells into multicellular structures and tested the effect of RGD adhesion sites on development and morphogenesis.

hiPSCs were cultured in DyNAtrix that was crosslinked in *mTeSR™1*, a chemically defined, serum-free medium optimized to maintain and expand hiPSCs. Notably, DyNAtrix did not require actin supplementation but stayed intact for the full culture duration (7 days). Similar results were obtained in other serum-free media with MDCK and trophoblast cultures (Figure 6b, c), indicating that the major source of nuclease is added serum, rather than secretion by cells. hiPSC viability in DyNAtrix and Matrigel were statistically indistinguishable (Supplementary Figure S13a). During the culture for one week, hiPSCs sustained continuous proliferation, formed hollow cysts that remained round-shaped, and continued expanding (Figure 6d). The size, morphology, and number of cysts appeared identical to those formed in a Matrigel reference culture. Confocal microscopy showed that the cysts strongly expressed pluripotency marker OCT4, further corroborating that the material is effectively supportive (Figure 6e). RGD peptides proved critical, as the number of observed cysts and their lumen size were greatly reduced in RGD-free DyNAtrix (Supplementary Figure S15), underlining the importance of mechanical hiPSC-matrix interactions.

We were initially skeptical whether it would be possible to stain the nuclear DNA of cells in DyNAtrix, as the presence of DNA crosslinkers could give rise to unacceptably high background fluorescence. However, due to the low DNA content, cell nuclei were readily stained with DAPI (4ʹ,6-diamidino-2-phenylindole) in all experiments (Figure 6a,e,g). Background fluorescence due to DNA crosslinker staining was negligible in 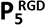, but could be detected for gels comprising 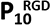 (Supplementary Figure S16).

The culture of MDCK cysts largely mirrored the results from hiPSC culture. MDCK cells were genetically modified to express fluorescently labeled E-cadherin and podocalyxin as markers for cell polarity and adhesion junctions. In initial experiments, cells were embedded in actin-protected DyNAtrix that was supplemented with 10% FBS. In order to remove any naturally-derived components, we also prepared DyNAtrix in a serum-free medium (UltraMDCK™), which led to identical results. In this environment, DyNAtrix remained stable for at least two weeks without nuclease protection. Confocal microscopy images showed that MDCK cells proliferate and form hollow cysts. The cells showed high viability on day 7 (Supplementary Figure S13b). As in hiPSCs, cyst morphologies strongly depended on the presentation of extracellular adhesion signals. Even though numerous cysts formed in DyNAtrix [–RGD], only 2.2% (±1.0%) of those cysts showed an apical-in polarity, while the majority exhibited an inverted or undetermined polarity (Supplementary Figure S17). In DyNAtrix [+RGD], cyst development increased to 20% (±1%) apical-in morphologies. In addition to the effect of adhesion signals, we also found that cyst polarity could be further modulated via the choice of the SRC. While initial studies were carried out with the default 14-nt-SRC, much higher levels of apical-in polarity (90%) were achieved with the 18nt SRCs (Figure 6b). This intriguing finding underlines the importance of mechanically programmable matrices to guide cell development. A detailed study on the effects of stress-relaxation on epithelial cell polarity in DyNAtrix is currently ongoing and will be published elsewhere.

We finally explored the development of human trophoblast stem cells in serum-free DyNAtrix. We chose this particular organoid model as the independently adjustable viscoelastic and biochemical cues could solve current challenges surrounding organoid polarity to more faithfully reproduce in-vivo morphologies^54^. In DyNAtrix [–RGD], trophoblast proliferation was low, and no organoids were observed (Figure 6f). In contrast, trophoblasts in DyNAtrix [+RGD] proliferated rapidly and had developed numerous placenta organoids within a week. The number, size, and shape of the organoids closely matched those grown in Matrigel at the same time point (Supplementary Figure S18). Placenta organoids in DyNAtrix and Matrigel showed similar viability (Supplementary Figure S13c). To demonstrate sustained proliferation, we adapted a recently reported protocol for long-term placenta organoid culture^54^. DyNAtrix crosslinkers were degraded with DNAse I on day 7 and the released organoids were successfully passaged two consecutive times, resulting in a total culture duration of 21 days (Supplementary Figures S23-24). The organoid morphology in DyNAtrix and Matrigel was investigated in immunofluorescence images obtained from confocal microscopy (Figure 6c,g, Supplementary Figure S19). We found strong expression of GATA3, a trophoblast-specific transcription factor typically found in all cells of placenta organoids (Figure 6c). Moreover, we confirmed the expression of GCM1, TEAD4, Syndecan, ENDOU, and E-cadherin, which are important markers for cell differentiation and organoid development^55, 56^ (Figure 6g, Supplementary Figures S19-22).

Overall, confocal imaging did not reveal any morphological or developmental differences of placenta organoids in DyNAtrix when compared to Matrigel. To the best of our knowledge, this is the first report describing the successful growth of human placenta organoids in a fully synthetic 3D matrix environment. Beyond the scope of this article, future studies will focus on utilizing the independently tunable stress-relaxation, elasticity, and adhesion signals in DyNAtrix to improve the physiological relevance of this organoid model for in-vivo placental development.

## Discussion

In this report, we have illustrated the power of DNA nanotechnology for soft materials engineering toward programmable and adaptive 3D cell culture matrices. The key feature of DNA is its unique potential for rational material design, allowing researchers to directly translate computationally predictable molecular properties into macroscopic material features: we show that the sequence and complexity of combinatorial crosslinker libraries (CCLs) determine network formation (Figure 2), thermodynamic stability (Figure 2b, Figure 3a), and elasticity (Figure 2c); stress relaxation crosslinkers (SRCs) control the (nano)mechanical stability (Figure 3); heat-activated crosslinkers (HACs) demonstrate customizable binding kinetics, which is critical for homogeneous cell encapsulation under optimal culture conditions (Figure 4). The concepts of CCLs, SRCs, and HACs can be combined, and the materials’ modularity allows future upgrades with supplementary DNA components that either tweak mechanical properties or introduce entirely new features. Achieving this exceptional level of control helps mimic many of the properties of biological tissues and naturally-derived matrices, but in a fully synthetic and compositionally defined material. This improves reproducibility and reduces biological complications and regulatory hurdles in the field of regenerative medicine. DyNAtrix exhibits high cyto-and hemocompatibility, is self-healing, and suitable for injection and 3D extrusion printing. All of its components can be easily sterilized by filtration or alcohol treatment and stored (ready-to-use) for years.

DNA-based soft materials with remarkable adaptive properties had been previously reported^5, 8, 57^. However, surprisingly few studies had explored their use in 3D cell and tissue culture^50, 58–60^. Problems relating to high required DNA content, premature gel degradation, and seamless integration with existing cell culture workflows had remained largely unsolved. In the present study we show that the combination of a UHMW polymer backbone with CCLs enables very low DNA concentrations—up to 2 orders of magnitude lower than previously reported synthetic DNA-crosslinked gels. This approach gives access to soft gels with elastic moduli in the range of 1–500 Pa, which are suitable for the study of many mechano-sensitive stem cell and organoid systems^61^. The lower end of the stiffness regime is particularly relevant for systems that benefit from ultra-soft environments, such as placenta organoids (as shown in this study) and stem cell models for embryo development. Due to the low DNA content, DyNAtrix is non-immunogenic, making it potentially suitable for medical applications, such as injectable cell-laden gels, drug release systems, or coatings for medical devices. The low DNA content also enables scalable high-throughput applications at relatively low material cost (Supplementary Table S4). Though DyNAtrix supported all tested cell types in this study, we note that it currently carries only a single type of adhesion ligands. In the future, it may become necessary to attach multiple ligands to the backbone to better mimic the complex adhesion properties and signaling of the native ECM.

The construction of biocompatible hydrogels that combine dynamic viscoelastic properties with long-term stability in cell culture has been a major challenge^21^. Importantly, the degradation of DyNAtrix under mild cell-compatible conditions can be either inhibited (by supplementing actin) or promoted (by supplementing DNase I). Actin protection—or the use of serum-free media—allows cell culture for prolonged periods of time. In particular, placenta organoids were cultured in DyNAtrix for up to three weeks, which is a new record for DNA-crosslinked ECM mimics. Such long culture times are critical for future studies on human organoid systems. The nuclease/actin-regulated deconstruction of DyNAtrix (and optional re-construction via molecular self-assembly) represent an externally controllable alternative to the commonly relied-upon matrix degradation by cell-secreted matrix-metalloproteinases.

Previous studies had demonstrated control over stress relaxation via chemical modifications to the polymer scaffold or side groups^26, 62–66^. Moreover, a DNA-crosslinked hydrogel had been shown to exhibit temperature-dependent stress-relaxation^67^. In cell culture, however, temperature cannot be altered at will, and the effects of chemical matrix modifications on cell development can be difficult to disentangle from pure stress relaxation effects. With the help of SRCs, for the first time the stress relaxation of a material can be adjusted systematically over a wide range by altering merely a few nucleobases on the DNA crosslinker sequence. The effects of different stress-relaxation times on the development of various cell and organoid models is subject of ongoing studies in our lab.

In summary, DyNAtrix allows a simple yet fully controlled noncovalent mix-and-match approach to matrix engineering. DNA modules and polymer backbones with different properties can be deployed in various combinations. For example, a HAC-crosslinked DyNAtrix can be further upgraded with fluorescent DNA stress sensors^68, 69^ to shed light onto the mechanical interactions of cells with their environment. DNA aptamers can be integrated to scavenge (or gradually release) specific biomolecules. Such modules can be activated on demand (e.g., by temperature), modified (e.g., by hybridization), and subsequently released off the polymer scaffold (e.g., by restriction enzymes). We note that the chosen polymer backbone has been previously shown to facilitate dynamic capture and release of nucleic acids from biological samples^39^. Therefore, the polymer matrix could in principle be directly repurposed after cell culture (e.g., via addition of highly diverse catcher strand libraries^70^) to isolate and process genomic DNA or mRNA to assist downstream sequencing analysis. In the future, DyNAtrix could be autonomously controlled by molecular logic gates and synthetic regulatory circuits based on DNA strand displacement cascades^49, 71^. We therefore envision this novel class of bioengineered materials to create unprecedented options for 3D cell and organoid culture and a range of other biomedical applications.

## Abbreviations

AA: Polyacrylamide
CCL: Combinatorial crosslinker library
CGC: Critical gelation concentration
DyNAtrix: Dynamic DNA-crosslinked matrix
ECM: Extracellular matrix
F1+2: Prothrombin fragment 1+2
FBS: Fetal bovine serum
HAC: Heat-activated crosslinker
hiPSC: Human induced pluripotent stem cell
hTSC: Human trophoblast stem cell
MDa: Megadalton
MDCK: Madin-Darby canine kidney
MSC: Mesenchymal stem cell
nt: nucleotide
PF4: Platelet factor 4
RGD: Arginylglycylaspartic acid
SRC: Stress-relaxation crosslinker
UHMW: Ultra-high molecular weight
w/v: weight to volume ratio

## Acknowledgements

E.K. acknowledges funding by the Federal Ministry of Education and Research of Germany (BMBF) in the program NanoMatFutur (grant no. 13XP5098). The authors thank the medical faculty at the University Hospital Dresden for providing MSCs, and CRTD Stem Cell Engineering Facility for providing hiPSCs. We would like to thank the following Services and Facilities of the MPI-CBG for their support: organoid and stem cells and light microscopy facilities. We thank Andrea Meinhardt, Aukha Stoppa, Dr. Yanuar Dwi Putra Limasale, Prannoy Seth, Markus Mukenhirn, and Dr. Valentina Magno for helpful discussions. We also thank Dr. Ron Orbach, Dr. Agnes Kriegne Toth-Petroczy, Dr. Michele Marass, and Prof. Boris Rybtchinski for giving valuable feedback on the paper draft.

## Author contributions

E.K. conceived the project. Y.-H.P., K.G., S.K.H., and E.K. designed the experiments. Y.-H.P. carried out sequence design, rheological experiments, gelation, MSC and hiPSC culture. S.K.H. carried out polymer synthesis and characterization, as well as MDCK culture and imaging. K.G. carried out DNA sequence design, thermodynamic predictions and early gelation experiments. A.R. performed peptide synthesis and characterization. G.K.A. helped plan and interpret rheological experiments. M.M. and S.K.H. carried out and analyzed blood incubation experiments. S.K.H., Y.-H.P., C.G., and J.L. helped plan, carry out and interpret trophoblast culture experiments. S.B. carried out and analyzed asymmetrical flow field-flow fractionation experiments. A.H. developed the genetically modified MDCK cells and helped with data analysis. C.W. helped plan and interpret matrix engineering and cell culture experiments. Y.-H.P. and E.K. wrote the initial manuscript draft. All authors discussed the results and helped revise the manuscript.

## Data availability

Supplementary information containing materials and methods, supplementary figures, tables, datasets, and accession numbers for biological materials are available with this paper. Additional datasets and materials generated during and/or analyzed during the current study are available from the corresponding author on reasonable request.

## Code availability

A python script for the statistical simulation of the maximum percentage of intramolecular crosslinks as a function of CCL complexity is available as Supplementary File 11. A python script for the thermodynamic calculations of CCL interactions is available as Supplementary File 12.

## Ethics statement

Informed consent was obtained from all recipients and/or donors of cells or tissues. The study involving human whole blood was covered by the ethic vote EK-BR-24/18-1 of the Sächsische Landesärztekammer. The blood was obtained from two voluntary ABO-matched donors who had not used any medicine in the past ten days. The study involving Human bone marrow–derived MSCs was covered by the ethic vote EK221102004 and EK47022007 at TU Dresden. MSCs were isolated from healthy female/male donors (aged 26–37) by the Medical Faculty at the University Hospital Dresden. The study involving hiPSCs was covered by the ethic vote EK363112012 at TU Dresden. hiPSCs were generated from MACS-sorted CD34+ cells from the peripheral blood of a healthy donor (aged 20-24). The CT27 patient-derived trophoblast stem cell line used in this study had been derived from placental cytotrophoblast cells and was obtained from the RIKEN stem cell bank (RCB4936:CT27). Human placentas were obtained from healthy women with signed informed consent of the donors, and the approval of the Ethics Committee of Tohoku University School of Medicine (Research license 2014-1-879).

