## Supplementary Information for "Dynamic matrices with DNA-encoded viscoelasticity for advanced cell and organoid culture"

### 1 Materials

Solvents and reagents were purchased from commercial sources and used as received, unless otherwise specified. Water was obtained from a Milli-Q system. Methanol (MeOH, ACS reagent grade) was obtained from Fisher Scientific. Molecular biology grade acrylamide (AA, Cat. #A9099), N,N´-methylenebisacrylamide (BAA, Cat. M7279), sodium acrylate (A, Cat. #408220), 19:1 acrylamide/bis-acrylamide (Cat. # A2917), and ammonium persulfate (APS, Cat. #A3678) were purchased from Sigma-Aldrich. Ultrapure N,N,N´,N´-tetramethylethylenediamine (TEMED, Cat. #15524010) and SYBR™ Gold Nucleic Acid Gel Stain (Cat. #S11494) were obtained from Thermo Scientific. The SequaGel-UreaGel System was purchased from National Diagnostics (Cat. #EC-833). Desalted oligonucleotides were purchased from Integrated DNA Technologies (IDT). DNA ladders were purchased from Thermo Fischer (Cat. # SM1211). Nitrogen gas (>99.999%) was used for experiments under inert conditions. Nitrogen gas was supplied by an in-house gas generator. To ensure an inert condition, it was purified through a Model 1000 oxygen trap from Sigma-Aldrich (Cat. #Z290246). All reagents containing unreacted acrylamide groups were stored at 4 or −20 °C, and protected from unnecessary exposure to light. Actin protein (>95% Pure, rabbit skeletal muscle, Cat. # AKL95) was purchased from Cytoskeleton, Inc. and DNase I (Cat. # M0303) was purchased from New England Biolabs, Inc..

Human bone marrow–derived MSCs were isolated from healthy female/male donors (aged 26–37) by the Medical Faculty at the University Hospital Dresden after obtaining their informed consent (Ethikkommission an der Technischen Universität Dresden, Ethic board no. EK263122004). MDCK II cells (ECACC 00062107) were provided by Alf Honigmann's group. Human pluripotent stem cells were generated in the "CRTD Stem Cell Engineering Facility, Technische Universität Dresden" by Shahryar Khattak and registered under the hPSCreg name CRTDi003-B (https://hpscreg.eu/cell-line/CRTDi003-B). The CT27 patient-derived trophoblast stem cell (TSC) line used in this study was obtained by the RIKEN stem cell bank (https://cellbank.brc.riken.jp/cell_bank/CellInfo/?cellNo=RCB4936). The CT27 TSCs were derived from placental cytotrophoblast cells as described in Okae et. al 2018 (https://doi.org/10.1016/j.stem.2017.11.004). Human placentas were obtained from healthy women with signed informed consent of the donors, and the approval of the Ethics Committee of Tohoku University School of Medicine (Research license 2014-1-879).

### 2 Methods

#### 2.1 Polymer synthesis

A general protocol can be found in reference [1]. In brief, acrylamide (50 mg/mL), sodium acrylate (0.5 mg/mL), and acrylamide-labeled anchor strand DNA (Table S1) were co-polymerized in TBE buffer (100 mM Tris, 100 mM boric acid, 2 mM EDTA, pH 8.2). For $\mathbf{P_1}$, $\mathbf{P_5}$, and $\mathbf{P_{10}}$ DNA concentrations in solution were 100 µM, 500 µM, and 1000 µM, respectively. For the corresponding RGD-functionalized derivatives $\mathbf{P_5^{RGD}}$ and $\mathbf{P_{10}^{RGD}}$, 10mM of acrylated RGD peptide (sequence: G(acryl-K)GGGRGDSP) was co-polymerized with the above solution. The polymerization reaction for $\mathbf{P_1}$, $\mathbf{P_5}$, and $\mathbf{P_5^{RGD}}$ was initiated with 0.005 wt% ammonium persulfate (APS) in presence of 0.005 wt% tetramethylethylenediamine (TEMED). Synthesis of $\mathbf{P_{10}}$ and $\mathbf{P_{10}^{RGD}}$ was initiated by 0.025 wt% TEMED and 0.025 wt% APS. In order to achieve high molecular weight and a narrow size distribution, it was necessary to carry out the reaction in high-purity nitrogen gas, which was passed through an oxygen trap on-site. The reaction was allowed to proceed overnight. A highly viscous polymer solution was obtained, indicating the formation of long polymer chains. NMR spectroscopy was used to verify high conversion of the monomer (Figure S3). The solution was diluted in 9 volumes of TE buffer (10 mM Tris, 1 mM EDTA, pH 8.0) and subsequently purified via methanol precipitation [1]. The pellet was resuspended in milliQ water and stored in aliquots at −20 °C. A small aliquot of the polymer solution was lyophilized and weighted to determine the synthesis yield. Yields: 89% (**$P_1$**), 98% (**$P_5$**), 93% (**$P_{10}$**), 92% (**$P_5^{RGD}$**), and 88% (**$P_{10}^{RGD}$**).

#### 2.2 Nearest-neighbor thermodynamic predictions

Explicit sequences of crosslinking strands and their complements were generated using a custom python script. The program sequentially replaces the ambiguous bases N with nucleobases (A, C, G, T) via multi-layer nested loops. This created a library of forward and reverse splint variants. The minimum free energies (MFE) of all explicit overlap domains were calculated against all explicit overlap domains in the library using NUPACK (version 4.0.0.27), using nearest-neighbor thermodynamic parameters according to reference [2]. The model parameters were set to T = 20°C, 150 mM NaCl, 75 µM total DNA concentration (37.5 µM for each splint strand), and a complex size of 2.  For each possible pair, the **Boltzmann distribution factor** $B_{x,y}$ was sequentially calculated for each variant of forward (x) against all possible reverse splint (y) strand at the expected theoretical concentration using the equation

$$B_{x,y} = \mathrm{e}^{\frac{-\mathrm{MFE(x,y)}}{kT}}$$

The normalized distribution was calculated to study the selective affinity of any forward splint variant in a pool of reverse splint sequences.  This was calculated for each pair as the ratio of distribution factor of one pair (x,y) by the sum of all pairs as

$$B_{x,y}^{normalized} = \frac{B_{x,y}}{\sum_y B_{x,y}}$$

We note that a precise solution can only be obtained by numerically solving a system of ordinary differential equations[2]. For simplification, we assumed that each splint strand is available at its initial concentration. In reality, non-complementary splint strand concentrations are expected to be significantly reduced, since they would be predominantly scavenged by their complementary binding partners. The obtained values for specific CCL pairs in the Boltzmann distribution that are based on our simplified calculation are therefore considered a lower limit for relative abundances.

#### 2.3 Polyacrylamide gel electrophoresis (PAGE)

The binding capacity of **$P_5$** was determined by PAGE. Samples were prepared in 1x TE buffer with a final concentration of 150 mM NaCl, and 0.4 %(w/v) solutions of **$P_5$** (2 µM of anchor strands). The complementary strand of the anchor strand (Strand ID#1) was added at series equivalent from 0.2x~ 2x. The samples were annealed on the thermal cycler at 95°C for 2 minutes, followed by instant cooling to 4°C and hold for 3 minutes. The annealed products were loaded onto a 10% native polyacrylamide gel and run at 120 V on a Mini-Cell®Novex system from Life technologies in 0.5× TBE buffer, using Consort EV265 power supply. The gel was stained with 1x SYBR™Goldfor 15 minutes and scanned on Typhoon FLA 9000 Scanners (GE Healthcare Life Sciences) using a blue LD laser (excitation at 473 nm) at 25 µm/pixel resolution. Image analysis and densitometric quantification of gel bands were performed in Fiji[3].

#### 2.4 Asymmetrical flow field-flow fractionation

Asymmetrical flow field-flow fractionation with light scattering detection (AF4-LS) has become an important technique for gentle and detailed characterization of bio-active systems[4]. We performed the AF4 studies with an Eclipse Dualtec system (Wyatt Technology Europe, Germany). The separation takes place in a hollow fiber, a 17-cm-long polyethersulfone fiber, (Microdyn-Nadir, Germany) with 0.8 mm inner diameter, 1.3 mm outer diameter, and 10 kDa molecular weight cut-off (corresponding to an average pore diameter of 5 nm). Flows were controlled with an Agilent Technologies 1260 series isocratic pump equipped with vacuum degasser. The detection system consisted of a multiangle laser light scattering detector (DAWN HELEOS II from Wyatt Technology Europe, Germany), operating at a wavelength of 659 nm with included QELS module (at detector 99°) and an absolute refractive index detector (Optilab T-rEX, Wyatt Technology Europe GmbH, Germany), operating at a wavelength of 659 nm. All injections were performed with an autosampler (1200 series; Agilent Technologies Deutschland GmbH). The channel flow rate (Fc) was maintained at 0.35 mL/min for all Af4 operations. Samples (inject load: ~50-100 µg) were injected during the focusing/relaxation step within 5 min. The focus flow was set to 0.85 mL/min. During the elution step, the cross-flow rate (Fx) was optimized by an exponential cross-flow gradient of 0.85 - 0.03 mL/min in 30 min. 10 mM TRIS buffer and 1 mM EDTA (pH 8.0) were used as eluent for all measurements. Collecting and processing of detector data were made by the Astra software, version 6.1.7 (Wyatt Technology, USA). The molar mass dependence of elution time was fitted with Berry (first-degree exponential).

#### 2.5 Nuclear magnetic resonance (NMR) spectroscopy

NMR samples were prepared by diluting 100 µL of the unpurified product in 700 µL of $D_2O$. $^1H$ NMR spectra were recorded at 30-32°C on a 500 MHz spectrometer (Bruker) with 2 seconds acquisition time and 32 transients. Chemical shifts (δ) are reported in parts per million (ppm) downfield from tetramethyl silane (TMS). $^1H$ NMR shifts are referenced to the residual hydrogen peak of $D_2O$ (4.79 ppm).

#### 2.6 Solid-phase peptide synthesis (SPSS)

All SPPS chemicals were purchased from IRIS Biotech GmbH. RGD peptides (sequence: G(acryl-K)GGGRGDSP) were synthesized on a Liberty Blue HT12™ automatic and microwave-assisted peptide synthesizer (CEM GmbH) using a Rink Amide resin and a 9-fluorenylmethoxycarbonyl protection strategy. Amino acid activation was achieved by N, N'-diisopropyl carbodiimide, and ethyl cyanohydroxyiminoacetate. Acetylation of the N-terminus was performed via incubating the resin-bound peptides in acetic anhydride for 2 h. The reaction mixture was continuously stirred and vigorously saturated by bubbled nitrogen to prevent oxidation reactions and afterward washed three times with N, N-dimethylformamide. Deprotection of the amino acid side chains and cleavage from the resin was accomplished using a mixture of trifluoroacetic acid ($\phi_i$ = 87.5%), phenol ($\phi_i$ = 5%), triisopropylsilane ($\phi_i$ = 2.5%) and MilliQ-water ($\phi_i$ = 5%) for 3 h at room temperature. The crude peptide was precipitated in anhydrous diethyl ether, collected by vacuum filtration, and dried under nitrogen flow. Further purification of the peptides was realized by high-performance liquid chromatography (Agilent 1200, Agilent Technologies) on a preparative C18 column (10 µm particle size, 100 Å pore size, 250 x 30 mm, Phenomenex Ltd). A linear gradient of MilliQ-water/acetonitrile and trifluoroacetic acid ($\phi_i$ = 0.1%) was used as the mobile phase. Finally, the concentrated peptide-containing solution was lyophilized (Alpha 2-4 LD plus freeze-dryer, Martin Christ) and the obtained product was stored at -20°C afterward until further usage.

#### 2.7 Assembly of DNA-crosslinked hydrogels for rheological measurements

Unless specified, the samples were prepared at the final concentration of 1% (w/v) polymer solution, added with one equivalent of DNA crosslinkers to the anchor strand (e.g., 75 µM crosslinkers for 1% (w/v) $P_5$ solution), and mixed in the buffer condition of 150 mM NaCl and 1x TE buffer. The samples were annealed on a BioRad C1000 Touch Thermal Cycler with the following steps:

1. Heating at 95°C for 3 minutes
2. Instant cooling from 95°C to 80°C
3. Holding at 80°C for 2 minutes
4. 1$^{st}$ cooling ramp from 80°C to 65°C at -0.3°C/min.
5. 2$^{nd}$ cooling ramp from 65°C to 37°C at -0.5°C/min.
6. Holding at 37°C for 24 hours before rheological measurements.

The first cooling ramp allows the adaptor domains of the crosslinkers to bind to the anchor strands, while the second ramp allows the overlap domains to properly bind to their complementary partners.

#### 2.8 Oscillatory rheological measurements

Viscoelasticity measurements were performed using an Anton Paar MCR301 rheometer with a 25 mm diameter cone-plate geometry and a cone angle of 0.5°. A temperature-controlled hood (Peltier system PTD-200, Anton Paar) was installed to provide an accurate temperature setting in the measuring area. A wet paper cylinder was placed around the circumference of the plate to maintain the humidity very close to saturation in the hood. This suppresses evaporation artefacts. Amplitude sweeps were performed from 0.1% to 1000% strain at 1.6 Hz. Frequency sweeps were performed from 0.1 Hz to 100 Hz at 10% strain. Temperature sweeps were performed from 20°C to 60°C at 10% strain and from 60°C to 70°C at 20% strain, at 1.6 Hz frequency. The heating and cooling steps were run reversibly for two cycles to exclude evaporation artefacts. Unless specified, otherwise, samples were measured in triplicates to ensure reproducible results. Stress-relaxation properties were measured at 15% strain. The strain was held constant while the shear stress was recorded as a function of time. Self-healing tests were conducted at 1.6Hz by altering 1000% and 10% strain for 5 minutes each and repeating for 4.5 times. For the heat-activated gelation tests, two precursor components were prepared at 1% (w/v) $\mathbf{P_5}$ mixed with: (1) 40 µM forward CCL-64 strands (Strand ID #6a) and 80 µM corresponding blocking strand (Strand ID #15a), and (2) 40 µM reverse CCL-64 library (Strand ID #6b) and 80 µM corresponding blocking strand (Strand ID #15b). Two precursors were heated at 95°C for 3 min, slowly annealed with a cooling ramp from 95°C to 4°C at -3°C/min, and kept at 4°C overnight to equilibrate. Before rheological measurement, two precursors were mixed on ice with a positive displacement pipette. After mixing, the samples were subjected onto the rheometer. The samples was held at 4°C for 5 minutes, then instantly heated up to 37°C, and held at 37°C for 60 minutes while shearing at 10% strain and 1.6 Hz frequency.

#### 2.9 Hydrogel printing

Before 3D printing, a microscope glass slide (76 × 26 mm, Thermo Scientific) was attached to the printing bed of the device. 200 µL hydrogel stained with SYBR Gold was transferred to a disposable 1 ml syringe (Omnican® 40) with a 30-guage needle (Omnican®). The syringe was centrifuged invertedly at 300 Xg for 2 min to remove air bubbles. The hydrogels were printed on BioScaffolder BS5.1 (GeSiM) at a speed of 2 mm/s, an extrusion rate of 40 µm/s and a layer height of 0.12 mm at room temperature. Immediately after printing, the glass slide was transferred to Opera Phenix Plus High-Content Screening System to acquire 3D structures at 5x magnification.

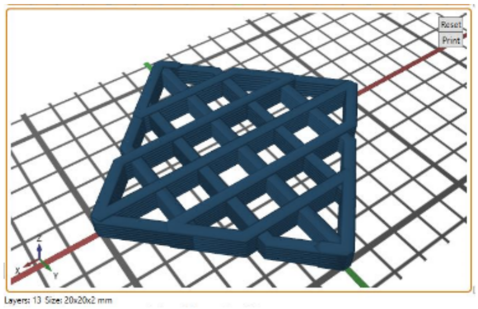


Scaffold design: The size of the grid is 20 x 20 x 2 mm (L x W x H)

#### 2.10 Mixing tests with the heat-activated crosslinkers

Two precursor solutions were prepared in 1x TE buffer at the final concentration of 1% (w/v) **$P_5$** with: (1) 40 µM forward CCL-64 strands (Strand ID #6a) and 80 µM corresponding blocking strand (Strand ID #15a), and (2) 40 µM reverse CCL-64 library (Strand ID #6b) and 80 µM corresponding blocking strand (Strand ID #15b). To visualize the mixing quality of the two precursors, 10 µM of a 6-FAM (Fluorescein)-modified strand (Strand ID #15) was added to the precursor. Both precursors were incubated a 4°C for 1 hour. After equilibration, two precursors were mixed with a positive displacement pipette and subjected 10µL/well to a 384-well plate (Grenier bio-one). The plate was incubated at 37°C for 30 minutes to initiate the gelation process. Finally, the images were acquired on Dragonfly High-Speed Confocal Microscope System (Oxford instrument) at 4x magnification.

#### 2.11 Swelling degree measurements

Lyophilized actin protein (Cytoskeleton inc., Cat. #AKL95-B) was reconstituted to 10 mg/mL with 100 µl of deionized water. After reconstitution the actin will be in the following buffer: 5 mM Tris-HCl pH 8.0, 0.2 mM $CaCl_2$, 0.2 mM ATP, 5% (w/v) sucrose, and 1% (w/v) dextran. Citric acid powder (192.124 MW) was prepared in 1x PBS to a final concentration of 200 mM, pH 7.0. The gels (1% (w/v) $P_5$ crosslinked with CCL-64) were stained at 10X SYBR Gold before swelling. 5 µl of the gels were subjected to a 384-well microplate (Greiner, No.: 788092). The samples were immersed in 20 µl of (i) 1x TE buffer, (ii) 10% FBS-containing DMEM, (iii) 50 µg/mL actin + 10% FBS-containing DMEM, or (iv) 10 mM citrate + 10% FBS-containing DMEM. The images were acquired every hour with a high-content spinning disk confocal microscope (Opera Phenix Plus High-Content Screening System, PerkinElmer) at 5x magnification for 48 hours at 37°C. The gel volume was quantified by the software IMARIS (Oxford Instrument) using the surface wizard and its statistic tool under a uniform threshold setting.

#### 2.12 Enzymatic digestion measurements

The FRET-paired probe that contains Cy5 fluorophore and Iowa Black® Dark quencher (Q) was purchased from IDT. 2 µM of Cy5-strand (Strand ID #13a) and 4 µM of Q-strand (Strand ID #13b) were mixed in 1x PBS buffer. The FRET-paired probe was annealed by the following program: heating at 95°C for 1 minute, instant cooling to 50°C for 2 minutes, and slowly cooling from 50°C to 20°C at -1.5°C/min. We prepared the samples at the final concentration of 100 nM Cy5-Q FRET probe, with increasing actin content from 2.5 to 320 µg/mL (Cytoskeleton inc., Cat. #AKL95-B) in the buffer condition of 150 mM NaCl, 1.8 mM CaCl2, 0.2 mM ATP (Jena Bioscience, Cat. # NU-1010) and 1x IDTE (IDT). 80 U/mL DNase I (NEB, Cat. # M0303S) was added to each sample shortly before measurement. We loaded 20 µl/well of each samples in triplicates into a 96-well qPCR plate. The fluorescence signals were read every 10 minutes for 24 hours and every 30 minutes for the following 36 hours at 37°C on a CFX96™ Real-Time PCR System (Bio-Rad).

#### 2.13 Whole blood incubation

Whole blood incubation was performed as described previously[5,6]. Gels were prepared on glass carriers with a poly(ethylene-alt-maleic anhydride) bonding layer[7], and inserted in in-house developed incubation chambers made of poly(tetrafluoroethylene), where 3.2 $cm^2$ test surface were exposed to 2 ml blood[5]. Reactively cleaned glass (RCA)[8] and Teflon™ AF (DuPont, USA) served as activating and inert reference surfaces, respectively.

Blood was obtained from two voluntary ABO-matched donors who had not used any medicine in the past ten days. The blood was immediately anticoagulated with 1.0 U mL-1 heparin (Ratiopharm, Ulm, Germany), pooled, and filled into the incubation chambers with the samples, avoiding an air interface. The chambers were incubated for two hours at 37°C at constant overhead rotation to prevent sedimentation. The blood was subsequently analyzed for the blood cell count (Beckmann Coulter AcTdiff, Krefeld Germany). Granulocyte and monocyte activation were determined by flow cytometry (LSR Fortessa, Becton Dickinson, Heidelberg, Germany), where granulocytes and monocytes were identified for their scatter characteristics and CD15 (clone SSEA-1, PE-conjugated, BioLegend, San Diego, Ca, USA) and CD14 (clone M5E2, APC conjugated, Becton Dickinson) positivity, respectively. The intensity of CD11b expression (clone ICRF44, PacificBlue-conjugated, BioLegend) was normalized to blood incubated with 100 EU/mL endotoxin. The rate of granulocytes with conformationally activated CD11b was determined using anti-CD11b clone CBRM1/5 (PE/Cy7-conjugated, BioLegend).

Soluble markers of coagulation activation (prothrombin fragment F1+2), blood platelet activation (platelet factor 4 (PF4)) and complement activation (complement fragment C5a) were determined from plasma using commercial ELISAs (Enzygnost F1+2, Siemens Healthineers, Munich, Germany; Zymutest PF4, Hyphen BioMed, Neuville-sur-Oise, France; C5a ELISA, DRG Instruments, Marburg, Germany) after stabilization of the blood with the recommended additives of the testkits and centrifugation.

The analysis was performed with a triplicate set of samples in parallel (n=3). The study was covered by the ethic vote EK-BR-24/18-1 of the Sächsische Landesärztekammer.

#### 2.14 General cell culture on 2D

##### 2.14.1 Mesenchymal stem cells (MSCs)

Mesenchymal stem cells (MSCs) were cultured in Dulbecco's Modified Eagle's medium (DMEM) GlutaMAX™ (Gibco, Cat. #21885025) supplemented with 10% Fetal Bovine Serum (FBS) (Sigma Aldrich, Cat. # F7524) and 100U Penicillin and 0.1 mg Streptomycin (Sigma Aldrich, Cat. #P4333) at 37°C and 5% CO2 in a humidified incubator. The cells were grown until 80%-90% confluence in T75 flask before detachment with 0.25% trypsin-EDTA solution.

##### 2.14.2 Madin–Darby Canine Kidney cells (strain II, MDCK II)

Auto-fluorescent Madin–Darby Canine Kidney cells (strain II, MDCK II) were provided by Alf Honigmann's group (MPI-CBG). These cells express E-cadherin-mNeonGreen and podocalyxin-mScarlet. The cells were cultured in minimum essential medium (MEM) with GlutaMAX™ Supplement (Gibco, Cat. #11140050), 1%MEM Non-Essential Amino Acids Solution (100X) (Gibco, Cat. #41090028), 1mM Sodium Pyruvate (Gibco, Cat. #11360070), 5% FBS (Sigma Aldrich, Cat. # F7524), and 100U Penicillin and 0.1 mg Streptomycin (Sigma Aldrich, Cat. #P4333) at 37°C and 5% CO2 in a humidified incubator. The cells were grown until 80%-90% confluence in T75/T25 flask before detachment with 0.25% trypsin-EDTA solution.

##### 2.14.3 Human induced pluripotent stem cells (hiPSCs)

Human induced pluripotent stem cells (hiPSCs) were cultured in mTeSR™1 (StemCell, Cat. # 85857) supplemented with 100U Penicillin and 0.1 mg Streptomycin (Sigma Aldrich, Cat. #P4333) on a 6-well plate coated with Matrigel (Corning, Cat. #354277). The cells were grown until 90% confluence and then detached by ReLeSR (StemCell, cat # 05872). The cell suspension was split at 1:4 or 1:6 ratio and seeded onto a new coated plate. For encapsulation, the obtained cell suspension was transferred to a low-adhesion 6-well plate to form cell aggregates in mTeSR™1 supplemented with 8 µM ROCK inhibitor Y-27632 (StemCell, cat # 72304) overnight.

##### 2.14.4 Human trophoblast stem cells (hTSCs)

Human trophoblast stem cells (hTSCs) were cultured as described previously[9]. In brief, a 6-well plate was coated with 5 µg/mL Col IV at 37°C for at least one hour. The cells were seeded and cultured in the TSC medium [DMEM/F12 supplemented with 0.1 mM 2-mercaptoethanol, 0.2% FBS, 0.5% Penicillin-Streptomycin, 0.3% BSA, 1% ITS-X supplement, 1.5 µg/mL L-ascorbic acid, 50 ng/mL EGF, 2 µM CHIR99021, 0.5 µM A83-01, 1 µM SB431542, 0.8 mM VPA and 5 µM Y27632] at 37°C and 5% CO2 in a humidified incubator. The TSC medium was replaced every two days. The cells were grown until 80% confluence before detachment with TrypLE (Thermo Fisher Scientific, Cat#12604031).

#### 2.15 3D Cell culture

##### 2.15.1 DyNAtrix precursors preparation

All materials were dissolved in water and stored at -20°C for cell culture experiments. For MSC and hiPSC cultures, two precursors were prepared at a final concentration of 1% (w/v) $\mathbf{P}_{\mathbf{5}}^{\mathbf{RGD}}$ with: (1) 37.5 µM forward splint (Strand ID #6a) and 75 µM blocking strand (Strand ID #15a), and (2) 37.5 µM reverse splint strands (Strand ID #6b) and 75 µM blocking strand (Strand ID #15b). As for MDCKII and hTSC cultures, two precursors were prepared at a final concentration of 1% (w/v) $\mathbf{P}_{\mathbf{10}}^{\mathbf{RGD}}$ with: (1) 75 µM forward splint (Strand ID #6a) and 150 µM blocking strand (Strand ID #15a), and (2) 75 µM reverse splint strands (Strand ID #6b) and 150 µM blocking strand (Strand ID #15b). Concentrated DMEM was added to the precursors to reach 1x final concentration. The two precursor solutions were equilibrated at 4°C overnight to allow the blocking strands to quantitatively bind to the splint strands.

##### 2.15.2 Cell encapsulation

A cell suspension was first mixed with one DyNAtrix precursor (1) and cooled to 4°C. Subsequently, it was mixed with the second precursor (2) on ice to reach a final cell density of:

- MSCs: 1 x $10^6$ cells/mL
- MDCKII: 4 x $10^5$ cells/mL
- hiPSCs: 5 x $10^3$ aggregates/mL
- hTSCs: 4 x $10^5$ cells/mL

5-8 µL/well of the cell-DyNAtrix mixture were subjected onto 384-well plates or 6-well plates containing micro-well inserts (ibidi, Cat. 80409). The samples were incubated at 37°C for 20-40 minutes to trigger the heat-activated gelation. After gelation, the well was filled up with individual culture medium:

- MSCs: Same as 2D culture.
- MDCKII: Same as 2D culture.

- hiPSCs: mTeSR™1 supplemented with 8 µM ROCK inhibitor Y-27632 in the first 24 hours. On the next day, the ROCK-containing medium was replaced with fresh mTeSR™1.
- hTSCs: organoid medium [DMEM/F12 supplemented with 1x N2 supplement, 1x Glutamax, 1x B27 supplement without vitamine A, 100 µg/mL Primocin, 1.25 mM N-acetyl-L-cysteine, 1.5 µM CHIR99021, 50 ng/mL human EGF, 80 ng/mL human R-spondin-1, 100 ng/mL human FGF-2, human 50 ng/mL HGF, 0.5 µM A83-01, 2.5 µM Prostaglandin E2, and 2 µM Y27632].

When serum-containing medium was used, 100µg/mL of rabbit skeletal muscle actin was added to suppress nuclease activity. The embedded cells were cultured at 37°C in a humidified incubator equilibrated with 5% $CO_2$.

For Matrigel groups, cells were embedded in 50% Matrigel. The cell seeding density, gelation time, and culture medium are identical as in DyNAtrix samples.

#### 2.16 Fluorescence Live/Dead assay

##### 2.16.1 Mesenchymal stem cells (MSCs)

Cells were stained in medium containing 3 µM calcein-AM (PromoKine, Cat. # PK-CA707-80011) and 0.75 µM Draq7 (invitrogen, Cat. # D15106 ) for 30 minutes at 37°C, 5% $CO_2$. Confocal images were taken on a high-content spinning disk confocal microscope (Opera Phenix Plus High-Content Screening System, PerkinElmer) at 10x magnification. 3D images were analyzed by IMARIS (Oxford Instrument) software using spots wizard for cell counting. Cell viability was quantified by dividing the number of live cells by the sum of live and dead cells.

In the cell release test, cells were stained with 3 µM calcein-AM for 30 minutes. Subsequently, DNase I (2U/well) was added on top of the gels. The confocal images were acquired every 20 minutes (Opera Phenix Plus High-Content Screening System, PerkinElmer) at 5x magnification for 2 hours at 37°C, 5% $CO_2$.

##### 2.16.2 Other cell types

The DyNAtrix was degraded by 160U/ml DNase I in 250 µL medium at 37°C, 5% $CO_2$ before live/dead staining. Matrigel was degraded in 250 µL cell recovery solution (Corning, Cat. # 354253). After degradation, cells were washed with medium once and incubated in staining medium [3 µM calcein-AM, 0.75 µM Draq7, and 2 drops/mL NucBlue (invitrogen, Cat # R37605)] for 30 minutes at 37°C, 5% $CO_2$. The confocal images were acquired at 20x magnification on Andor Dragonfly Confocal Microscope System (Oxford instrument) or with spinning disk confocal microscope (Andor Revolution WD Borealis Mosaic) with Olympus silicone 30X/1.08 U Plan SApo objective with silicon oil immersion medium and 2µm Z-step.

#### 2.17 Bioprinting

MDCK II cells were encapsulated in 400 µL DyNAtrix (1 %(w/v) $\mathbf{P_5^{RGD}}$ + CCL-64 with blocking strands) following the protocol 2.15. The cell-laden hydrogel was transferred directly into a 1 ml Luer-Lock syringe (Omnifix®-F Solo, Cat. # 916700) with a 30-guage dispensing tip (Vieweg, Cat. # 500908). The syringe was centrifuged at 300 Xg for 3 min to remove air bubbles. The cell-laden gel was printed on a glass-bottom 6-well plate on a BioScaffolder BS5.1 (GeSiM) at a speed of 2 mm/s and an extrusion rate of 40 µm/s for 2 layers with 0.12 mm layer height. One hour after printing, the constructs were stained in 200 µL medium containing 3 µM calcein-AM, 0.75 µM Draq7, and 2 drops/mL NucBlue for 30 minutes. The images were acquired on Opera Phenix Plus High-Content Screening System at 10x magnification.

#### 2.18 Fluorescence immunostaining

##### 2.18.1 Human induced pluripotent stem cells (hiPSCs)

The cells were fixed with 2% paraformaldehyde (PFA) for 40 minutes at RT, permeabilized and blocked with 0.1%Triton X-100 in 2%BSA/PBS for 1 hour at 4°C. The cells were incubated with the primary antibodies [1:300 anti-Oct3/4 (BD Biosciences, Cat. # 611202)] in 2%BSA/PBS overnight at 4°C. Wash the samples with PBS 3 times, then add the secondary antibodies [1: 200 Alexa Fluor 488, 1: 200 Phalloidin ATTO 550, and 1: 10,000 Hoechst 33342] in 2%BSA/PBS overnight at 4°C. The confocal images were acquired with high-content spinning disk confocal microscope (Opera Phenix Plus High-Content Screening System, PerkinElmer).

##### 2.18.2 Human trophoblast stem cells (hTSCs)

hTSCs and placenta organoids were fixed with 2% PFA for 40 minutes at 4°C, permeabilized with 0.5% Tween-20/PBS for 30 mins, and blocked with 3% BSA (Sigma) + 0.1% Tween-20 for 1 hour. The cells were incubated with the primary antibodies in the blocking solution for 2 days at 4°C. After washing with PBS for 3 times, the cells were incubated with the secondary antibodies in the blocking solution for 2 days at 4°C. The following primary antibodies were used: anti-E-cadherin (Invitrogen, Cat. # 13-1900, 1:200), anti-Syndecan (Sigma, Cat. # HPA006185, 1:200), anti-GATA3 (R&D, Cat. # AF2605, 1:200), anti-ENDOU (Sigma, Cat. # HPA012388, 1:200) anti-GCM1 (Atlas, Cat. # HPA011343, 1: 200), and anti-TEAD4 (abcam, Cat. # ab58310, 1: 200). The following secondary antibodies were used: Alexa Fluor 488-, 594-, and 647-conjugated antibodies. Nuclei were stained with Hoechst (1:200). The confocal images were acquired with (Zeiss LSM 880 Airy inverted microscope and the Zeiss multi-immersion 63x/1.3 LCI Plan-Neofluar objective with Water immersion medium and 2µm z step.

#### 2.19 Passaging of placenta organoids

We followed and adapted a recent protocol established by the Turco Lab[10]. In brief, the gels were collected to an Eppendorf tube and degraded on day 7 of culture. We added 150 µL DMEM/F12 medium to Matrigel samples and pipetted them for 400 times through a small-bore tip to break up the gels. For DyNAtrix, we added 160U DNase I in 150 µL medium and incubated them for 40 minutes to degrade the gels. Matrigel and DyNAtrix were added with 1 mL DMEM/F12 and centrifuge at 600g for 6 min at RT. After removing the supernatant, all samples were added with 500 µL of pre-warmed Accutase and incubated for 5-6 min at 37°C. After incubation, the samples were washed once with 1 mL DMEM/F12. We suspended the organoids in 150 µL DMEM/F12 medium and pipetted for 80 times to break up the organoids. Finally, the samples were centrifuged at 600g for 6 min and re-suspended in 20 µL DMEM/F12 medium. The cell suspension was embedded in the gels at a final 1:4 splitting ratio.

#### 2.20 Statistical simulations of intra- vs. intermolecular crosslinks

Statistical simulations were carried out with a custom script that was written in Python 3. The script randomly selects $N_a$ crosslinker strands from a crosslinker library of size $N_x$. $N_a$ denotes the available number of anchor sites per polymer chain. The script quantifies the fraction of strands that would find a perfectly complementary binding partner on the same polymer chain, in order to determine the maximal percentage of intra-molecular crosslinks. Na and Nx are set to [1, 2, 4, 8, 16, 32, 64, 128, 256, 512], the script systematically scans all Na vs Nx combinations, and this scan is repeated 1000 times to create an average. The results are then exported in the form of a heatmap plot.

### 3 Supplementary Notes

#### 3.1 Benefit of a polymer backbone with high molecular weight

Unless there is significant entanglement (i.e., at high polymer concentration and high molecular weight), a polymer molecule can only contribute to the elasticity of the material, if it has more than 2 crosslinks that connect it to the network[11–13]. If the weight of the polymer backbone was only a few kilodalton, even a small number of DNA strands per backbone would represent the majority of the material by mass, since each DNA strand has a molecular weight of several kilodalton (~7kDa for each anchor strand in DyNAtrix). In fact, in most DNA-crosslinked polymer hydrogels the DNA crosslinkers represent the majority of the dry weight of the material[14,15]. In contrast, DyNAtrix has an ultra-high molecular weight backbone (~3 MDa; Tables S2 and S3), achieved by low initiator concentration and stringent exclusion of oxygen during polymerization (cf. section 2.1). In $\mathbf{P_1}$, $\mathbf{P_5}$, and $\mathbf{P_{10}}$ the calculated weight fraction of DNA is only 1.0%, 4.8% and 9%, respectively. The DNA handles can thus represent only a minor fraction of the material's dry weight while efficiently contributing to the material's elastic properties.

#### 3.2 Number of DNA anchor strands per polymer molecule

To estimate the number of anchor strands, we used the equation

$$M_P = M_A \cdot N_A + M_{AA} \cdot N_{AA} + M_{DNA} \cdot N_{DNA}$$

where $M_P$ is the number average molecular weight of the polymer chain (Table S2); $M_A$, $M_{AA}$, and $M_{DNA}$ are the molecular weights of acrylamide (AA), acrylate (A), and acrylamide-labeled DNA groups, respectively; $N_A$, $N_{AA}$, and $N_{DNA}$ are the numbers of corresponding monomers in the polymer chain.

As $M_A \cdot N_A << M_{AA} \cdot N_{AA}$ for all polymer derivatives, we used the simplified equation

$$M_P \approx M_{AA} \cdot N_{AA} + M_{DNA} \cdot N_{DNA}$$

The monomers ratio $r = N_{AA}/N_{DNA}$ in the synthesis is ~$7 \cdot 10^3$, $1.4 \cdot 10^3$, and $7 \cdot 10^2$ for $\mathbf{P_1}$, $\mathbf{P_5}$, and $\mathbf{P_{10}}$, respectively. Since ~75% of DNA anchor strands are both included in the polymer and accessible for binding (Figure S2), $r$ was corrected to be ~$1 \cdot 10^4$, $2 \cdot 10^3$, and $1 \cdot 10^3$ for $\mathbf{P_1}$, $\mathbf{P_5}$, and $\mathbf{P_{10}}$, respectively. Replacing $N_{AA}$ and rearranging the equation we obtain:

$$N_{DNA} \approx \frac{M_P}{r \cdot M_{AA} + M_{DNA}}$$

#### 3.3 Affine network model

According to the affine network model, the elastic modulus, G', is directly proportional to the density of effective (i.e. *inter*-molecular) crosslinks, $v_e$:[11]

$$G' = A \cdot v_e \cdot R \cdot T$$

where R is the ideal gas constant, T is the temperature in Kelvin degree, and A is a constant that equals to 1 for affine networks. The equation was used to derive $v_e$ values. Crosslinking efficiencies (CE) were calculated as CE = $v_e \;/\; v_{e,\max}$ , where $v_{e,\max}$ is the maximally possible crosslinker density at a given concentration.

#### 3.4 Design of heat-activated crosslinkers (HACs)

The addition of blocking strands (**B**) converts ordinary crosslinker splints (**S**) into HACs (see scheme below). **B** was designed to bind a large part of the crosslinker overlap domain (**c**/**c'**) to prevent premature crosslinking at low temperature. The length of the binding sequence (**b**/**b'**) was chosen to be 10 nt. This domain length was expected to be stable at 4°C, while at 37°C (close to its predicted melting temperature, $T_m$ ~ 40°C) **S** and **B** were expected to frequently dissociate and re-associate. No more than 2 consecutive bases were allowed to remain unpaired on ***c***, since longer single-stranded domains could potentially act as toeholds for a strand displacement pathway that would lead to premature crosslinking at low temperature. In order to provide additional thermodynamic driving force for the crosslinking, **b** was flanked by two 2-nt single-stranded overhangs ($\mathbf{f_1}$ and $\mathbf{f_2}$). Analogously, **b'** was flanked by two 2-nt domains ($\mathbf{f_1'}$ and $\mathbf{f_2'}$) that were complementary to the flanking domains in **b**. The purpose of the **f** domains was to add a thermodynamically favorable bias to the crosslinking reaction: upon their release from **S**, two complementary **B** strands can form a 14-nt dimer that is thermodynamically stable, thereby removing the blocking strands from the equilibrium. Specific blocking strand sequences are listed in Table S1.

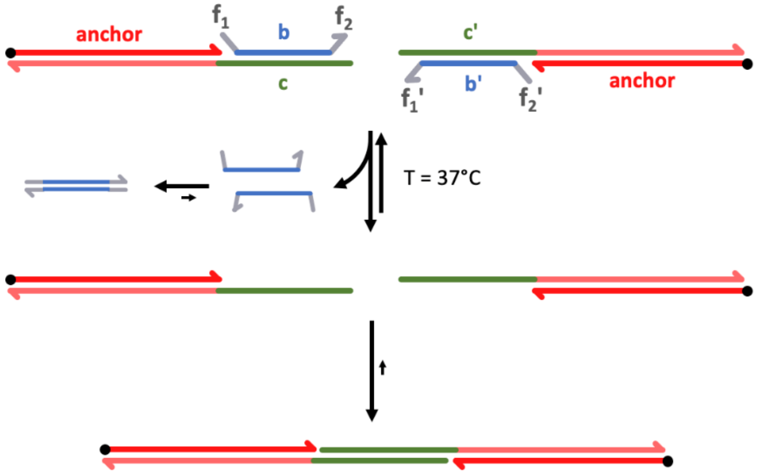

### 4 Supplementary Figures

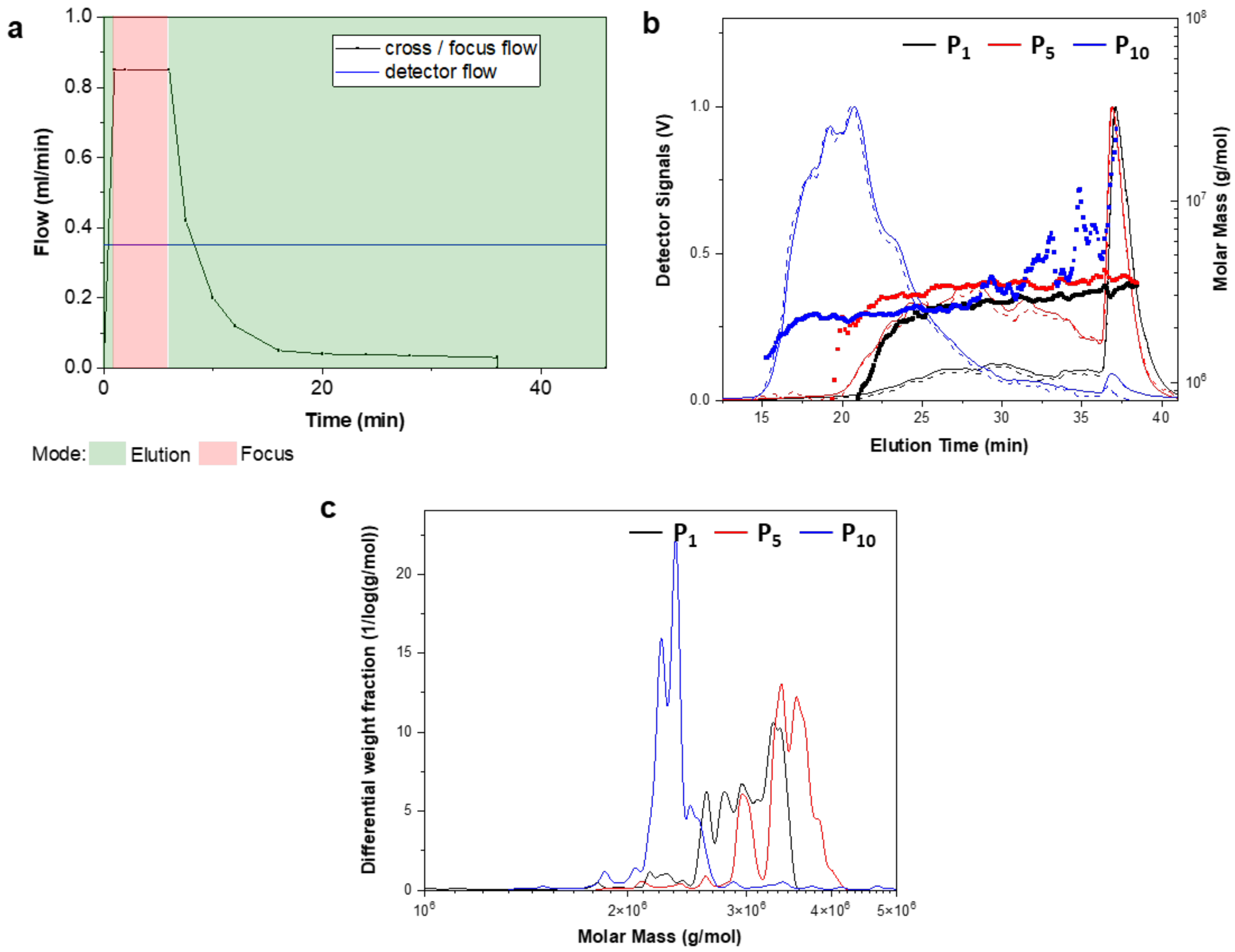


*Figure S 1. Asymmetrical flow field-flow fractionation with light scattering (AF4-LS) of **$P_1$, $P_5$, and $P_{10}$**. a) Optimized flow profile for AF4 separation, b) Fractograms, detector signals (solid line: multi-angle light scattering, dashed line: refractive index) and c) molar mass distributions.*

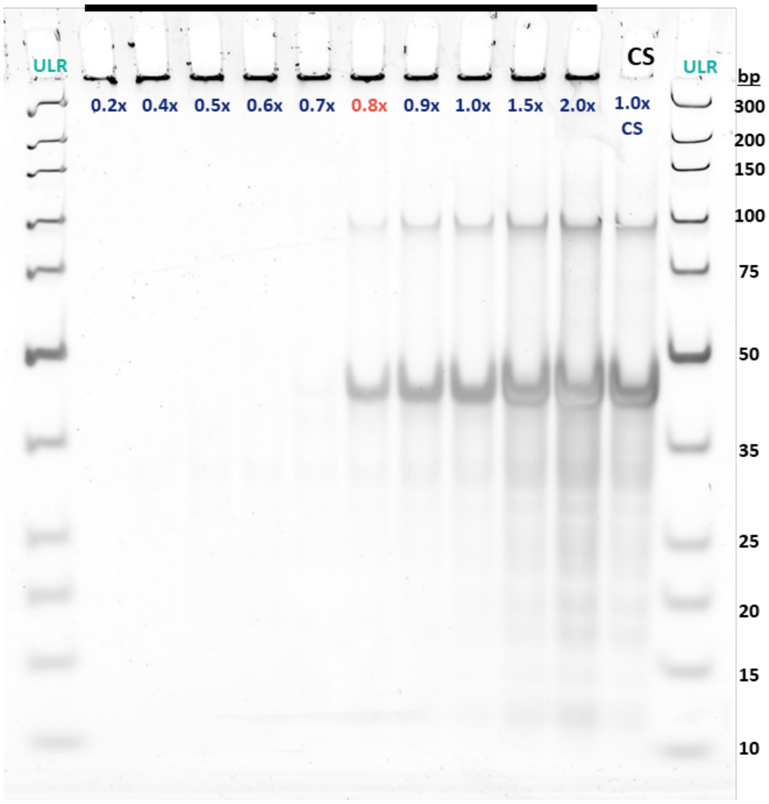


*Figure S 2. DNA binding efficiency test to quantify the number of accessible anchor strands per polymer chain. Taking **$P_5$** as an example, 0.4 % (w/v) polymer, which contains a maximum concentration of 2 µM anchor strands, was mixed with the complementary strand (CS) at ratios ranging from 0.2x to 2.0x. Complementary strands at concentrations below 0.8x did not appear as distinct bands on the gel, indicating that they were quantitatively bound to the polymer. 0.8x of complementary strands saturated the anchor strands and show clear bands of excessive CS on the gel. Thus, the concentration of available anchor strands was determined to be approximately 1.5 µM.*

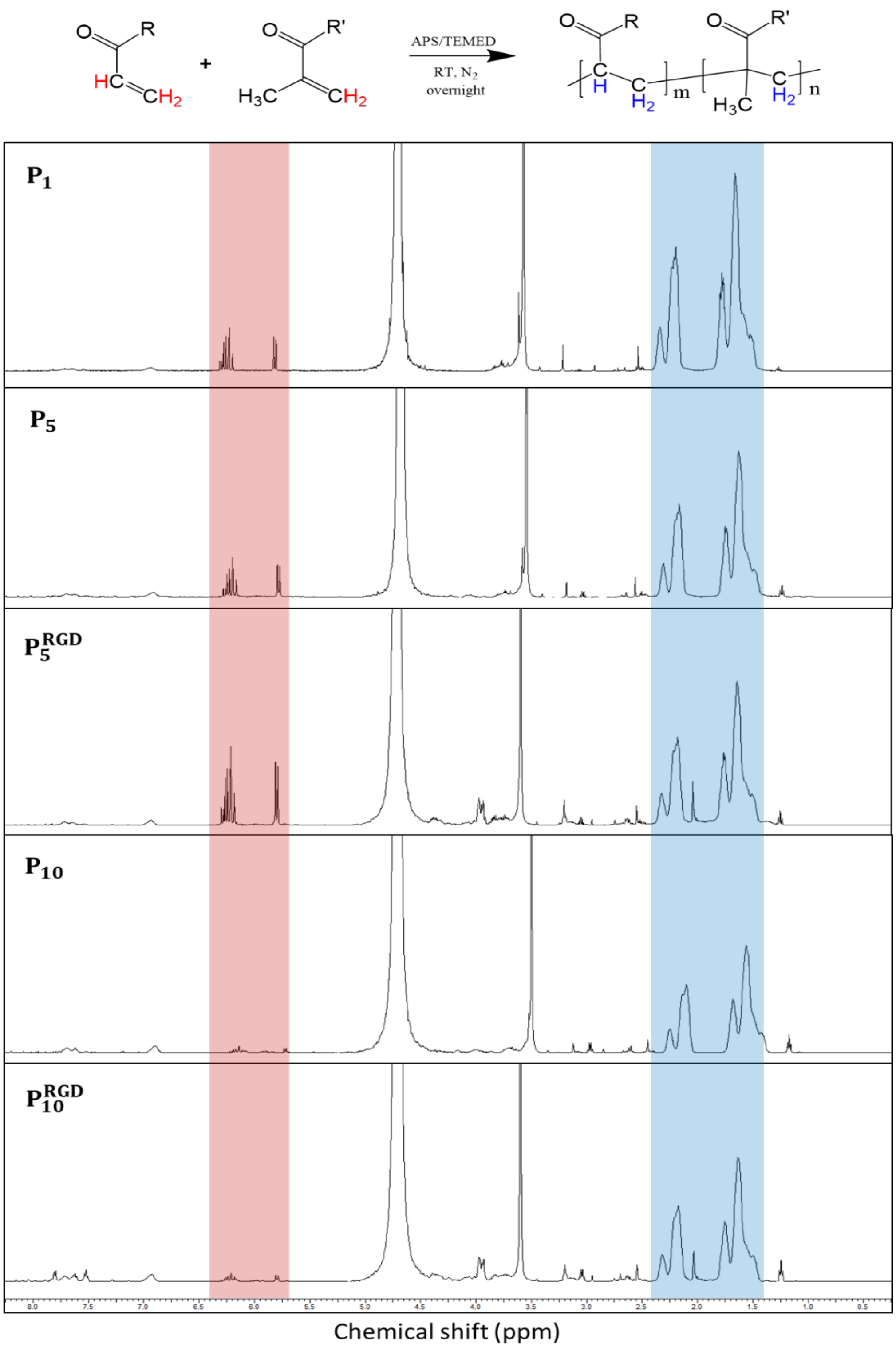


*Figure S 3. $^1$H-NMR spectra of **$P_1$**, **$P_5$**, **$P_5^{RGD}$**, **$P_{10}$**, and **$P_{10}^{RGD}$** in $D_2O$ after completed reaction and prior to methanol precipitation purification. The conversion of the reaction was calculated by quantifying free residual acrylamide monomer protons ($\delta$~5.7-6.4; red) vs. polymer backbone protons ($\delta$~1.4-2.4; blue). Conversions are 96%, 94%, 90%, 99%, and 98% for **$P_1$**, **$P_5$**, **$P_5^{RGD}$**, **$P_{10}$**, and **$P_{10}^{RGD}$**, respectively. R = -$NH_2$ , -OH or -NH-peptides; R' = -NH-oligonucleotides.*

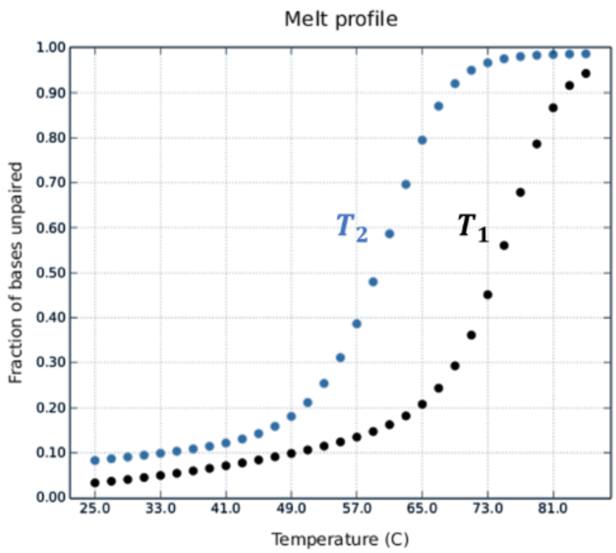


*Figure S 4. Predicted melting profile of the 22-nt anchor domain (black) and a 14-nt overlap domain (blue) of dual splint crosslinkers. Splints are predicted bind to anchor strands at a higher melting temperature ($T_1$) and then pair to a complementary splint at a lower melting temperature ($T_2$). The melting profiles were simulated in NUPACK*[16].

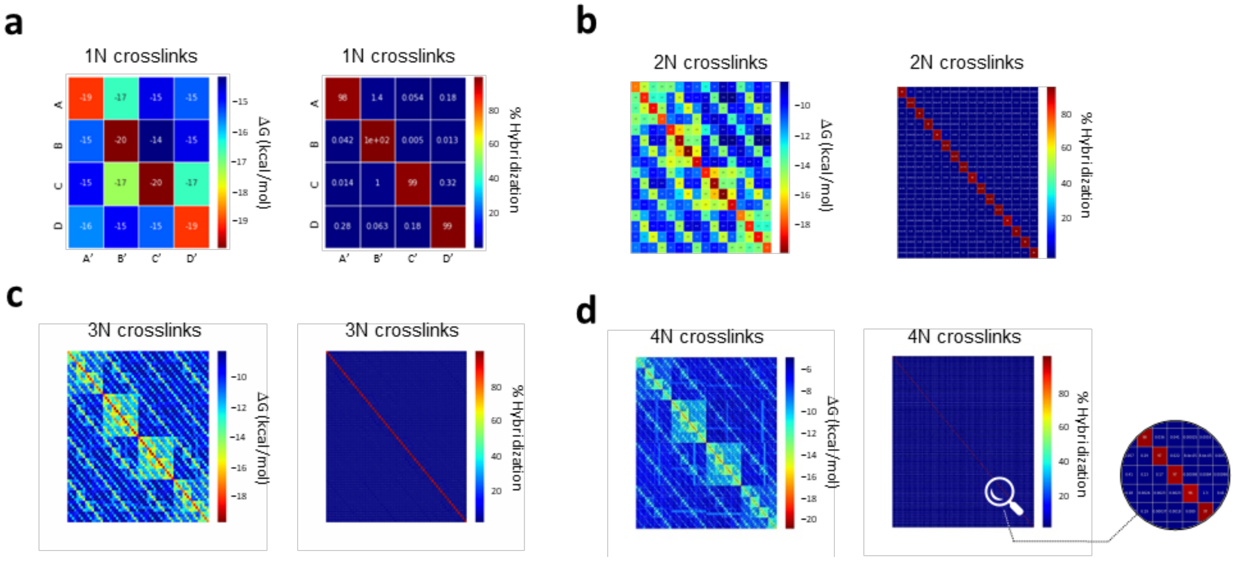


*Figure S5. Predicted minimum free energy (left) and the normalized Boltzmann distribution (right) of the combinatorial crosslinker libraries with 1 – 4 ambiguous bases. The simulation was carried out using NUPACK*[16]*. High resolution images are provided as Supplementary Files 1-10.*

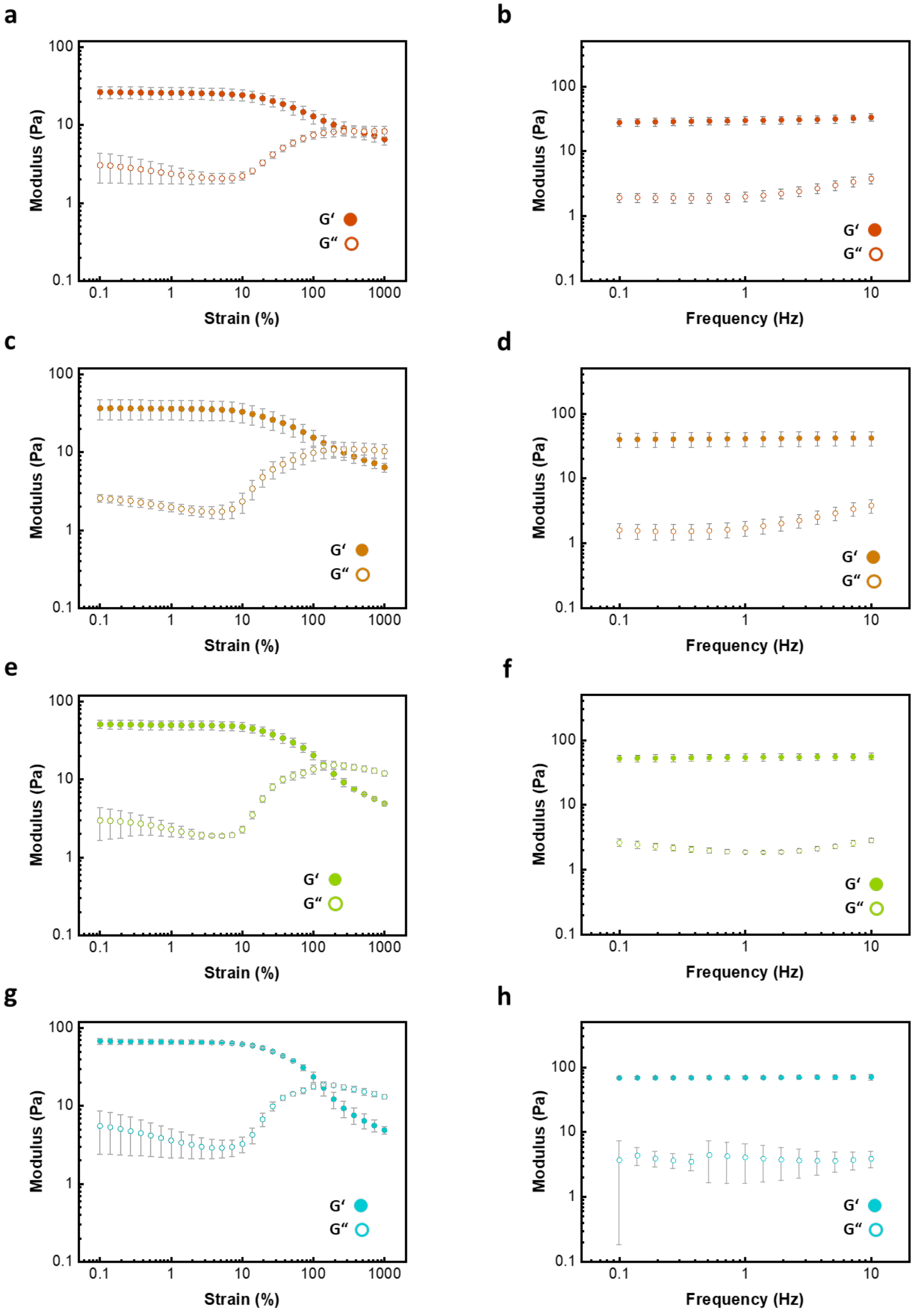
a
b
c
d
e
f
g
h
Modulus (Pa)
Strain (%)
Frequency (Hz)
G‘
G“
0.1
1
10
100
1000

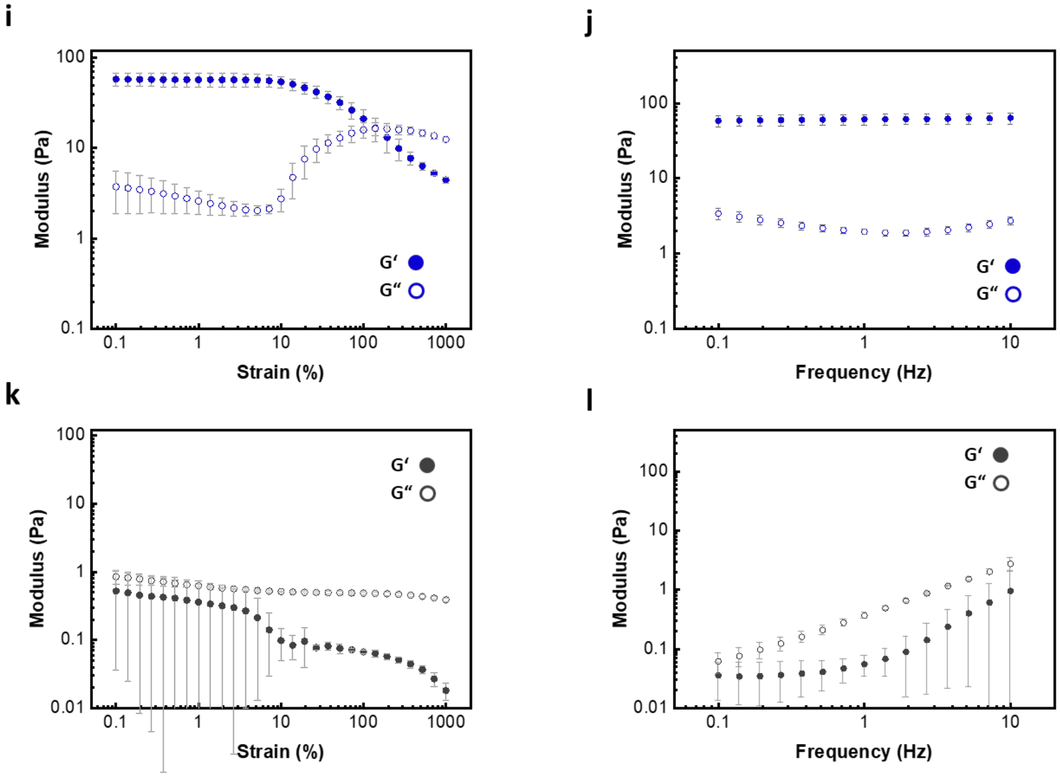


*Figure S 6. Amplitude sweeps (left) and frequency sweeps (right) of 1% (w/v)* ***$P_5$*** *hydrogels crosslinked by (a,b) CCL-1; (c,d) CCL-4; (e,f) CCL-16; (g,h) CCL-64; (i,j) CCL-256; or (k,j) without CCL. Error bars indicate the standard deviation from 3 independent repeat experiments. All samples were measured at 37℃.*

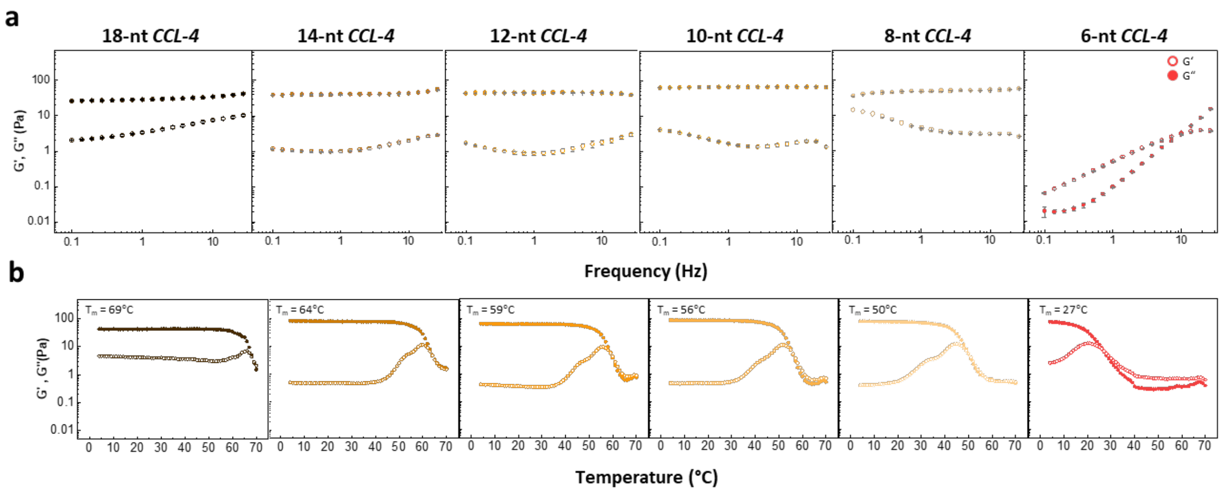


*Figure S 7. (a) Frequency sweeps (0.1–30 Hz) of SRC-crosslinked* ***$P_5$*** *(1% (w/v)) at 37°C. SRCs represent CCL-4 with overlap domains ranging from 6 to 18 nt. All gels exhibit gel-like mechanical properties (G’>>G’’), except for the 6-nt CCL-4, which is solid-like at 37°C only at high frequencies (>10 Hz). Error bars: mean±S.D. from one hysteresis measurement. (b) Temperature-dependent curves (4–70°C) of the respective samples in (a). Below their melting temperature, all gels exhibit a plateau region with comparable storage moduli. Melting temperatures, $T_m$, defined as the crossover temperature for G’ and G’’, are significantly above 37°C for SRCs between 8 and 18 nt.*

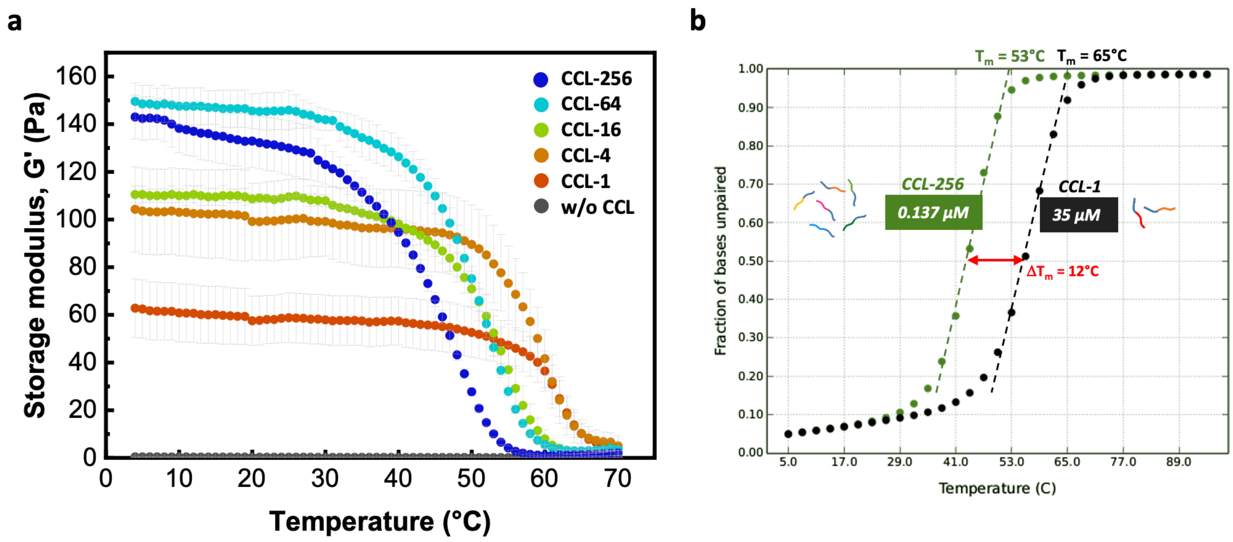


*Figure S 8. (a) Temperature sweep of 1% (w/v)* ***$P_5$*** *hydrogels crosslinked by different CCLs. The samples were measured at 10% strain and 1.6Hz in an oscillatory rheometer. Error bars indicate the standard deviation from 3 independent repeats. (b) NUPACK calculation for predicting the melting temperature ($T_m$) of different splint concentrations corresponding to the actual concentration of distinct splint pairs in CCL-1 (35 µM) and CCL-256 (0.137 µM).*

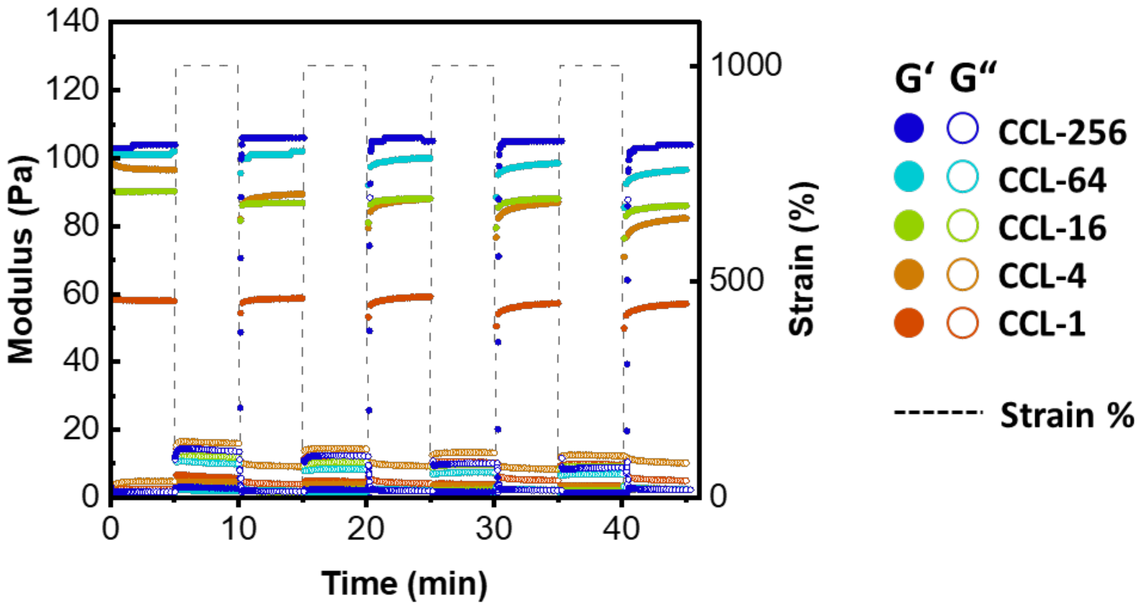


*Figure S 9. Rapid self-healing tests of 1% (w/v)* ***$P_5$*** *hydrogels crosslinked by different CCLs. The samples were sheared at 1000% (disruption) and 10% (recovery) strain at 1.6Hz, 37℃ for 5 repetitive cycles.*

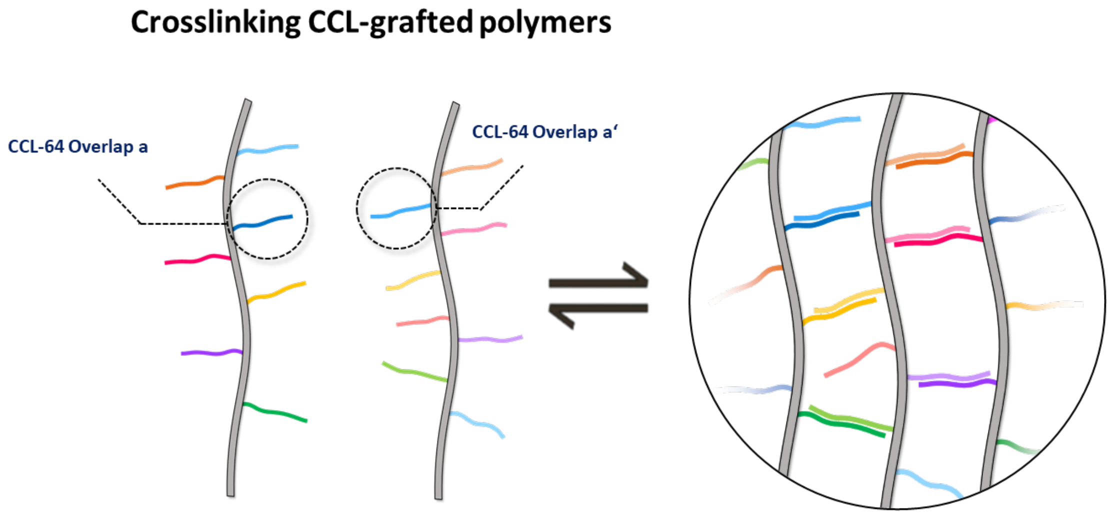


*Figure S 10. The scheme of crosslinking two complementary CCL-grafted polymers. To reduce DNA content, an overlap domain can be directly grafted onto the polymer backbone. When the two polymers are well mixed and thermally annealed they form stable supramolecular networks.*

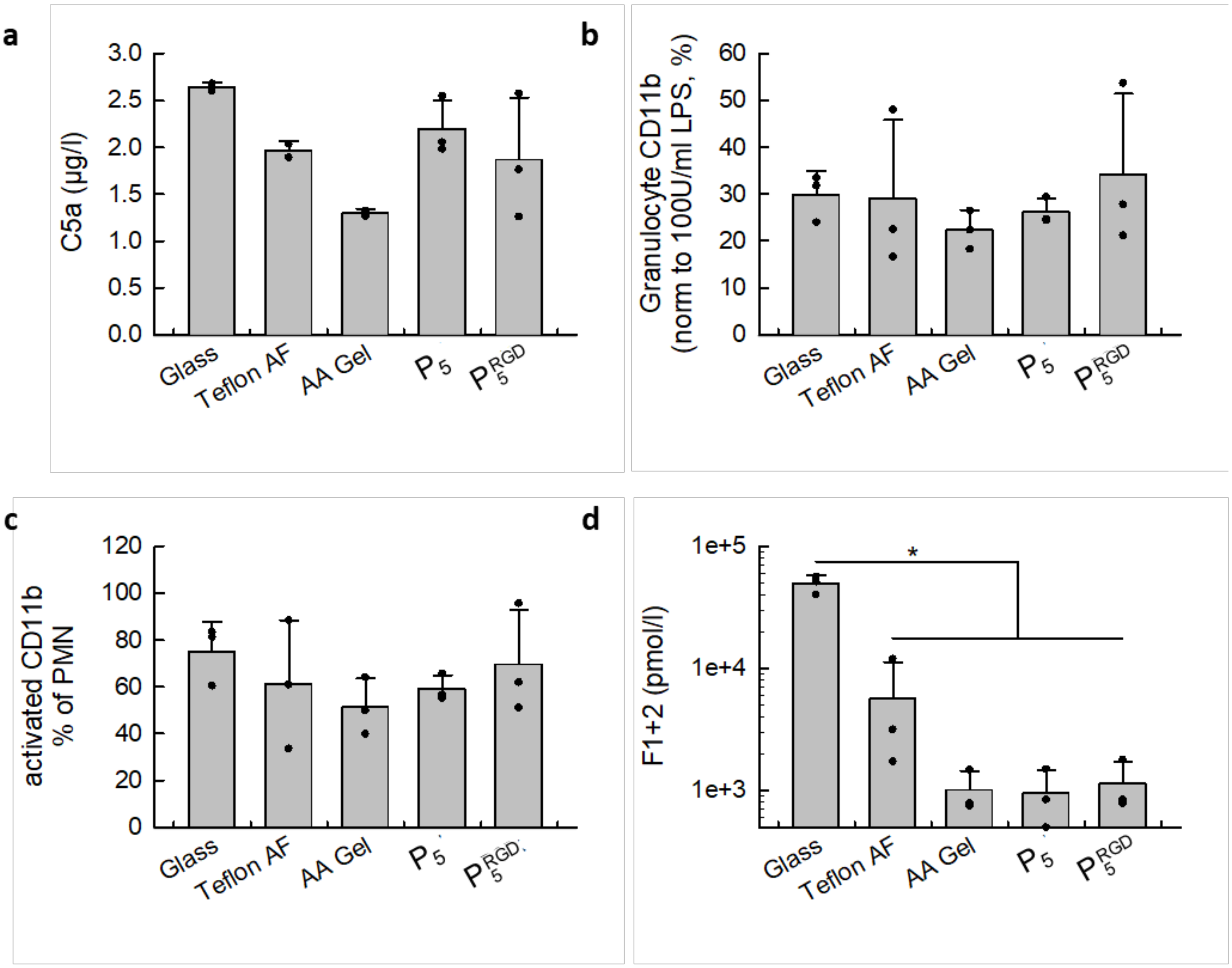


*Figure S 11. Innate immune response and hemocompatibility tests. 1% (w/v)* $\mathbf{P_5}$ *and* $\mathbf{P_5^{RGD}}$ *hydrogels crosslinked with CCL-64 were incubated in whole human blood. a) Markers for complement activation (C5a), b) granulocytes CD11b, c) activated CD11b as fraction of granulocytes, and d) hemostasis marker prothrombin fragment 1+2 (F1+2). The asterisk* indicates p<0.05 in one-way analysis of variance (ANOVA) on ranks with Student-Newman-Keuls post-hoc test.*

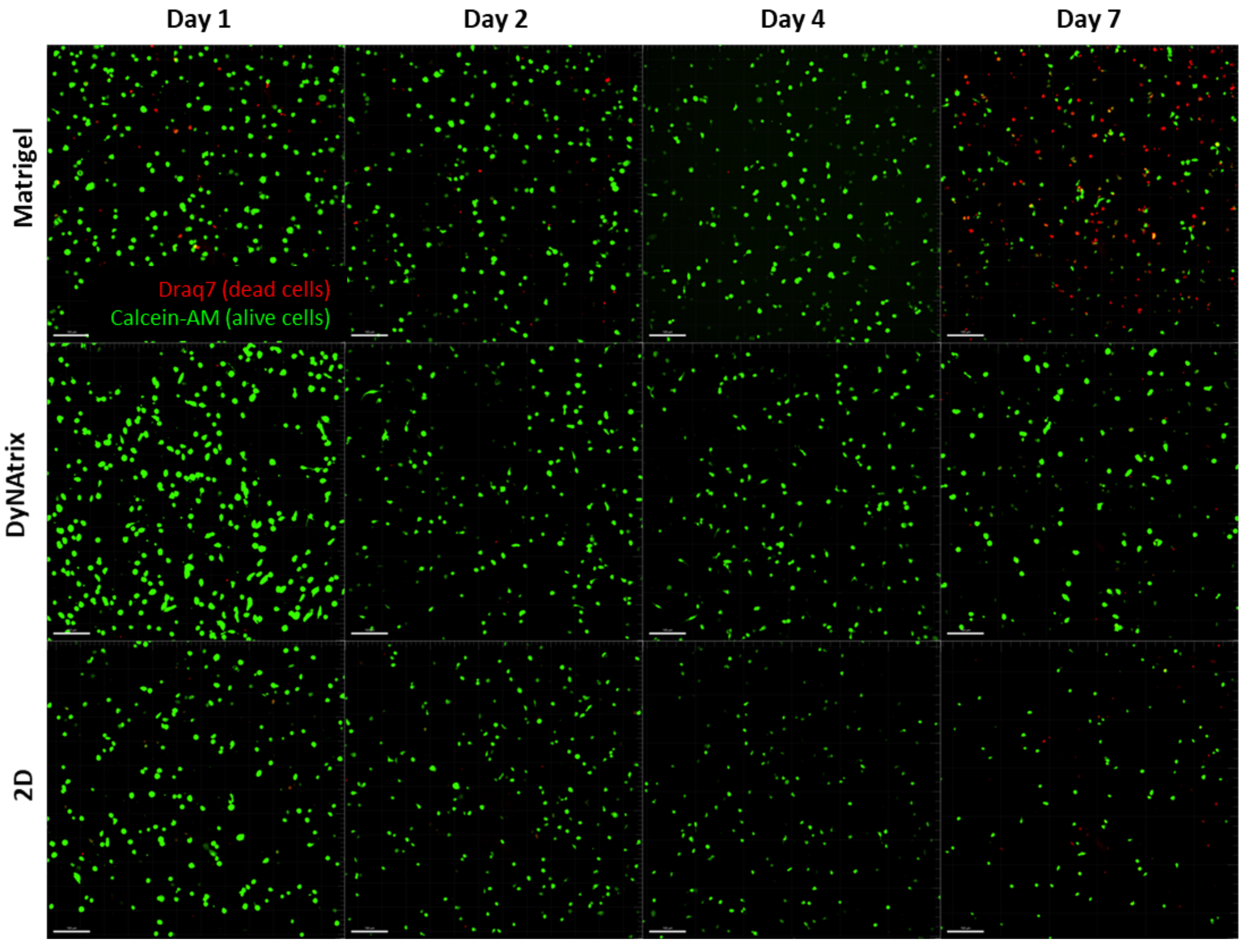


*Figure S 12. hMSCs viability tests with calcein-AM (live cells) and Draq7 (dead cells). The cells were cultured in Matrigel, DyNAtrix (1% (w/v)* $\boldsymbol{P_5^{RGD}}$ *+ CCL-64), and a 2D cell culture dish. Variations in cell density between days 1 and 2 are due to initial swelling of the gels. The viability is calculated using the ratio of total live cell divided by the sum of live and dead cells.*

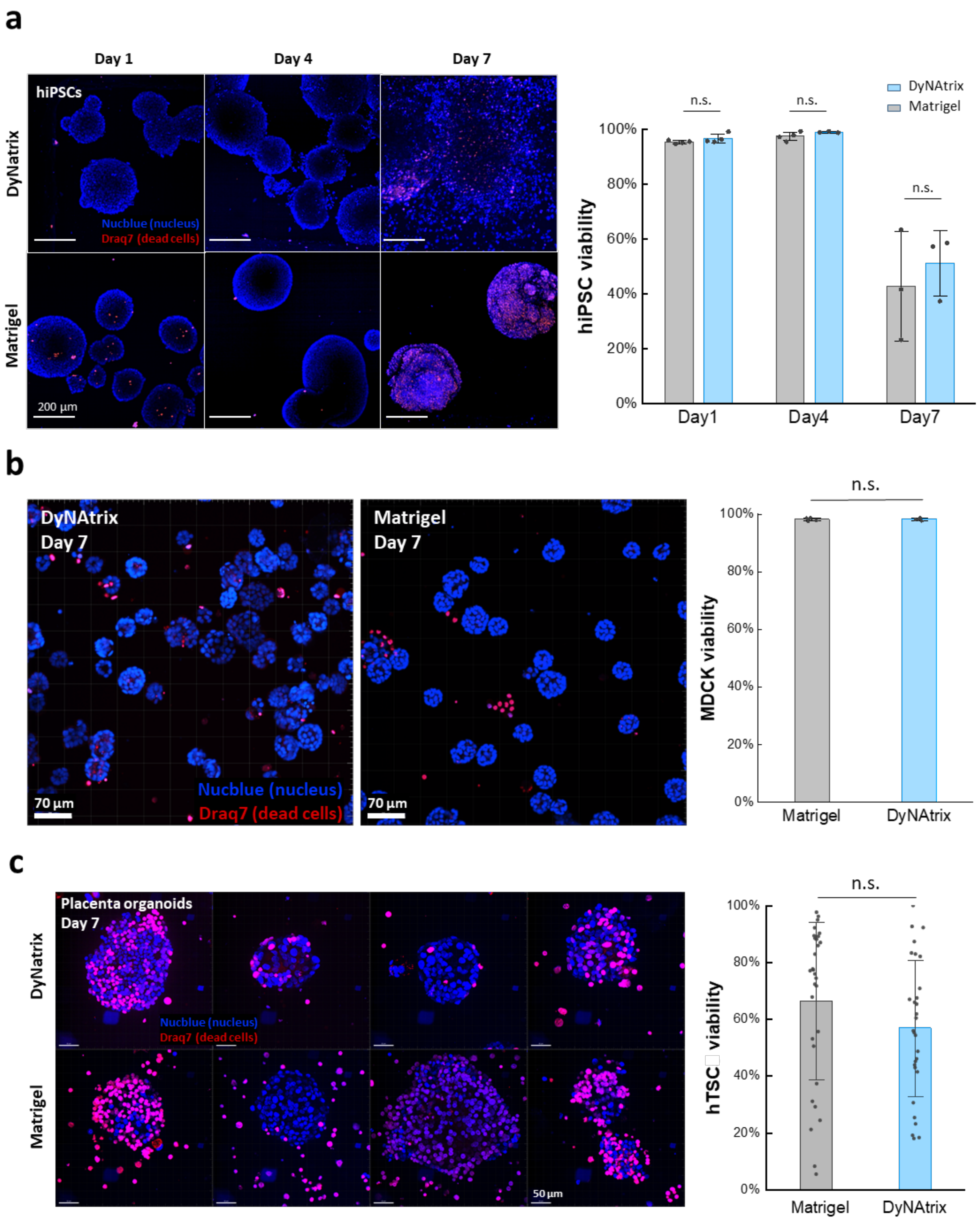


*Figure S 13. Viability tests of hiPSC (a), MDCK cells (b), and placenta organoids (c) stained with Draq7 (dead cells) and NucBlue (nuclei). a) hiPSCs were cultured in DyNAtrix (1% (w/v)* $\mathbf{P_5}^{RGD}$ *+ CCL-64) and Matrigel. Viability was high (>97%) for DyNAtrix and Matrigel on days 1 and 4. On day 7 increased cell death was observed in both matrices, likely because the size and density of the cysts became too large for sufficient nutrient uptake. Statistical analysis was evaluated with a two-way ANOVA test followed by*

*Tukey post-hoc analysis. b) MDCK cells were cultured in DyNAtrix (1% (w/v) $\mathbf{P_{10}^{RGD}}$ + CCL-64) and Matrigel. Statistical analysis was evaluated with a two-sample t-test. c) hTSCs were cultured in DyNAtrix (1% (w/v) $\mathbf{P_{10}^{RGD}}$ + CCL-64) vs. Matrigel and grown for 7 days into placenta organoids. Statistical analysis was evaluated with a two-sample t-test. The viability was calculated using the ratio of total dead cell divided by the total number of nuclei. Error bars indicate the standard deviation from n replicates (hiPSC: n = 3-4; MDCK: n = 5; hTSC: n= 30). n.s. = not significant (p>0.05).*

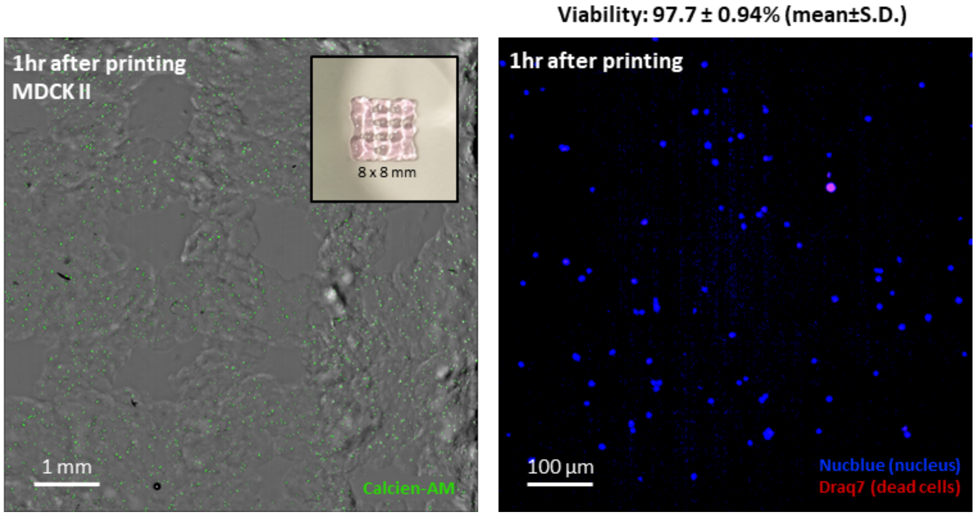


*Figure S 14. Viability test with MDCK cells after extrusion printing. The cells were embedded in DyNAtrix (1% (w/v) $\mathbf{P_5^{RGD}}$ + CCL-64), and subsequently the cell-laden gels were extruded through a 1 mL syringe with a 30G nozzle on a BioScaffolder (GeSiM). 1 hour after printing, the cells were stained with Calcein-Am (live), Draq7 (dead), and NucBlue (nuclei). The viability was calculated using the ratio of total dead cell divided by the total number of nuclei in 4 structures that had been printed under identical conditions.*

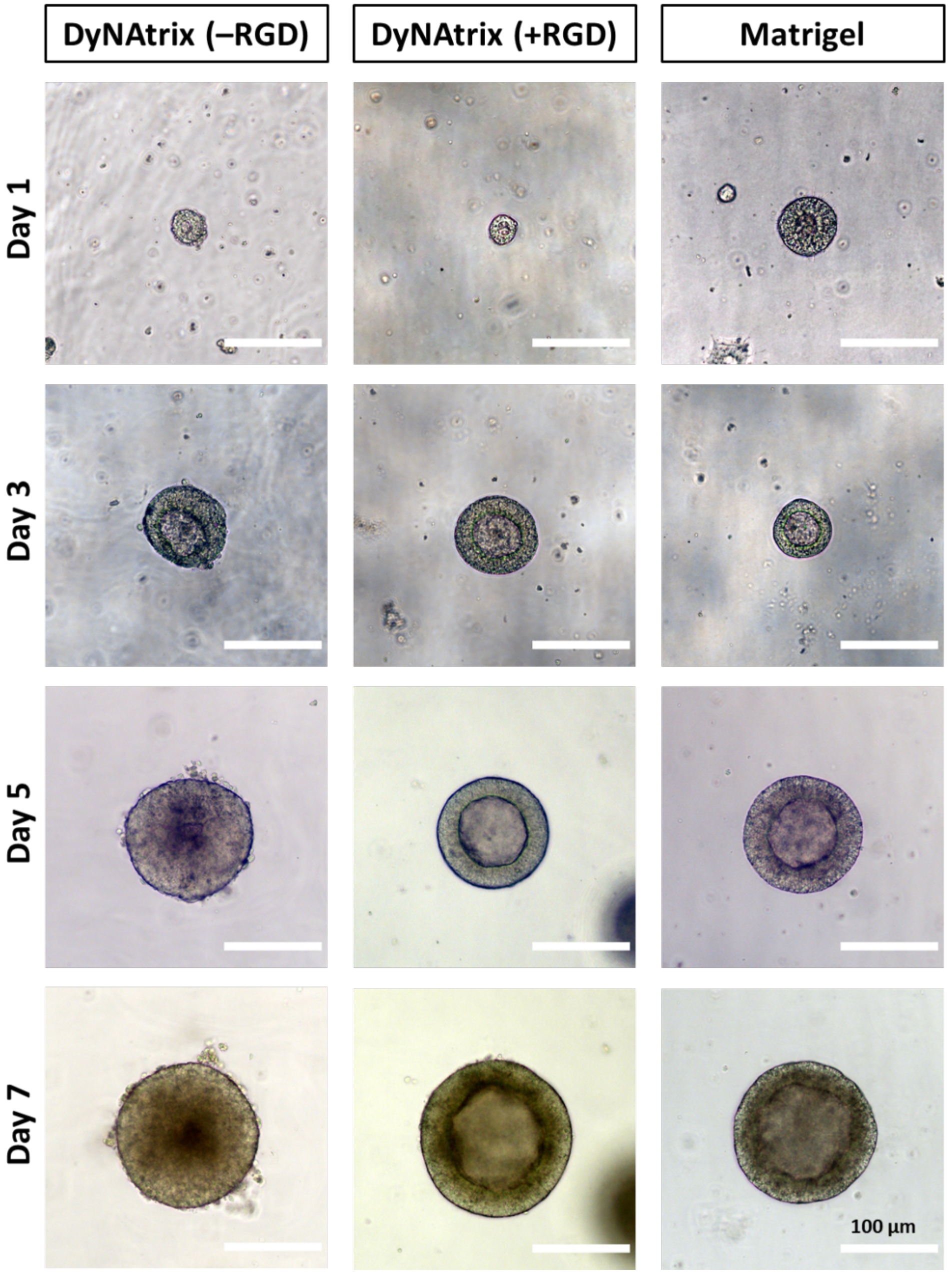


*Figure S 15. Representative brightfield images of hiPSC cysts cultured in DyNAtrix [-RGD] (1% (w/v)* $\boldsymbol{P_5}$ *+ CCL64) and DyNAtrix [+RGD] (1% (w/v)* $\boldsymbol{P_5^{RGD}}$ *+ CCL-64) vs. Matrigel on day 1, 3, 5, and 7.*

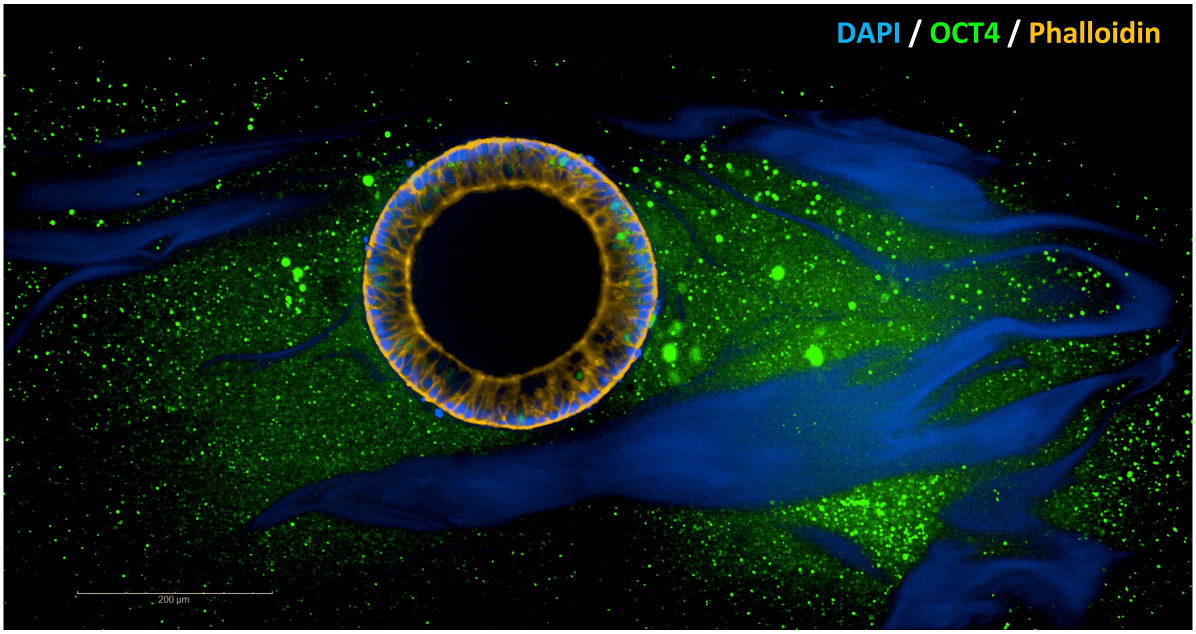


*Figure S 16. Confocal microscope image of a hiPSC cyst stained inside DyNAtrix [+RGD] (1% (w/v) $\mathbf{P_{10}^{RGD}}$ + CCL-64) after culture for 7 days. The cells were stained with pluripotency marker OCT4 (green). F-actin was stained with Phalloidin (yellow). The cell nuclei were stained with DAPI (blue). The DAPI signal arising from extracellular DNA crosslinkers can be detectable, resulting in blue schlieren. The differences in DAPI signal are enhanced by background subtraction.*

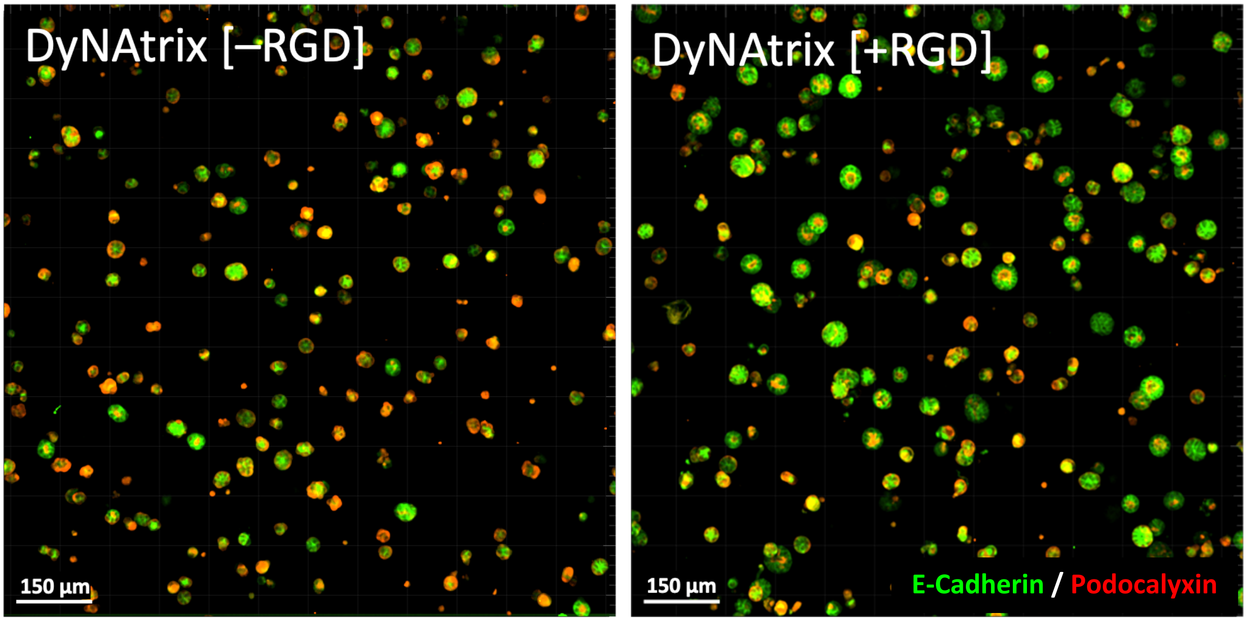


*Figure S 17. Culture of MDCK cysts in DyNAtrix with 1% (w/v) $\mathbf{P_5}$ vs. $\mathbf{P_5^{RGD}}$ backbone crosslinked with CCL-64. A serum-free medium (UltraMDCK™) was used. Images were taken on day 5 of the culture.*

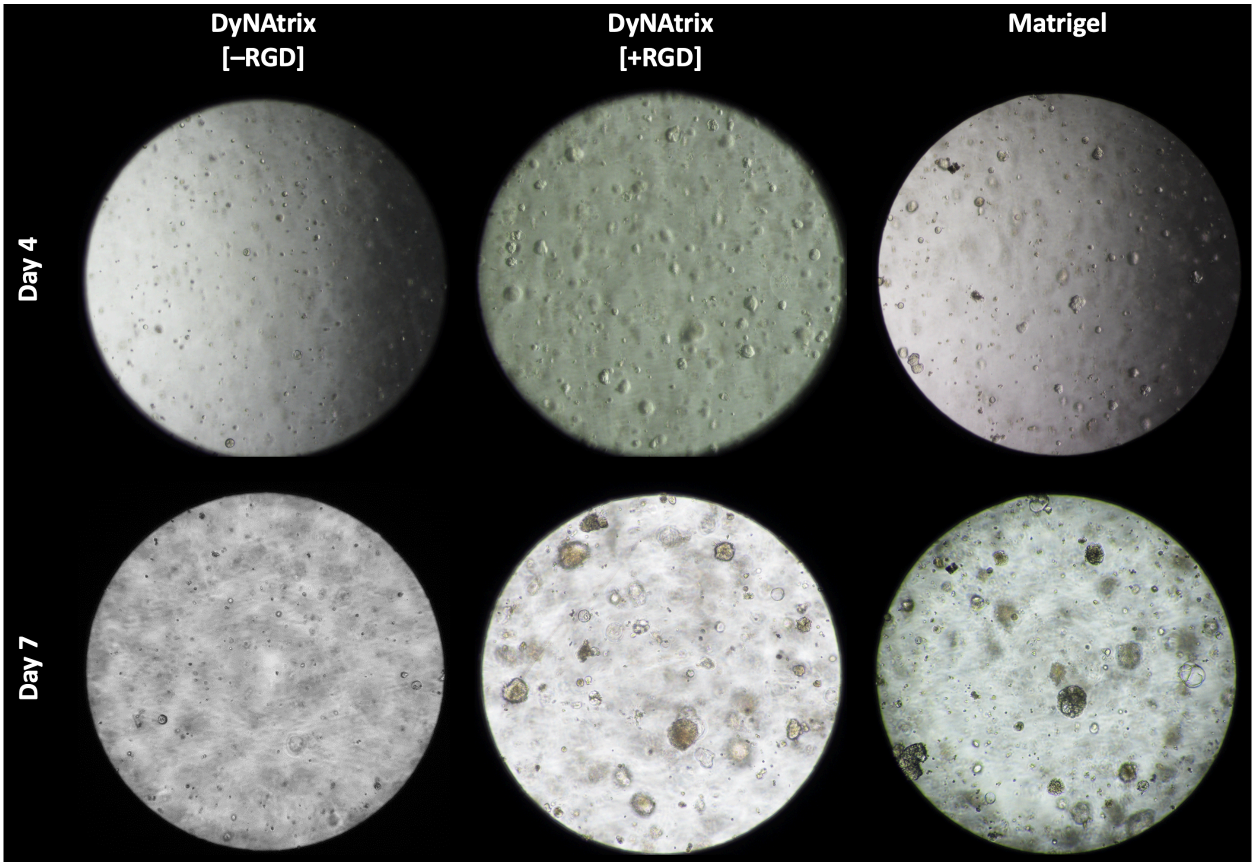


*Figure S 18. Brightfield microscopy images of placenta organoids grown in DyNAtrix [+/–RGD] vs Matrigel (day 4 and 7).*

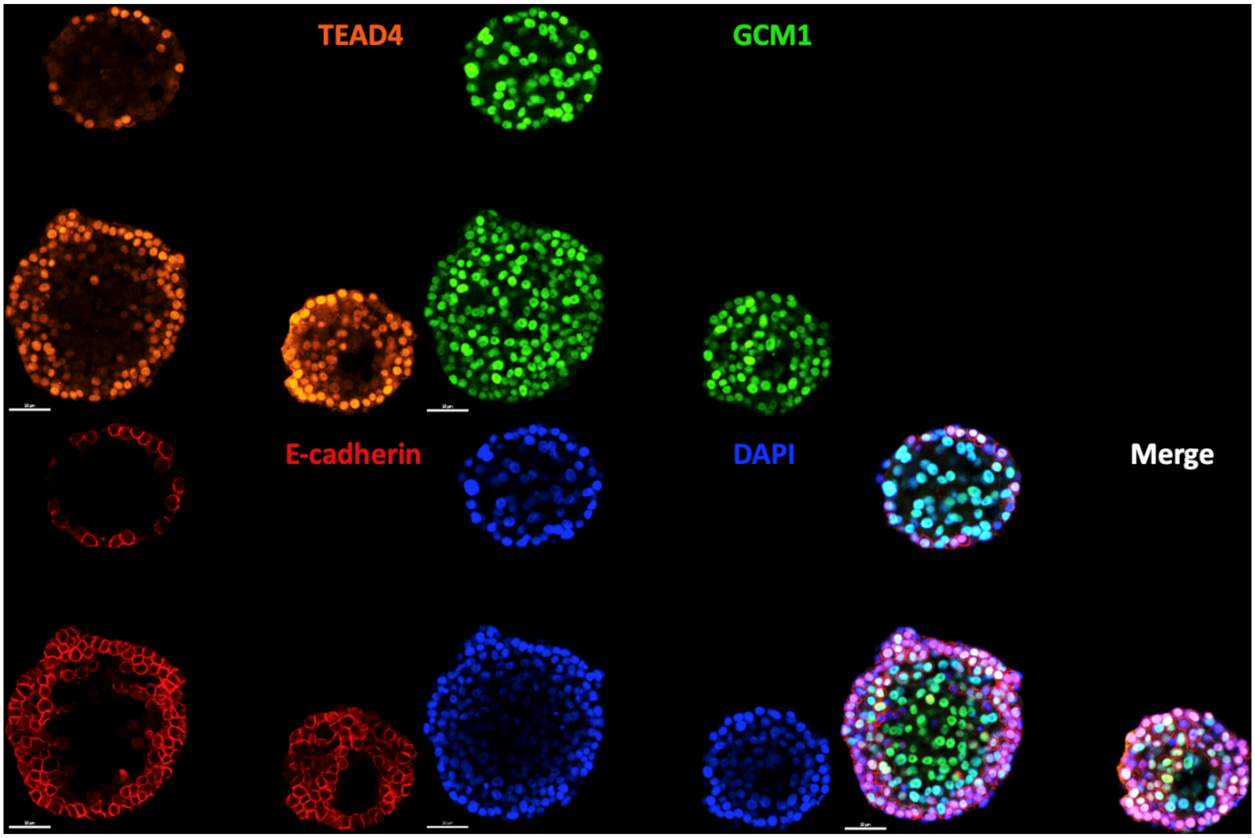


*Figure S 19. Confocal images of placenta organoids grown in DyNAtrix (1% (w/v)* $\mathbf{P}_{10}^{RGD}$ *+ CCL-64) on day 7 and stained for TEAD4, GCM1, E-Cadherin, and nuclei (DAPI). Scale bar: 50 µm.*

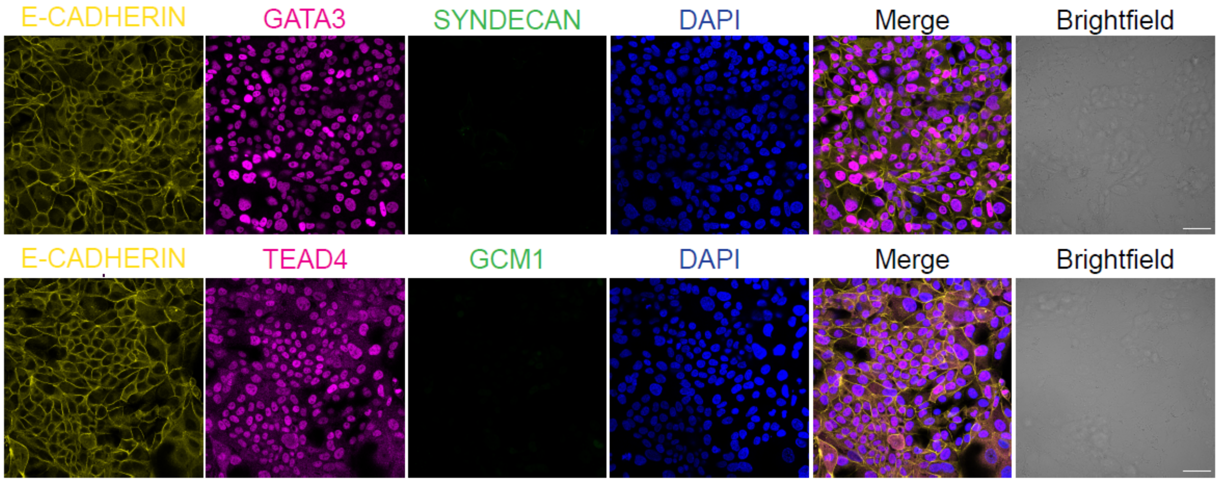


*Figure S 20. Immunostaining of 2D cultured trophoblast cells on day 7. The cells were stained for GATA3, TEAD4, SYDECAN, GCM1 and E-Cadherin. The monolayer cells showed low expression of differentiation markers GCM1 and Syndecan, but strong expression of stemness markers TEAD4 and GATA3. Scale bar = 50 µm.*

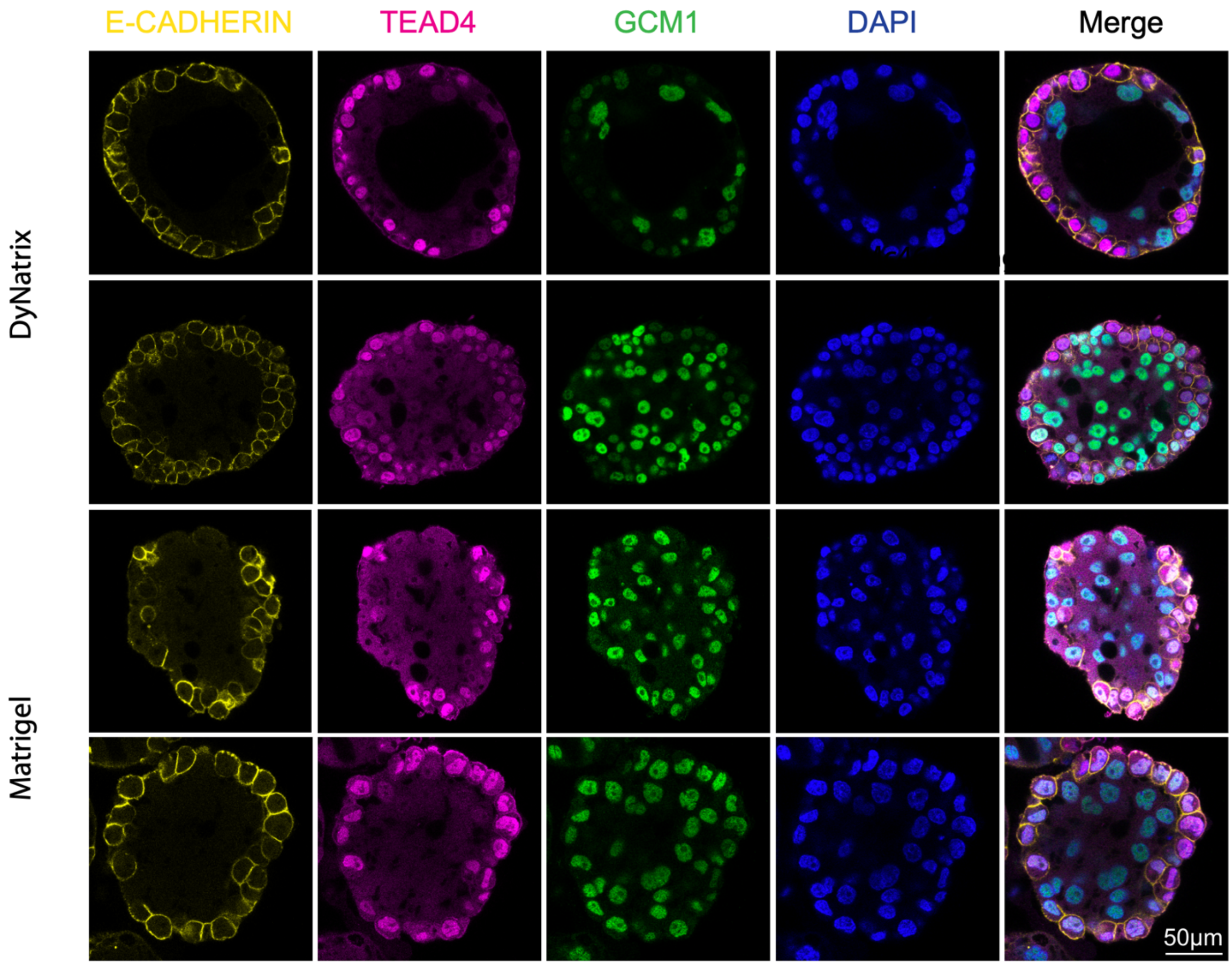


*Figure S 21. Representative confocal images of several placenta organoids grown in DyNAtrix (1% (w/v)* $\boldsymbol{P_{10}^{RGD}}$ *+ CCL-64) vs. Matrigel (day 7). Organoids were 100–200 µm in diameter and showed expression of E-Cadherin, TEAD4 and GCM1. Scale bar: 50 µm.*

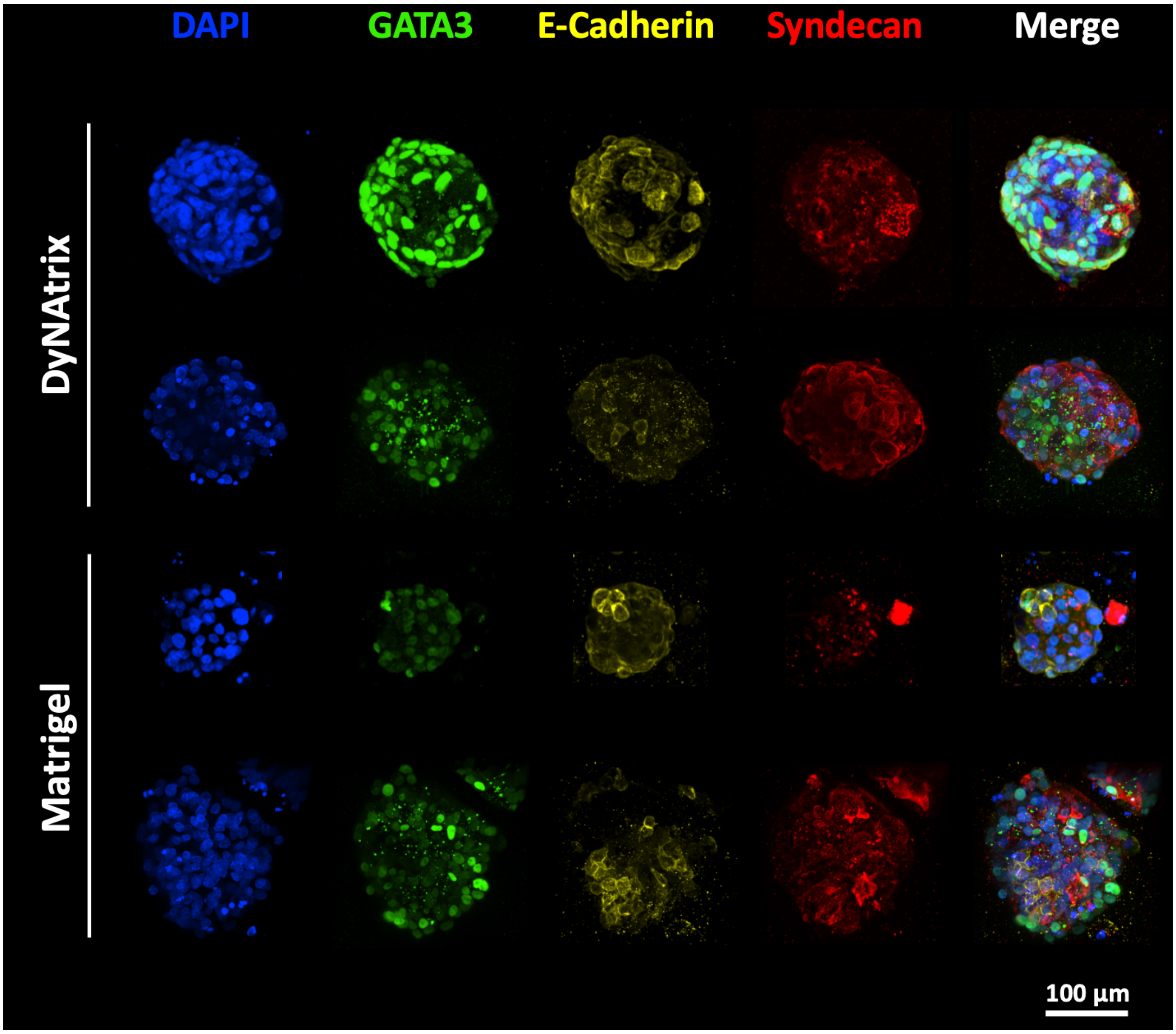


*Figure S 22. Representative confocal images of several placenta organoids grown in DyNAtrix (1% (w/v)* $\mathbf{P_5^{RGD}}$ *+ CCL-64) vs. Matrigel (day 7). Organoids were 100–200 µm in diameter and showed expression of GATA3, E-Cadherin and Syndecan.*

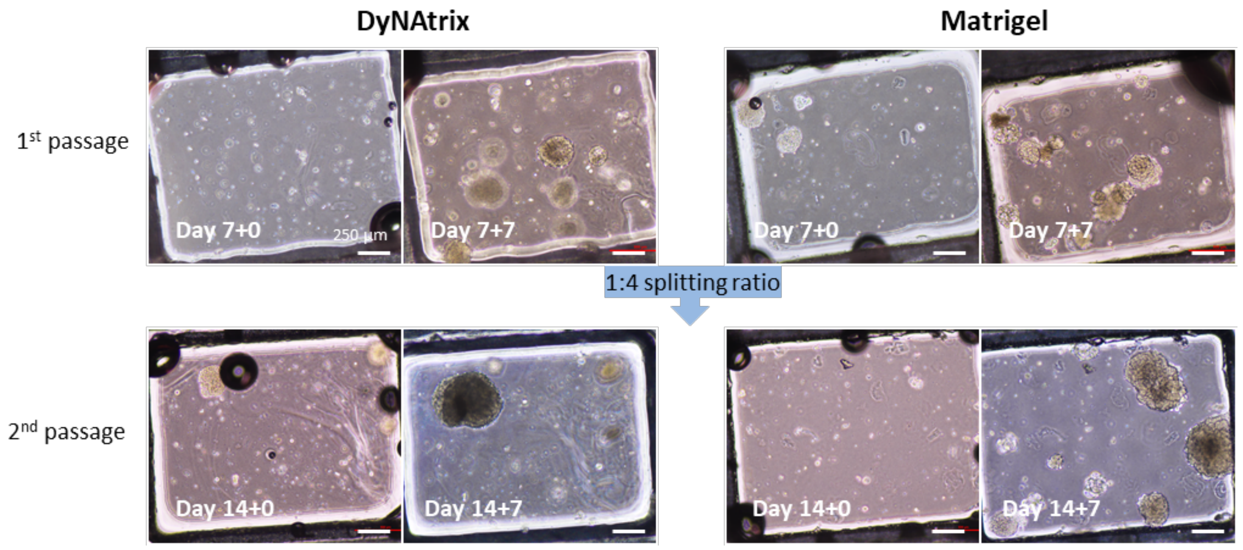


*Figure S 23. Long-term placenta organoid culture for up to 21 days in DyNAtrix (1% (w/v) $\mathbf{P_{10}^{RGD}}$ + CCL-64) vs. Matrigel. Brightfield images show organoids that were passaged twice after an initial 3D culture for 7 days. The passaging workflow for DyNAtrix was adapted from a protocol that had been previously established for Matrigel.*[10] *Scale bars: 250 µm.*

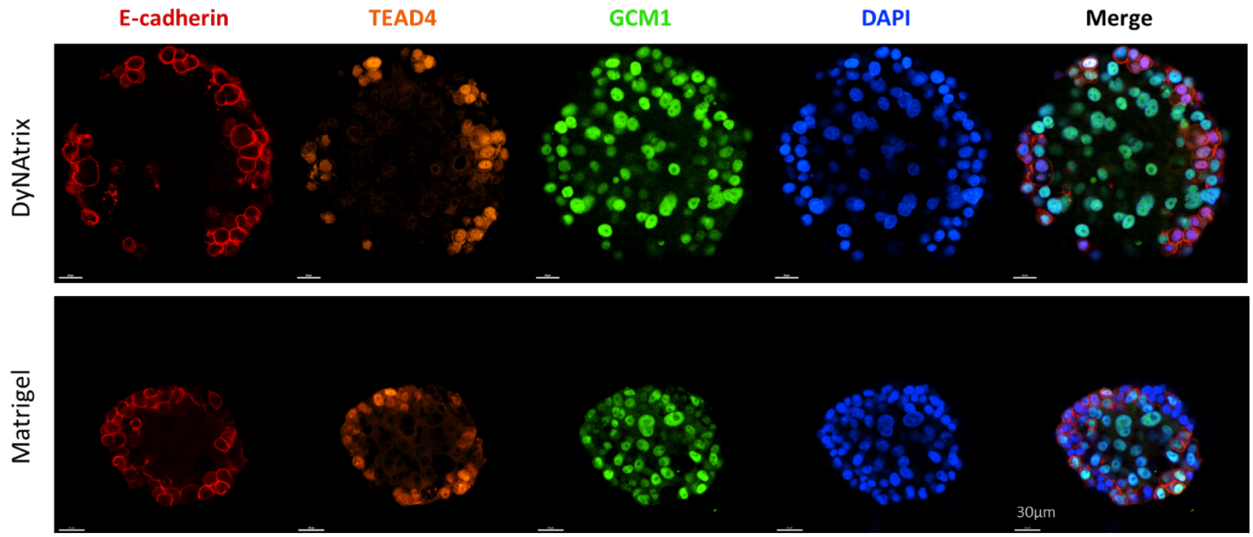


*Figure S 24. Confocal images of 2nd passaged placenta organoids in DyNAtrix (1% (w/v) $\mathbf{P_{10}^{RGD}}$ + CCL-64) vs. Matrigel (day 14+7). The organoids showed expression of E-Cadherin, TEAD4 and GCM1, similar to the one before passaging. Scale bar: 30 µm.*

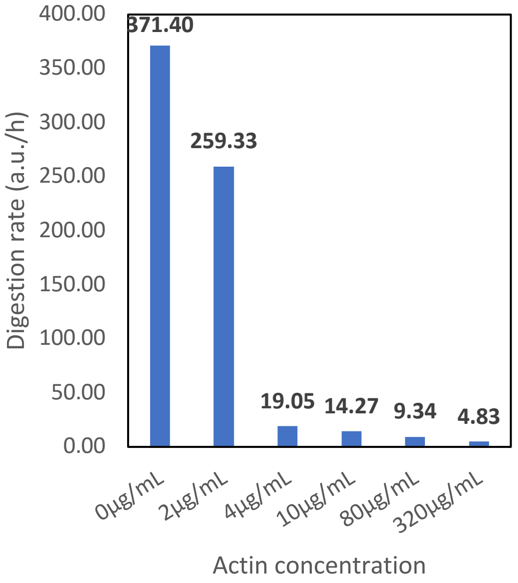


*Figure S 25. The digestion rate of FRET-paired DNA in DNase-containing medium supplemented with different concentration of actin. The rate is determined by the slope of the linear region of the regression line in the beginning of the fluorescence curve ($R^2$ > 0.98).*

#### 5 Supplementary Tables

**Table S 1.** List of the DNA sequences used in this study. Red: adaptor site complementary to anchor strand; blue: overlap domain; magenta: ambiguous N base (A,T,C, or G). 5Acryd: Acrydite; IAbRQSp: Iowa black RQ-Sp; Cy5: cyanine 5; 6-FAM: 6-carboxyfluorescein.

| # | Anchor strand | Length [nt] |
|---|---|---|
| 1 | /5Acryd/GACGGCTCATAAGGCTCTAATC | 22 |

| # | Single-splint crosslinker library | Length [nt] |
|---|---|---|
| 2 | GATTAGAGCCTTATGAGCCGTCGATTAGAGCCTTATGAGCCGTC | 44 |

| # | 1-splint library (*CCL-1*) | Length [nt] |
|---|---|---|
| 3a | TTAGTCAGTGTCCCGATTAGAGCCTTATGAGCCGTC | 36 |
| 3b | GGGACACTGACTAAGATTAGAGCCTTATGAGCCGTC | 36 |

| # | 4-splint library (*CCL-4*) (*SRC-14nt*) | Length [nt] |
|---|---|---|
| 4a | TTAGTCANTGTCCCGATTAGAGCCTTATGAGCCGTC | 36 |
| 4b | GGGACANTGACTAAGATTAGAGCCTTATGAGCCGTC | 36 |

| # | 16-splint library (*CCL-16*) | Length [nt] |
|---|---|---|
| 5a | TTAGTNAGTNTCCCGATTAGAGCCTTATGAGCCGTC | 36 |
| 5b | GGGANACTNACTAAGATTAGAGCCTTATGAGCCGTC | 36 |

| # | 64-splint library (*CCL-64*) | Length [nt] |
|---|---|---|
| 6a | TTAGTNANTNTCCCGATTAGAGCCTTATGAGCCGTC | 36 |
| 6b | GGGANANTNACTAAGATTAGAGCCTTATGAGCCGTC | 36 |

| # | 256-splint library (*CCL-256*) | Length [nt] |
|---|---|---|
| 7a | TTAGTNANTNTNCCGATTAGAGCCTTATGAGCCGTC | 36 |
| 7b | GGNANANTNACTAAGATTAGAGCCTTATGAGCCGTC | 36 |

| # | 18nt 4-splint library (*SRC-18nt*) | Length [nt] |
|---|---|---|
| 8a | GTTTAGTCANTGTCCCGTGATTAGAGCCTTATGAGCCGTC | 40 |
| 8b | ACGGGACANTGACTAAACGATTAGAGCCTTATGAGCCGTC | 40 |

| # | 12nt 4-splint library (*SRC-12nt*) | Length [nt] |
|---|---|---|
| 9a | TAGTCANTGTCCGATTAGAGCCTTATGAGCCGTC | 34 |
| 9b | GGACANTGACTAGATTAGAGCCTTATGAGCCGTC | 34 |

| # | 10nt 4-splint library (*SRC-10nt*) | Length [nt] |
|---|---|---|
| 10a | AGTCANTGTCGATTAGAGCCTTATGAGCCGTC | 32 |
| 10b | GACANTGACTGATTAGAGCCTTATGAGCCGTC | 32 |

| # | 8nt 4-splint library (*SRC-8nt*) | Length [nt] |
|---|---|---|
| 11a | GTCANTGTGATTAGAGCCTTATGAGCCGTC | 30 |
| 11b | ACANTGACGATTAGAGCCTTATGAGCCGTC | 30 |

| # | 6nt 4-splint library (*SRC-6nt*) | Length [nt] |
|---|---|---|
| 12a | TCANTGGATTAGAGCCTTATGAGCCGTC | 28 |
| 12b | CANTGAGATTAGAGCCTTATGAGCCGTC | 28 |

| # | FRET probe | Length [nt] |
|---|---|---|
| 13a | /Cy5/CCGAGGACTGAGGGTTTTAGGAGTTGGTCTATAATCATGG | 40 |
| 13b | AGACCAACTCCTAAAACCCTCAGTCCTCGG/IAbRQSp/ | 30 |

| # | Heat-activated blocking strands (converts *CCL-4* into a *HAC*) | Length [nt] |
|---|---|---|
| 14a | AAGACANTGACTGT | 14 |
| 14b | ACAGTCANTGTCTT | 14 |

| # | Heat-activated blocking strands (converts *CCL-64* into a *HAC*) | Length [nt] |
|---|---|---|
| 15a | AAGANANTNACTGT | 14 |
| 15b | ACAGTNANTNTCTT | 14 |

| # | 6-FAM 5'-end modified strand | Length [nt] |
|---|---|---|
| 16 | /56-FAM/AAGAGTACAGTCCAGATTAGAGCCTTATGAGCCGTC | 36 |

| # | Directly-grafted 64-splint library (*CCL-64*) | Length [nt] |
|---|---|---|
| 17a | /5Acryd/TTAGTNANTNTCCC | 14 |
| 17b | /5Acryd/GGGANANTNACTAA | 14 |

**Table S 2.** AF4-LS characterization of DyNAtrix backbones.

| | $M_w$ [*1] (kg/mol) | $M_n$ [*1] (kg/mol) | Đ ($M_w/M_n$) | $R_g$ (nm) | $R_h$ (nm) | $R_g/R_h$ | $\rho_{app}$ [*2] (g/l) | $V_h$ [*3] (µm³) |
|---|---|---|---|---|---|---|---|---|
| $\mathbf{P}_1$ | 2,820 | 2,220 | 1.27 | 94 | 89 | 1.06 | 1.27 | 0.0029 |
| $\mathbf{P}_5$ | 3,343 | 2,907 | 1.15 | 96 | 99 | 0.97 | 1.29 | 0.0041 |
| $\mathbf{P}_5^{\mathrm{RGD}}$ | 3,450 | 2,530 | 1.43 | 96 | 71 | 1.38 | 3.38 | 0.0016 |
| $\mathbf{P}_{10}$ | 2,280 | 2,140 | 1.06 | 91 | 60 | 1.53 | 3.99 | 0.0009 |
| $\mathbf{P}_{10}^{\mathrm{RGD}}$ | 2,560 | 2,440 | 1.05 | 87 | 55 | 1.58 | 5.94 | 0.0007 |

$M_W$ = mass average molecular weight; $M_n$ = number average molecular weight; Đ = polydispersity index; $R_g$ = radius of gyration; $R_h$ = hydrodynamic radius; $\rho = \rho_{app} =$ apparent density; $V_h$ = apparent volume

[*1] dn/dc = 0.170 mL/g; [*2] calculated from $M_n$ and $R_h$; [*3] calculated from $R_h$

**Table S 3.** Batch-to-batch *variation: comparison of the mass average polymer molecular weight, $M_W$, of three different synthesis products of $\boldsymbol{P}_{37}$ and three different synthesis products of $\boldsymbol{P}_{37}^{RGD}$.*

| | $M_w$ (kg/mol) | |
|---|---|---|
| | $\mathbf{P}_5$ | $\mathbf{P}_5^{\mathrm{RGD}}$ |
| *Synthesis 1* | 3,360 | 2,960 |
| *Synthesis 2* | 3,950 | 3,680 |
| *Synthesis 3* | 2,720 | 3,710 |
| **Average** | **3,343** | **3,450** |
| ± Std. Dev. | 615 | 347 |

**Table S 4.** Cost calculation of DyNAtrix.

| **Materials/ Reagents** | **Unit** | **Unit cost** | **Cost per mL of 1% gel** |
|---|---|---|---|
| Anchor strand | 1 nmol | € 0.06 | € 5.72 |
| Fmoc-L-Lys(Acryloyl)-OH | 1 mg | € 0.17 | € 1.38 |
| DNA Crosslinkers | 1 nmol | € 0.08 | € 6.00 |
| Heat-activated blocking strands | 1 nmol | € 0.08 | € 12.00 |
| **DyNAtrix [+RGD]** | | | **€ 25.10** |
| Actin (optional) | 1 µg | € 0.10 | € 39.64 |
| **DyNAtrix [+RGD] [+ actin]** | | | **€ 64.74** |
