## Supplementary figures and images for "Dynamic matrices with DNA-encoded viscoelasticity for advanced cell and organoid culture"

### Supplementary File 1

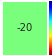

### Supplementary File 2

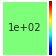

### Supplementary File 3

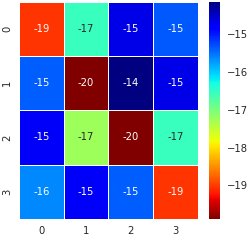

### Supplementary File 4

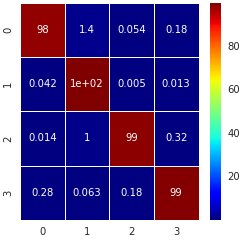

### Supplementary File 5

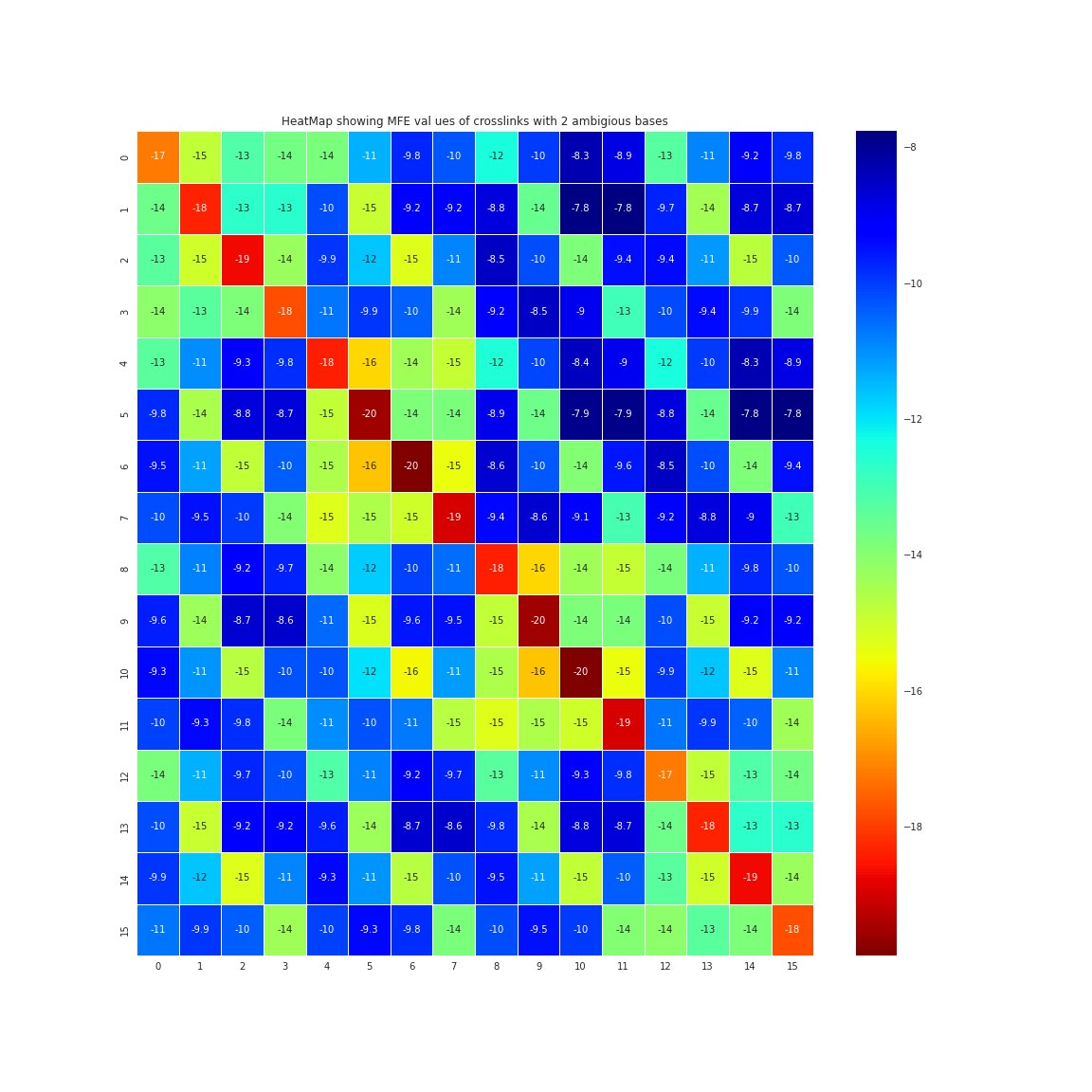

### Supplementary File 6

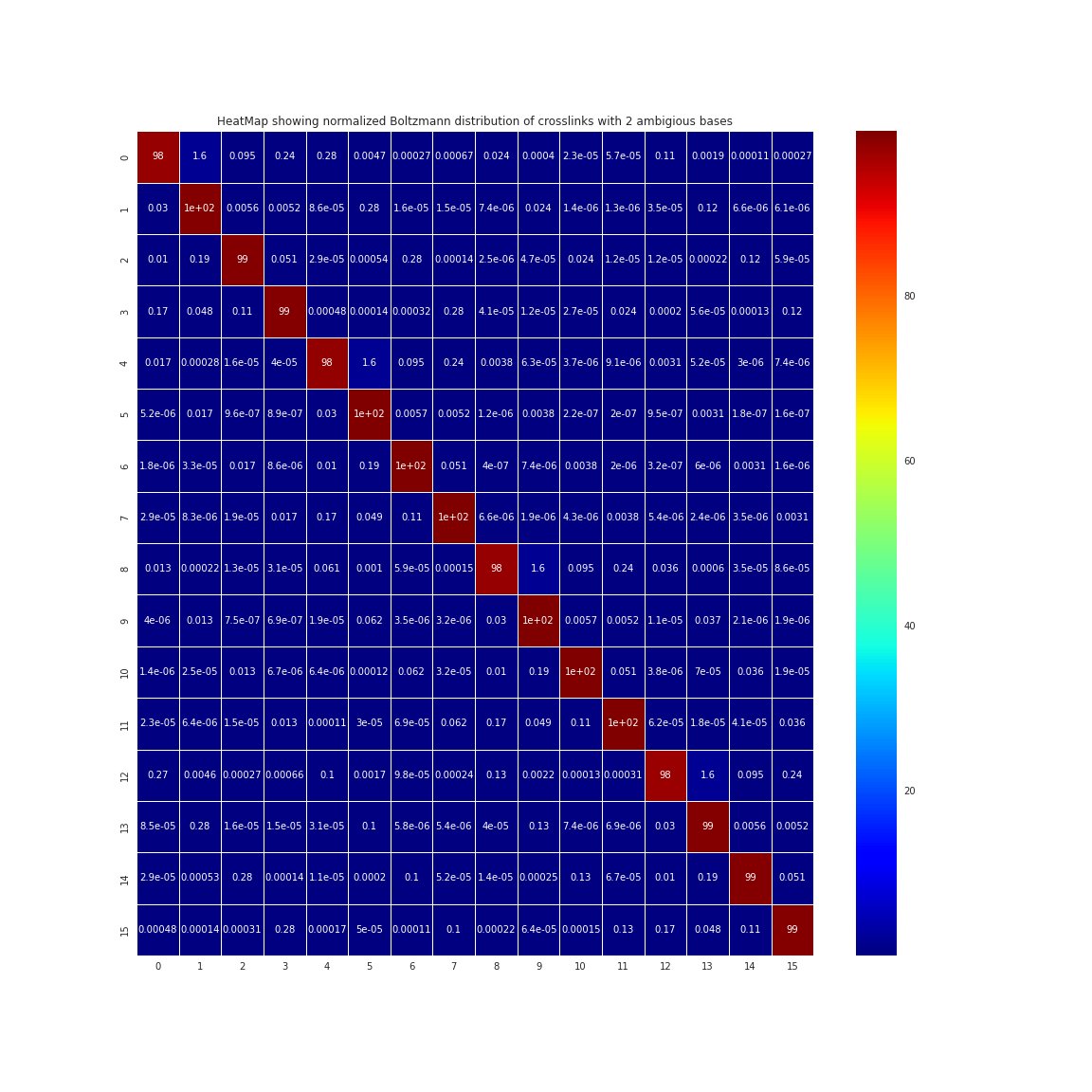

### Supplementary File 7

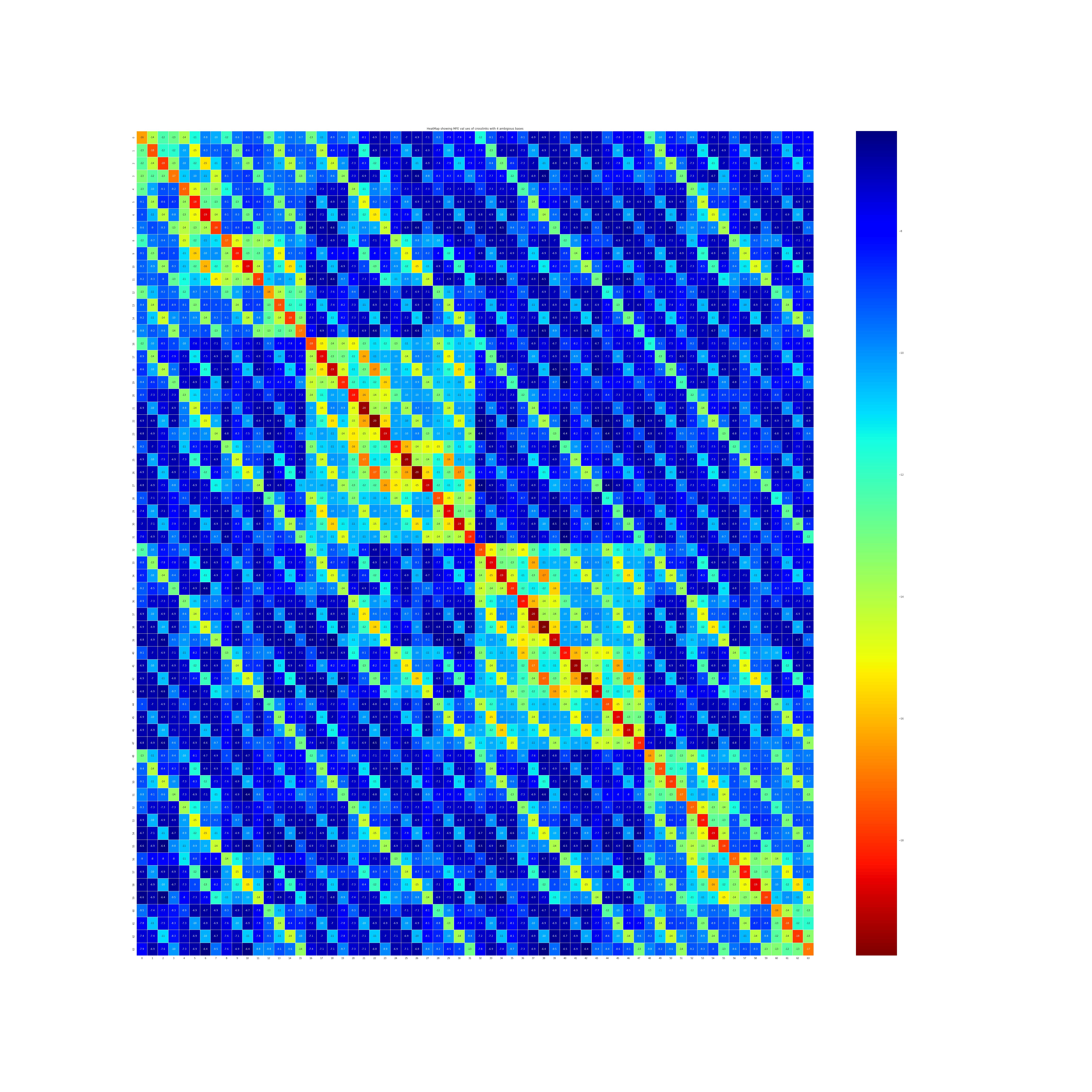

### Supplementary File 8

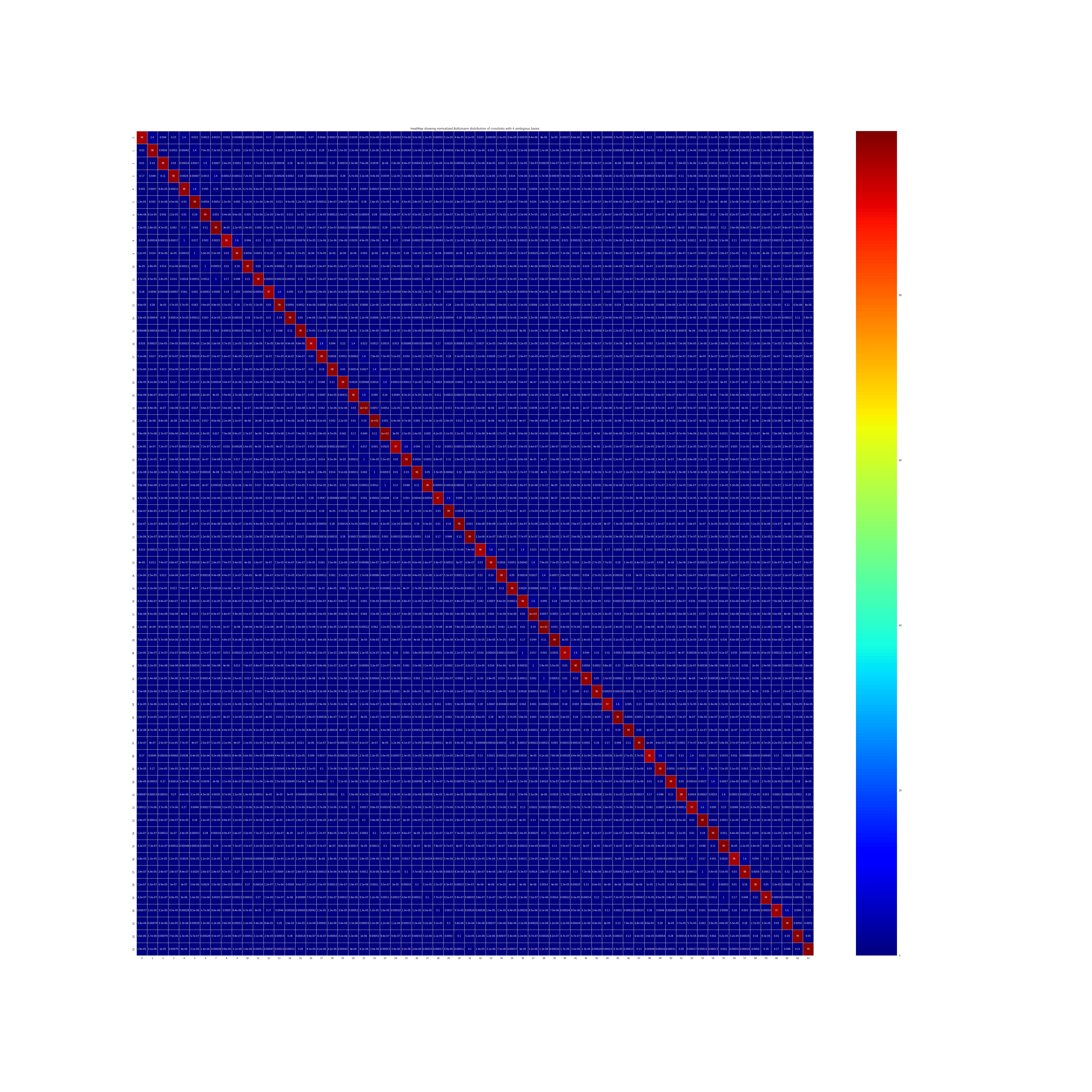

### Supplementary File 9

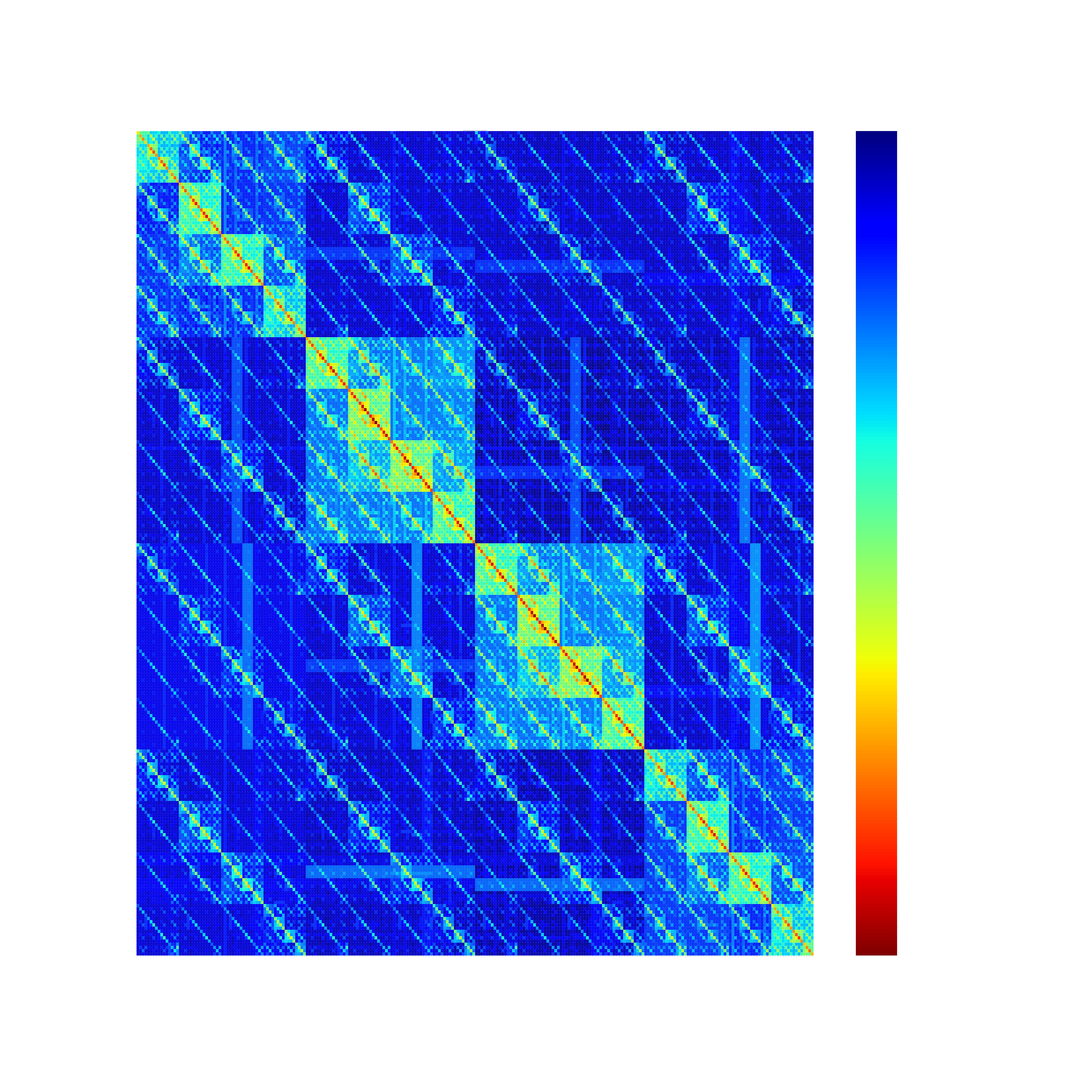

### Supplementary File 10

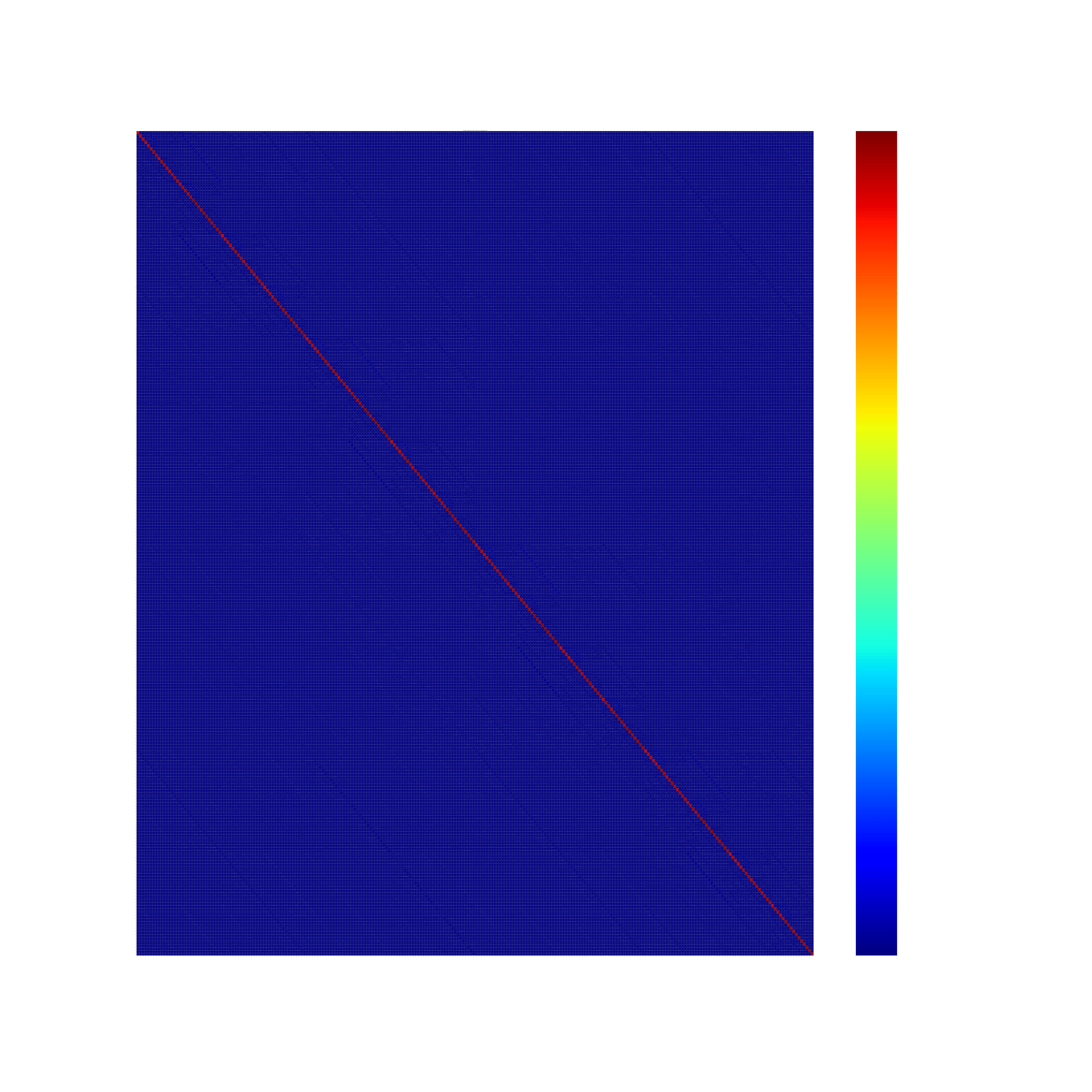

### Supplementary Video 1

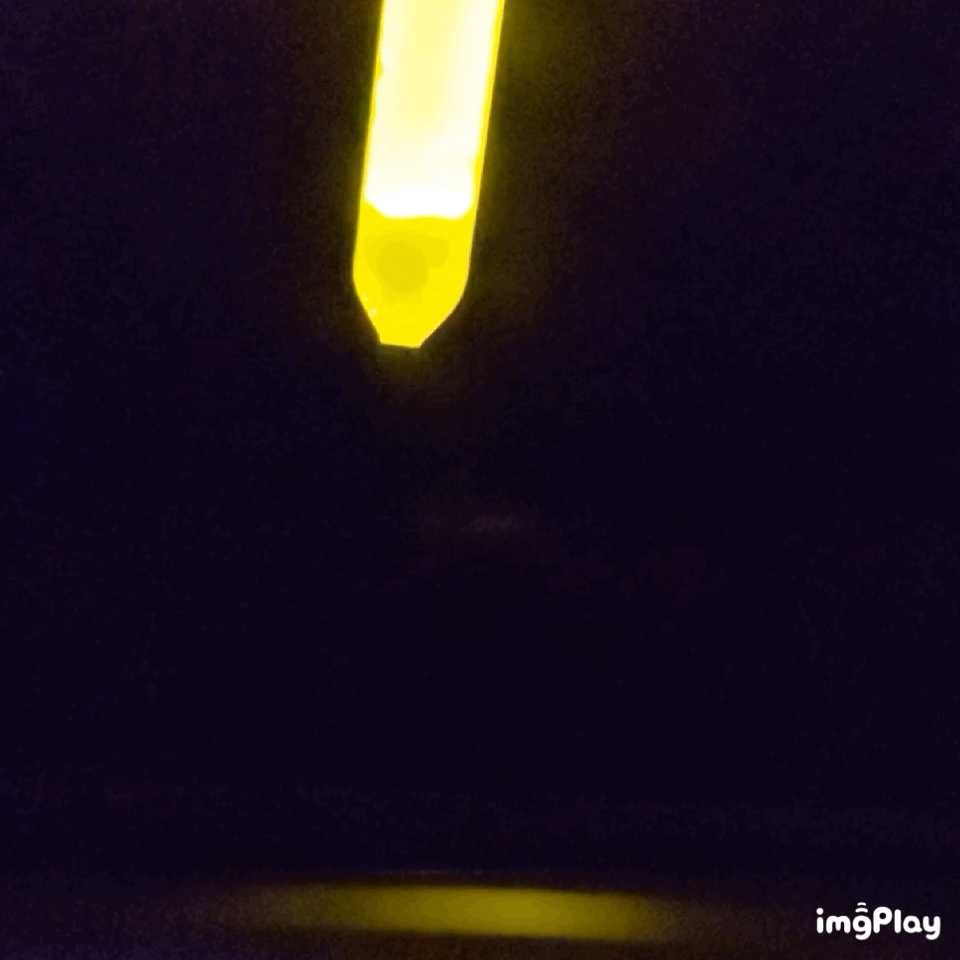
